# Domestication in dry-cured meat *Penicillium* fungi: convergent specific phenotypes and horizontal gene transfers without strong genetic subdivision

**DOI:** 10.1101/2022.03.25.485132

**Authors:** Ying-Chu Lo, Jade Bruxaux, Ricardo C. Rodríguez de la Vega, Samuel O’Donnell, Alodie Snirc, Monika Coton, Mélanie Le Piver, Stéphanie Le Prieur, Daniel Roueyre, Joëlle Dupont, Jos Houbraken, Robert Debuchy, Jeanne Ropars, Tatiana Giraud, Antoine Branca

## Abstract

Some fungi have been domesticated for food production, with genetic differentiation between populations from food and wild environments, and food populations often acquiring beneficial traits through horizontal gene transfers (HGTs). Studying their adaptation to human-made substrates are of fundamental and applied importance, for understanding adaptation processes and for further strain improvement. We studied here the population structures and phenotypes of two distantly related *Penicillium* species used for dry-cured meat production, *P. nalgiovense*, the most common species in the dry-cured meat food industry, and *P. salamii*, used locally by farms. Both species displayed low genetic diversity, lacking differentiation between strains isolated from dry-cured meat and those from other environments. Nevertheless, the strains collected from dry-cured meat within each species displayed slower proteolysis and lipolysis than their wild conspecifics, and those of *P. nalgiovense* were whiter. Phenotypically, the non-dry-cured meat strains were more similar to their sister species than to their conspecific dry-cured meat strains, indicating an evolution of specific phenotypes in dry-cured meat strains. A comparison of available *Penicillium* genomes from various environments revealed HGTs, particularly between *P. nalgiovense* and *P. salamii* (representing almost 1.5 Mb of cumulative length). HGTs additionally involved *P. biforme*, also found in dry-cured meat products. We further detected positive selection based on amino-acid changes. Our findings suggest that selection by humans has shaped the *P. salamii* and *P. nalgiovense* populations used for dry-cured meat production, which constitutes domestication. Several genetic and phenotypic changes were similar in *P. salamii*, *P. nalgiovense,* and *P. biforme*, indicating convergent adaptation to the same human-made environment. Our findings have implications for fundamental knowledge on adaptation and for the food industry: the discovery of different phenotypes and of two mating types paves the way for strain improvement by conventional breeding, to elucidate the genomic bases of beneficial phenotypes and to generate diversity.

## Introduction

The way in which populations adapt to their environment is a fundamental issue in biology. Domestication is widely used as a model for understanding adaptation as it corresponds to a recent adaptation process caused by strong, human-driven selection (Darwin, 1859, 1868). Humans have grown and bred various organisms, choosing individuals with the most interesting features for their particular uses or for consumption. For instance, many morphological traits differ between domesticated maize and its wild ancestor, including floral morphology, seed size, dispersal ability, stem number and length (Hufford et al., 2012; Stitzer & Ross-Ibarra, 2018). Charles Darwin used domesticated pigeons as a model for investigating adaptation and natural selection processes (Darwin, 1859, 1868). The parallel adaptation of distantly related species to the same environment provides an excellent study model for investigating whether evolution is repeatable — i.e., whether independent adaptation events to the same ecological niche involve similar phenotypic and genomic changes — a situation known as convergence (Chan et al., 2010; Colosimo et al., 2004; Duboué et al., 2011). In addition to adaptive changes, domesticated populations may suffer from bottlenecks, in which a drastic loss of genetic diversity occurs due to the strong selection exerted by humans. Such bottlenecks may jeopardize the ability of the domesticated populations to adapt (Maruyama & Fuerst, 1985; Nei et al., 1975). Studying domesticated organisms also has applied importance, for understanding the genomic bases of desired traits, for further variety generation and for maintaining genetic diversity that could allow future improvement.

Fungi are good models for studies of adaptation using domestication. Their experimental tractability makes it possible to perform fitness tests in controlled environments, and their small and compact genomes facilitate genomic studies to elucidate the genetic basis of adaptation (Gibbons et al., 2012; Gladieux et al., 2014; Legras et al., 2018; Ropars et al., 2015; Watarai et al., 2019; Wolfe & Dutton, 2015). Many fungi have been domesticated for food production (Dupont et al., 2017; Steensels et al., 2019), the chief examples being *Saccharomyces, Aspergillus* and *Penicillium*. *Saccharomyces cerevisiae* is the most studied domesticated fungal species, with different populations used for winemaking, baking and brewing (Almeida et al., 2014; Gallone et al., 2016; C. Gonçalves et al., 2018; Legras et al., 2018; Peter et al., 2018). Fungi are also used for cheese ripening. For example, *P. roqueforti* is used for blue-veined cheeses and *P. camemberti* is used for soft cheeses, such as Brie and Camembert. Both fungi have recently emerged as useful models for adaptation and domestication studies (Ropars et al., 2015, 2020; Dumas et al., 2020; Cheeseman et al., 2014; Ekseth et al., 2014; Williams et al., 1986; Bodinaku Ina et al., 2019; Crequer et al., 2023). The *Saccharomyces cerevisiae, P. camemberti* and *P. roqueforti* populations used in the food industry have specific traits that are beneficial for food production and enable them to thrive in their human-made environments to a much greater extent than other populations (Dumas et al., 2020; Marsit & Dequin, 2015; Ropars et al., 2020), providing evidence of genuine domestication. The genomic basis of several of these adaptive traits has been determined in *Saccharomyces* and *Penicillium* food populations, with the identification of gene duplications, interspecific hybridizations and horizontal gene transfers (Almeida et al., 2014; Cheeseman et al., 2014; C. Gonçalves et al., 2018; Guillamón & Barrio, 2017; Marsit et al., 2015, 2017; Ropars et al., 2015). Several adaptive horizontal gene transfers have occurred between distantly related cheese *Penicillium* fungi, leading to rapid convergent adaptation through the very same genes and genomic mechanisms (Cheeseman et al., 2014; Ropars et al., 2015). As expected from a domestication process in selfers or asexuals, the fungal populations used for food production have lost much of their genetic variability, as demonstrated for *S. cerevisiae* from wine and beer and *Penicillium* fungi from cheese (Dumas et al., 2020; Legras et al., 2018; Ropars et al., 2020). Most fungi can be cultured asexually, leading to a particularly large loss of genetic variability in domesticated fungi (Dumas et al., 2020; Ropars et al., 2020).

Like cheese fungi, *Penicillium nalgiovense* is an excellent model for studying domestication and adaptation. This species has been used to inoculate the surface of most dry-cured meat products for decades, to enhance the ripening process (M. Coton et al., 2021; Garcia et al., 2001; Ludemann et al., 2004; Sunesen & Stahnke, 2003). Another recently described fungus, *Penicillium salamii*, also occurs in dry-cured meats (Perrone et al., 2015) and may be included in starter cultures or form part of the natural mycobiota at local production sites (M. Coton et al., 2021; Magistà et al., 2016). Several traits have probably been subject to similar selection processes in both these fungi, either deliberately through human intervention or unwittingly, to enable these fungi to thrive in dry-cured meats and produce high quality products, with an appealing appearance and aroma, and a better shelf life and product safety. The fungi colonize the surface, thereby helping to preserve the dry-cured meat and to protect it against excessive desiccation, irradiation by light, oxygen and undesirable microorganisms, such as contaminating toxigenic molds (Chávez et al., 2011; Nielsen et al., 1998). These fungal species also help to improve the aroma of the dry-cured meat, through the metabolites they produce with their lipase, protease, transaminase and dehydrogenase activities (Leroy et al., 2006; Sunesen & Stahnke, 2003). However, if proteolytic and lipolytic activities are too high, this can result in an undesirable texture, bitterness and a rancid taste (Toldrá, 1998). Another important trait is the color of *Penicillium* fungi, which is also under selection, to meet human preferences (Filtenborg et al., 1996) or because the protection that pigments provide in nature, e.g., against desiccation, radiation and various stresses, may not be needed in food products anymore (Lin & Xu, 2020). Salt is added to dry-cured meats during production as a taste enhancer; it also helps to exclude undesirable microorganisms by reducing water activity, one of the key mechanisms of preservation in these products. The *Penicillium* fungi used to inoculate dry-cured meats may, therefore, have evolved a high degree of salt tolerance. *Penicillium nalgiovense* and *P*. *salamii* have also been isolated from environments other than dry-cured meats, such as the soil, plants, sewage and dog bones (Visagie et al., 2014). However, it remains unknown whether the dry-cured meat *Penicillium* fungi display particular phenotypes and/or form populations that are genetically isolated from those found in other environments. *Penicillium nalgiovense* and *P*. *salamii* lineages have diverged at least 30 million years ago (Houbraken et al., 2020; Perrone et al., 2015; Steenwyk et al., 2019), which corresponds, approximately, to the divergence between humans and Cercopithecoids (macaques, langurs…) (Dos Reis et al., 2018; Pozzi et al., 2014); a convergent adaptation to dry-cured meat, if observed, would therefore constitute an interesting case of parallel adaptation.

The aim of this study was to determine whether, within these two *Penicillium* species, the strains used in dry-cured meat production had features typical of domesticated organisms, by investigating whether dry-cured meat strains 1) were genetically different from the strains isolated from other environments, 2) displayed a genome-wide reduction of genetic diversity potentially indicative of strong population reduction and/or clonality, 3) had specific traits that would make them better suited to human needs or preferences than other strains, 4) displayed footprints of genomic adaptation, and 5) displayed convergent phenotypic and/or genomic changes. We addressed these questions by isolating strains from dry-cured meat, sequencing the genomes of multiple *P*. *nalgiovense* and *P*. *salamii* strains collected from dry-cured meat and other environments, and searching for genetic differentiation, recombination footprints, horizontal gene transfers, specific genome arrangements and traces of positive selection in terms of high rates of non-synonymous changes. Phenotypic differences between *Penicillium* fungi from dry-cured meat environments and other species/populations were also investigated by comparing their growth rates at different salt concentrations, color, lipid and protein degradation rates and their ability to produce harmful toxins. For interpretation of the phenotypic differences between strains from dry-cured meat and other environments in terms of domestication, we further compared the phenotypes of the non-dry-cured meat strains to those of their respective sister species (*P. chrysogenum* for *P. nalgiovense* and *P. olsonii* for *P. salamii*). Under the hypothesis that dry-cured meat populations are domesticated, we would expect traits essential for dry-cured meat products to have evolved specifically in dry-cured meat strains, and to be absent in strains from other environments and in the corresponding sister species. In addition, because we found horizontally transferred genomic regions shared between *P*. *nalgiovense, P*. *salamii* and *P. biforme* and because *P. biforme* is also used to inoculate dry-cured meat, albeit rarely, we also compared phenotypes between strains from dry-cured meat and other environments for *P. biforme* and its sister species, *P. palitans*. We cannot entirely rule out that feral strains could be present among fungi isolated from non-dry-cured meat environments (i.e., escaped from dry-cured meat). In that case, however, they could only reduce the differences between dry-cured meat and non-dry-cured meat groups. Hence, any differences found between dry-cured meat and non-dry-cured meat groups are conservative regarding the occurrence of feral strains. Furthermore, we tried to identify feral strains by looking for strains isolated from non-dry-cured meat environments but resembling phenotypically and/or genetically dry-cured meat strains.

## Material and Methods

### Strain collection and DNA extraction

We used 94 strains for experiments and genome analyses including 22 isolated from dry-cured meat samples in this study (see below; Table S1), 33 from the CBS collection housed at the Westerdijk Fungal Biodiversity Institute (Utrecht, the Netherlands), 33 from the public collection of the national museum of natural history in Paris (LCP) and six from previous studies. Strain isolation, DNA extraction and species identification protocols are described in the Supplementary methods.

### Genome sequencing and genome assembly

For short-read sequencing, DNA libraries and sequencing were performed by the INRAE GenoToul platform (Toulouse, France). Paired-end libraries of 2x150 bp fragments were prepared with TruSeq Nano DNA Library Prep Kits (Illumina), and sequencing was performed on a HiSeq3000 Illumina sequencer, with a mean final coverage of 38x and a median of 10x (genomes of 27 to 35 Mb). We sequenced the genomes of 55 strains collected from either dry-cured meat or other environments (Table S1).

We used the P6/C4 Pacific Biosciences SMRT technology to obtain a high-quality reference genome sequence of the LCP06525 *P. salamii* strain and the ESE00252 *P. nalgiovense* strain, both from dry-cured meat. Nuclear genomic DNA was extracted with the cetyltrimethylammonium bromide (CTAB) extraction procedure described previously (Cheeseman et al., 2014). Sequencing was performed by the UCSD IGM Genomics Facility La Jolla, CA, USA. Long- and short-read assemblies are described in the Supplementary methods. All the genomes were deposited on ENA under the study accession number PRJEB44534 (Table S1).

### Population genetic analyses

We mapped cleaned sequencing reads to the reference genomes LCP06525 (*P. salamii*) and LCP06099 (FM193; *P. nalgiovense*; accession numbers HG815136–HG815288 and HG815290–HG816004; Ropars et al., 2015) using Stampy with default parameters (v1.0.32; Lunter & Goodson, 2011). SNPs were called for each individual using the genome analysis toolkit GATK (v4.1.0.0; McKenna et al., 2010) with the *RealignerTargetCreator*, *IndelRealigner* and *UnifiedGenotyper* functions using default parameters except that we used a ploidy of one. For each species, we merged the variant call formats (VCF) files using the *CombineVariants* function in GATK and tagged the sites as follows: quality (with low qual-tag), quality by depth (QD < 2.0), mapping quality (MQ < 40), Fisher strand bias (FS > 60; a statistic to detect strand bias in variant calling in reverse and forward strands), mapping quality rank sum (MQRankSum < 12.5; to test read quality differences between reference and alternative alleles), read position rank (ReadPosRank < 8.0; based on the distance from the end of reads compared with alternative allele) and missing data (allowing maximum 10% missing data). The filter-tags were prepared using vcftools with PASS/filtered tag and GATK with the *VariantFiltration* function. Indels and tagged SNPs were entirely removed from the original VCF files using vcftools (v0.1.16; Danecek et al., 2011). SNPs were used to analyze population structure and linkage disequilibrium (LD). To produce an alignment for a phylogenetic analysis, we had to include the *P. nalgiovense* PacBio sequence (ESE00252) and the respective outgroups which were available as fasta files and therefore absent in the vcf file. We thus used NUCmer v3.1 from the MUMmer3 package (Kurtz et al., 2004) to align those sequences against the two reference genomes (LCP06099 for *P. nalgiovense* and LCP06525 for *P. salamii*) and called SNPs. We then merged those SNPs with the previous vcf files in R v3.6.2 (R Core Team, 2019). We built population trees using RAxML (v8.2.7; Stamatakis, 2014), run with the general time-reversible model of nucleotide substitution with the gamma model of rate heterogeneity (GTRGAMMA) and 100 bootstraps.

In order to analyze population subdivision, we filtered our SNP dataset with bcftools (v1.7; Li, 2011) to keep only distant SNPs to avoid too strong LD (r^2^ lower than 0.2 in a window of 1kbp), with a minor allele present in at least two individuals per species (minor allele frequency of 0.05). We then ran STRUCTURE (v2.3.4; Pritchard et al., 2000) with the admixture model during 20,000 iterations and removed the first 10,000 iterations as burn-in, for a number of populations (K) varying from 2 to 10, with 10 independent runs for each K. We combined the results with the R package pophelper v2.3.0 (Francis, 2017). We used the full fasta alignment built to estimate the population tree to run a principal component analysis following the code used in Konishi et al. (2019) and ran it in R v3.6.2 (R Core Team, 2019). We performed neighbor-net analyses with the *phangorn* R package (Schliep, 2011). The substitution model used for building the distance matrix was JC69 (Jukes & Cantor, 1969; Li, 2011). The two most divergent *P. nalgiovense* strains, CBS297.97 and DTO 204-F1, were removed from the neighbor-net analyses to better see the remaining relationships.

We searched for the mating-type locus in the genomes and identified the alleles as MAT1-1 or MAT1-2 with tblastx v2.6.0 and the *P. chrysogenum* strain ATCC02889 MAT1-1 (AM904544.1) and strain NRRL1249B21 MAT1-2 (AM904545.1) sequences as references. The presence of the two different mating types in a population could potentially allow sexual reproduction and recombination (Böhm et al., 2015; Ropars et al., 2014), while the presence of a single mating type suggests strict asexual reproduction. Fungi able to undergo sexual reproduction between strains carrying the same mating-type idiomorph in their haploid genome are indeed extremely rare and this phenomenon is not known in any *Penicillium* fungus (Wilson et al., 2021).

### Detection of horizontal gene transfers between *P. salamii* and *P. nalgiovense*

We aimed at detecting very recent horizontal gene transfers (HGTs) that could be involved in parallel adaptation to the dry-cured meat environment, having therefore occurred in both *P. salamii* and *P. nalgiovense* during domestication and thus human times. Genomic regions resulting from such recent HGTs were expected to be nearly identical between the two distant species *P. salamii* and *P. nalgiovense* in contrast to the genomic background, and lacking in many non-dry-cured meat strains and/or species (e.g. Cheeseman et al., 2014). By the very nature of such recent HGTs, the resulting genomic regions (HTRs for horizontally transferred regions) are lacking in many strains and are almost identical in the strains in which they are present, which prevents using phylogenetic incongruence between different regions in the genomes to detect or validate HGTs, but provides stronger evidence. Other HTRs, present only in one of these species, may be involved in adaptation to dry-cured meat. However, we focused on the shared ones, because we were particularly interested in parallel adaptation.

In order to identify HTRs, we first searched for thresholds above which we could consider that regions were more similar across a longer length than expected given the species phylogenetic distance. Given that the phylogenetic distance between *P. olsonii* and *P. chrysogenum* or *P. rubens* (two species formerly considered as one) is the same as the distance between *P. nalgiovense* and *P. salamii*, sequences that would be more similar and longer than the ones typically detected between *P. olsonii* and *P. chrysogenum* or *P. rubens* would not be expected given the phylogenetic distance separating *P. nalgiovense* from *P. salamii,* and would thus more likely correspond to recent gene flow, i.e. HGT or introgression. More details are given on the detection of horizontal gene transfers in Supplementary Methods, as well as on transposable element detection and filtering, and on gene function enrichment.

For plotting similarity levels along genomes, we merged several “reference” HTRs based on overlapping sequences, and we kept only those larger than 35 kbp, collectively adding to 1.1 Mbp. Global alignments between these long HTRs and each of the dry-cured meat *Penicillum* genome assemblies were obtained with minimap2 v2.22-r1105-dirty (Li, 2018) using the -cx asm20 (default settings for whole genome alignment up to 20% sequence divergence) and -- cs=long options. Plotting was performed based on the Pairwise mApping Format (*.paf) files with plot_coverage function of pafr (https://dwinter.github.io/pafr/index.html) for contig tracks and Rideogram (Hao et al., 2020) for synteny links in R v3.6.2 (R Core Team, 2019). The percentage of identity was calculated as the number of matching bases divided by the length of the query.

### Presence and similarity of HTRs in other species

We searched for the presence of the candidate HTRs in other *Penicillium* (n=47), *Aspergillus* (n=4) and *Monascus* (n=1) species by globally aligning the consensus sequences of the eight largest HTRs with genome assemblies (Table S2) using minimap2 (-cs asm20 and --cs=long options). We kept matches that either: 1) covered the entire HTR; 2) covered an entire contig on the assembly with at least 1 kbp of matched bases, or 3) that were larger than 10 kbp and found at the edges of larger contigs, allowing to a maximum of two such matches per HTR.

We calculated the fraction of each HTR present in the genomes as the sum of non-ambiguous matches length divided by the total length of the focal HTR, and the percentage of identity as the number of matched bases to the HTR consensus sequences divided by the sum of non-ambiguous matches length.

To further support the horizontal transfer inference, we compared the topologies of the gene genealogies to that of the species tree. For this goal, we extracted the genes present in the putative HTRs in each of the 54 *Penicillium*, *Aspergillus* and *Monascus* species (including the reference *P. salamii* and *P. nalgiovense* genomes) and used OrthoDB v. 11 (Kuznetsov et al., 2023) to obtain the publicly available orthologous sequences at the Eurotiales level. For each orthologous group, we removed the sequences that were duplicated within at least a strain to avoid paralogy issues. We kept only genes with at least five different species. We then aligned sequences in each group using mafft (Katoh et al., 2002) v7.310 and built gene trees using iqtree2 v2.0.3., an automatic detection of the best evolutionary mutation model, and 10,000 ultra-fast bootstraps (Nguyen et al., 2015) We compared each gene tree with the pruned Astral species tree (keeping only the same species as in the gene tree), using the approximately unbiased (AU) test (Shimodaira, 2002) implemented in iqtree2 with 10,000 replicates to assess whether the gene tree topology differed significantly from the species tree topology.

### Detection of selection: SnIPRE analyses and Tajima’s D

We calculated Tajima’s D (Tajima, 1989) using the *strataG* R package (v2.0.2; Archer et al., 2017) and tested genome-wide neutral evolution or recent change in demography by comparing the observed values to the 95% confidence interval around zero. A negative Tajima’s D indicates recent selective sweeps, weak negative selection or recent population expansion while a positive Tajima’s D indicates balancing selection, population structure or recent population contraction. We obtained nucleotide diversity indexes (π, the average number of pairwise nucleotide differences per site (Tajima, 1989); and θ_W_, the observed nucleotide diversity calculated from the number of segregating sites (Watterson, 1975)) using the strataG (v2.0.2; Archer et al., 2017) and pegas (v0.14; Paradis, 2010) packages and scaling to the length of the reference genome. We used vcftools to measure LD, in windows of 15,000 bp and with a minor allele frequency cutoff of 0.2 (Danecek et al., 2011).

We also aimed at detecting individual genes evolving under positive selection in terms of high rates of non-synonymous substitutions. Because the two species studied here are used in industrial contexts (Garcia et al., 2001; Ludemann et al., 2004; Magistà et al., 2016; Sunesen & Stahnke, 2003), and therefore potentially cultured in high quantities after an initial bottleneck, we cannot assume a constant effective population size through time. Even more importantly, these fungi are cultivated asexually to be sold as starters in the industry, therefore constituting clonal lineages (see results). Most of the classic frameworks for detecting genes under positive selection are therefore unsuitable. We decided to use SnIPRE (Eilertson et al., 2012), a Bayesian generalization of the McDonald and Kreitman analyses (McDonald & Kreitman, 1991), which does not make any assumption on demography or reproduction mode. This method detects genes in which amino-acid changes are more or less frequent than expected under neutrality (inferring positive or purifying selection, respectively), by identifying genes in which non-synonymous divergence to an outgroup is higher or lower than expected based on observed synonymous and nonsynonymous SNPs in the in-group, thus accounting for gene-specific mutation rates. We ran the tests for *P. nalgiovense* using *P. chrysogenum* as an outgroup and for *P. salamii* using *P. olsonii* as an outgroup. For this goal, we applied the method described in supplementary for the phylogeny reconstruction to obtain single-copy genes present in all the *P. nalgiovense* strains and five *P. chrysogenum* on one dataset, and in all the *P. salamii* strains and five *P. olsonii* on a second dataset, allowing one missing sequence per orthologous gene alignment. From these alignments, we recovered the number of synonymous and non-synonymous sites, and among each category the number of fixed and polymorphic sites in each of *P. salamii* and *P. nalgiovense* gene with the MKtest program of libsequence (Thornton, 2003). We estimated the mean number of synonymous and polymorphic sites by performing a weighted average on the polydNdS output. The MKtest and polydNdS programs belong to the package “analysis” of libsequence (Thornton, 2003). We collected the results of these analyses and used them as input for the SnIPRE analysis. We ran it in its Bayesian implementation following the script providing by the authors for 25,000 iterations and removed a burn-in of 10,000 iterations, with a thinning value of four. To test whether the proportion of genes under positive, negative, or neutral selection was different between one specific COG (cluster of orthologous groups) functional category and the rest of the genome, we used Fisher’s exact tests as described in the supplementary for the HTRs. As strains from dry-cured meat did not cluster together neither in the phylogenetic trees nor in the population genomic analyses (see results), we did not compare these strains with others within each species.

### Phenotypic tests

We compared various phenotypic traits between species and strains from different environments: growth rate at different salt concentrations, spore production, lipolysis, and proteolysis abilities. We first studied *P. nalgiovense* and *P. salamii*, the most common species inoculated for dry-cured meat production, with their respective sister species *P. chrysogenum* and *P. olsonii*. However, we found shared horizontally transferred regions between these dry-cured meat species and the *P. biforme* complex, including the two clonal lineages *P. camemberti* var. *camemberti* and *P. camemberti* var. *caseifulvum* (Ropars et al., 2020), and these can also be, even if more rarely, inoculated in dry-cured meat. We therefore also analyzed phenotypes in *P. biforme*, that display genetic and phenotypic variability, in contrast to the cheese clonal *P. camemberti* varieties (Ropars et al., 2020), and compared them to phenotypes of their sister species *P. palitans.* For all the experiments below, we used all available *P. nalgiovense*, *P. salamii* and *P*. *biforme* strains that we could find in public collections, from non-food environments, and similar numbers of strains, chosen at random, from dry-cured meat and from their closely related outgroup species. The Table S1 gives the number and identity of strains used for each species, type of environment and experiment. The Supplementary methods provide details on phenotypic measures as well as on mycotoxin or penicillin production.

### Statistical analyses of the phenotypic measures

We analyzed several phenotypic traits to test our hypothesis that dry-cured meat fungi are domesticated: growth and spore production at different salt concentrations using malt extract agar medium, colony color on malt extract agar medium, and lipolytic and proteolytic activities. If dry-cured meat fungi have been domesticated, we expected that: 1) there would be significant differences between dry-cured meat and non-dry-cured meat strains within each species, and 2) there would be no significant differences between the non-dry-cured meat strains and the sister species not used for dry-cured meat production; this would indeed suggest that only the fungi used for dry-cured meat production have evolved specific phenotypes related to adaptations to the dry-cured meat substrate. Because we found that *P. nalgiovense* and *P. salamii* shared horizontally transferred regions with the *P. biforme* complex, which can also be inoculated in dry-cured meat, we included *P. biforme* in phenotypic assays. We therefore tested first whether there were significant phenotypic differences between dry-cured meat and non-dry-cured meat strains within each of the species used for dry-cured meat maturation, i.e., *P. nalgiovense, P. salamii* and *P. biforme.* Then, we tested whether there were significant differences between the non-dry-cured meat strains (excluding dry-cured meat strains from this analysis) and the sister species populations, not used for dry-cured meat production (i.e., *P. olsonii* for *P. salamii, P. chrysogenum* for *P. nalgiovense* and *P. palitans* for *P. biforme*). To be able to compare sister species, we included in the tests a “clade” effect, clades corresponding to sister species pairs, to control for the phylogenetic relatedness in the analysis.

For all analyses, fixed effects were added sequentially to test for their significance through likelihood ratio tests until reaching the full model with all possible fixed effects. We thereafter only introduce the full model. All post-hoc tests were conducted using the emmeans function of package emmeans (Russell et al., 2018) with Tukey correction for pairwise mean comparison. Post-hoc tests were run when we found a significant effect either between sister species or between populations from dry-cured meat and non-dry-cured meat.

For analyzing growth at different salt concentrations, we fitted two linear mixed models on the log-transformed colony area, keeping the day of measurement as a random variable and the salt concentration as fixed effects in each of them. In the first one, testing the differences between strains from dry-cured meat and other environments, we further included as fixed effects the species and its interaction with salt concentration, as well as the isolation environment (dry-cured meat or not) nested within species and its interaction with salt concentration. In the second one, where we tested the differences between non-dry-cured meat strains and the sister species, we included as fixed effects the clade and its interaction with salt concentration, as well as the species nested within the clade and its interactions with salt concentration.

For quantifying spore number after growth at various salt concentrations, as too few spores were recovered at 18% salt concentration, we only analyzed growth data for 0%, 2% and 10% salt concentration. As spores were only counted once (after the 17-day growth experiment), we did not include any random effect in the model. We therefore fitted two linear models using as a dependent variable the log-transformed number of spores divided by the colony area to normalize the effect of growth on sporulation.

For lipolysis and proteolysis analyses, we fitted two linear mixed models on the distance to the lysis zone using the day of measurement and the batch number as random effects. To test for differences between dry-cured meat and other environment strains, we used as fixed effects species and isolation environment (dry-cured meat or not) nested within species. To test for differences between non-dry-cured meat populations and the sister species, we included as fixed effects the clade and the species nested within the clade.

For colony whiteness, we fitted two linear mixed models using the same fixed effects as for the two models used for proteolysis and lipolysis tests. As three color measurements per strain were done, we added the measured circle location and strain ID as random effects. For colony color, because red and green values were strongly correlated (r=0.998), we only used red and blue components in RGB space to fit generalized linear mixed models with a binomial response. We considered as random effects the measured circle location and strain ID. Each model used the same fixed effects as those for proteolysis and lipolysis tests.

All linear mixed models were fitted using the lmer() function of *lme4* R package (Bates et al., 2015; R Core Team, 2019) and glmer() for the generalized linear mixed models. Simple linear models were fitted with lm() function (R Core Team, 2019). Likelihood ratio tests between models were performed using anova() function in R (R Core Team, 2019). For all tests, we first checked that assumptions were met, i.e., homoscedasticity, independence of residuals, and normal and identical distribution of residuals.

## Results

### Genetic diversity and population structure in *P. salamii* and *P. nalgiovense*

We collected 140 dry-cured meat casings, from which we isolated 134 monospore-derived strains. The most frequent *Penicillium* species isolated from dry-cured meat was *P*. *nalgiovense* (*n*=70) followed by *P*. *salamii* (*n*=16). We also found *P*. *biforme* (*n*=5), *P*. *chrysogenum* (*n*=5), *P*. *solitum* (*n*=5) and *P. camemberti* (*n*=5) and, sporadically, other *Penicillium* species and other genera (Table 1). In six cases (5% of the 106 samples with these six most abundant *Penicillium* species), we isolated two different species from the same sample, but never *P. nalgiovense* and *P. salamii* together (Fig. S1).

**Table 1.** Species identified among the 134 monospore-derived strains isolated from dry-cured meat casings.

| Species | Number of identified<br>monospores |
| --- | --- |
| **Total *Penicillium*** | 124 |
| *P. nalgiovense* | 70 |
| *P. salamii* | 16 |
| *P. biforme* | 5 |
| *P. camemberti* | 5 |
| *P. chrysogenum* | 5 |
| *P. solitum* | 5 |
| *P. polonicum* | 4 |
| *P. brevicompactum* | 2 |
| *P. chermesinum* | 2 |
| *P. crustosum* | 2 |
| *P. nordicum* | 2 |
| *P. expansum* | 1 |
| *P. glabrum* | 1 |
| *P. italicum* | 1 |
| *P. olsonii* | 1 |
| *P. roqueforti* | 1 |
| *P. tularense* | 1 |
| **Total other genera** | 10 |
| *Aspergillus* sp. | 2 |
| *Fusarium proliferatum* | 2 |
| *Scopulariopsis* sp. | 2 |
| *Alternaria* sp. | 1 |
| *Cladosporium salinae* | 1 |
| *Cladosporium*<br>*spinulosum* | 1 |
| *Geotrichum candidum* | 1 |

We obtained high-quality genome assemblies for *P. salamii* LCP06525 (37.00 Mbp) and *P. nalgiovense* ESE00252 (34.56 Mbp) with long-read (PacBio) sequencing technology and we sequenced the genomes of 55 strains with Illumina technology, including some of the strains we isolated as well as other available strains (Table S1). We identified 1,395,641 SNPs by mapping reads from the 21 Illumina-sequenced *P*. *salamii* genomes onto the *P. salamii* reference genome. A population tree based on 2,977,098 SNPs after the addition of the outgroup (Fig. 1A) revealed genetic differentiation by sampling environment in *P*. *salamii*. Two clades contained mostly dry-cured meat strains, whereas the other two clades contained mostly non-dry-cured meat strains. The frequencies of the two isolation environments differed significantly between the three clades with more than one strain (Chi²= 7.6723; df = 2; *p*-value = 0.02). No association with geographic origin was observed for *P*. *salamii* strains; the least diverse clade included only strains from Europe, but was also a clade containing mostly dry-cured meat strains (Fig. 1A).

**Figure 1.**
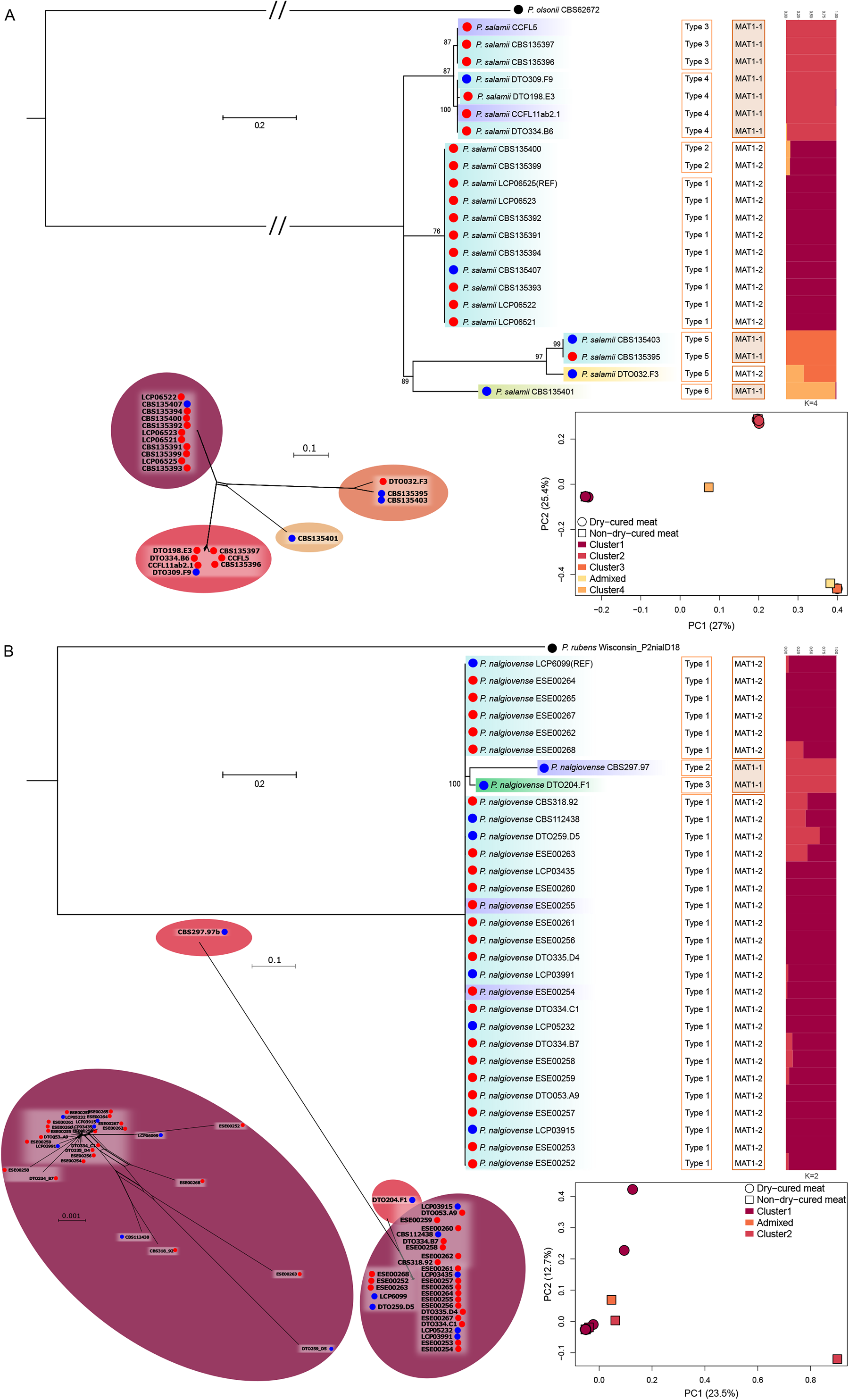
Rooted maximum likelihood population trees, principal component analysis (PCA), STRUCTURE and splitstree results based on genome-wide single nucleotide polymorphisms (SNPs) of *Penicillium salamii* (A) and *P*. *nalgiovense* (B), with information on genomic architecture types and mating-types. Bootstraps are indicated for the well-supported nodes (i.e. bootstraps higher than 70) and the scale indicates branch length (substitution rate per polymorphic site). The strain geographical origins are shown with colors (blue: Europe, purple: America, yellow: Australia, light green: Africa, green: Middle East). The color circles indicate environment of isolation, with dry-cured meat in red and other environments in blue. Architecture types correspond to groups of genomes with high collinearity. For the STRUCTURE plot, each bar represents an individual, with its estimated proportions of genetic information assigned to different population clusters, represented by colors, for K = 4 for *P. salamii* and K = 2 for *P. nalgiovense*. See Fig. S2 for barplots at other K values, giving similar patterns. In the PCA, the shape indicates the environment of isolation, and the color follows the STRUCTURE results. For each PCA axis, the percentage of variance explained is indicated between brackets. For the splitstrees, the bubbles colors follow the STRUCTURE results. For *P. nalgiovense*, the two most divergent strains CBS297.97 and DTO204.F1 were removed from the left splitstree. The *P. nalgiovense* tree has been rooted with *P. rubens* instead of *P. chrysogenum* because of the availability of a better genome assembly at the time of the analysis and of the very high genetic similarity between *P. rubens* and *P. chrysogenum*.

For the 28 *P*. *nalgiovense* genomes obtained by Illumina sequencing, reads were mapped onto the *P*. *nalgiovense* reference genome (LCP06099=FM193), leading to the identification of only 380,884 SNPs (381,996 when the other available PacBio genome ESE00252 was added), indicating a very low level of polymorphism. The *P*. *nalgiovense* population tree, based on a 2,367,990 SNPs with the outgroup (Fig. 1B), had very short branches for most strains, suggesting that these strains represent a clonal lineage. No genetic differentiation was observed, with no effect of sampling environment or geographic location in particular. Only two strains displayed some genetic differences from the main lineage.

We analyzed 1,766 distant SNPs for *P. nalgiovense* and 7,685 for *P. salamii*, to investigate the population structures of these two species. For *P. salamii*, the STRUCTURE analysis supported four well-delimited genetic clusters, consistent with the population tree (Fig. 1A, Fig. S2A). For *P. nalgiovense,* the STRUCTURE analysis was also consistent with the population tree, with two well-delimited genetic clusters (Fig. 1B, Fig. S2B); the other genetic clusters corresponded only to admixed strains, and were not, therefore, considered to correspond to genuine clusters. The PCA (Fig. 1) recovered patterns similar to those revealed by the population trees and STRUCTURE analyses: four clusters for *P. salamii*, with a tendency towards separation by geographic origin, and weaker structure for *P. nalgiovense*, with the most differentiated strain being from North America.

We identified six different genomic architecture types among *P. salamii* strains (Fig. 1A, Fig. S3). The genomes of *P. nalgiovense* strains were highly collinear, except for two strains displaying large rearrangements relative to the other strains (Fig. 1B). These collinearity groups in both *P. salamii* and *P. nalgiovense* corresponded to the genetic clusters detected above (Fig. 1). In *P. salamii*, genomic rearrangements differentiated the four strains obtained outside of Europe (USA and Middle East).

Nucleotide and genetic diversities were much higher in *P*. *salamii* (π = 0.0111 bp^-1^, SE = 1 x 10^-5^; θ_W_ = 0.0105 bp^-1^, SE = 7 x 10^-4^) than in *P*. *nalgiovense* (π = 0.0009 bp^-1^, SE = 1 x 10^-6^; θ_W_ = 0.0030 bp^-1^, SE = 2 x 10^-4^), which had diversity levels similar to those in *P. roqueforti* (π = 0.0011 bp^-1^; θ_W_ = 0.0007 bp^-1^; Dumas et al., 2020) and *P. biforme* (π = 0.0011 bp^-1^; θ_W_ = 0.0011 bp^-1^; Ropars et al., 2020). We detected both mating types (MAT1-1 and MAT1-2) in *P. salamii*, in balanced proportions and with a distribution across strains corresponding to the genetic structure and genomic architecture types (Fig. 1A). Each of the two MAT groups displayed relatively low nucleotide and genetic diversities (MAT1-1 group: π = 0.0007 bp^-1^, SE = 3 x 10^-6^; θ_W_ = 0.0006 bp^-1^, SE = 1 x 10^-4^; MAT1-2 group: π = 2 x 10^-5^ bp^-1^, SE = 4 x 10^-7^; θ_W_ = 3 x 10^-5^ bp^-1^, SE = 3 x 10^-6^). In *P. nalgiovense*, only the two most divergent strains carried MAT1-1, all the other strains being identified as MAT1-2, supporting the hypothesis of the use of a single clonal lineage for dry-cured meat production (Fig. 1B). The MAT1-2 group displayed very low diversity (π = 4 x 10^-5^ bp^-1^, SE = 5 x 10^-7^; θ_W_ = 8 x 10^-5^ bp^-1^, SE = 5 x 10^-6^).

Overall, *P. nalgiovense* displayed high levels of genome-wide LD, with no detectable decay over physical distance (Fig. S4). This finding is consistent with a complete absence of recombination due to exclusively asexual multiplication in this lineage. In *P. salamii,* we observed a sharp decay in LD within 200 bp, but LD level remained high. Analysis of LD within each of the two *P. salamii* populations with sufficient numbers of strains and both mating types revealed the same LD pattern as in *P. nalgiovense* (Fig. S4). This LD pattern in *P. salamii* is consistent with the asexual multiplication of two lineages of opposite mating types, with recombination events occurring in the ancestral population, before the asexual multiplication of these two lineages. The obtained neighbor-net networks further support the population structure identified and recent clonality in *P. salamii* and *P*. *nalgiovense* (Fig. 1).

### Detection of horizontal gene transfers between *P. nalgiovense* and *P. salamii*

We aimed to detect very recent horizontally transferred regions (HTRs), present in both *P. nalgiovense* and *P. salamii*, *i.e.*, resulting from events (HGTs) during domestication by humans that could potentially be involved in parallel adaptation to the dry-cured meat environment. The genomic regions resulting from recent horizontal gene transfers are expected to be almost identical between the distantly related *P. salamii* and *P. nalgiovense* species, would not necessarily be present in all strains and would be absent from many non-dry-cured meat species (e.g. Cheeseman et al., 2014).

We searched for recent HTRs between *P. nalgiovense* and *P. salamii*, by identifying thresholds above which regions could be considered to be more similar across a longer length than expected from the phylogenetic distance between the species. For comparisons of genomic sequences of more than 1,000 bp between the non-dry-cured meat sister species and their respective sister species (i.e., comparisons of *P. chrysogenum* and *P. rubens*, two species formerly considered as a single species, with *P. olsonii*), 98.21% was the highest level of similarity (Fig. S5A). We therefore set the identity threshold for identifying potential HTRs at 98%, because lower levels of similarity might be expected to occur by chance for such long sequences between sister species. The longest of the nearly identical sequences was 11,414 bp long (Fig. S5B). However, as the second longest sequence was only 6,972 bp long, we decided to set the threshold length for identifying potential HTRs to 7,000 bp, because shorter fragments with such high levels of similarity might occur by chance between the sister species found in dry-cured meat products.

Our analysis of genomic regions of more than 7,000 bp in length with more than 98% similarity between pairs of *P. nalgiovense* and *P. salamii* strains identified 26 putative HTRs. These putative HTRs were between 7,224 bp and 496,686 bp long (Table S3). The cumulative total length of the putative HTRs was 1,447,064 bp, including 145,640 bp assigned to transposable elements (TEs) of known families and 558,247 bp assigned to TEs of unknown families (10% and 38.6%, respectively; Table S3). We identified 34 (65%) of the 52 (2x26) consensus HTR extremities (the very last bases) as belonging to TEs, and all but three of the extremities had a detected TE within 1,000 bp, suggesting a possible role for TEs in the horizontal transfer of some of these regions. After the removal of all TEs of known and unknown families identified in the *P. nalgiovense* and *P. salamii* reference genomes (“repeat-masked” analyses), three putative HTRs had lengths below 500 bp and were therefore discarded for subsequent analyses (Table S3).

Plotting the identified putative HTRs on the contigs of the *P. salamii* and *P. nalgiovense* reference genomes showed that some groups of HTRs clustered very close to each other (Fig. 2), suggesting that they may represent a single horizontal transfer event in these strains (in particular HTRs 4, 5, 6, 7 and 8). The other HTRs were located at the edge of large contigs (e.g. Pnal 04, Pnal 07, Pnal 11, Pnal 19) or in very small contigs and may therefore also actually belong to the same regions but being not assembled due to repetitive elements. We found five occurrences of the DUF3435 domain, characteristic of the Starship elements involved in HGTs in fungi (Gluck-Thaler et al., 2022), in the two large clusters of HTRs. The HTRs were collinear between the reference genomes of *P. salamii* and *P. nalgiovense* despite widespread genomic rearrangements elsewhere (Fig. 2), which brings further support to the inference of recent horizontal transfers. We present hereafter results separating the 23 initially identified HTRs as no criterion appeared optimal to decide which should be merged. There has nevertheless likely been just a few independent horizontal transfer events with cumulative length of recent HTRs reaching almost 1.5 Mb.

**Figure 2.**
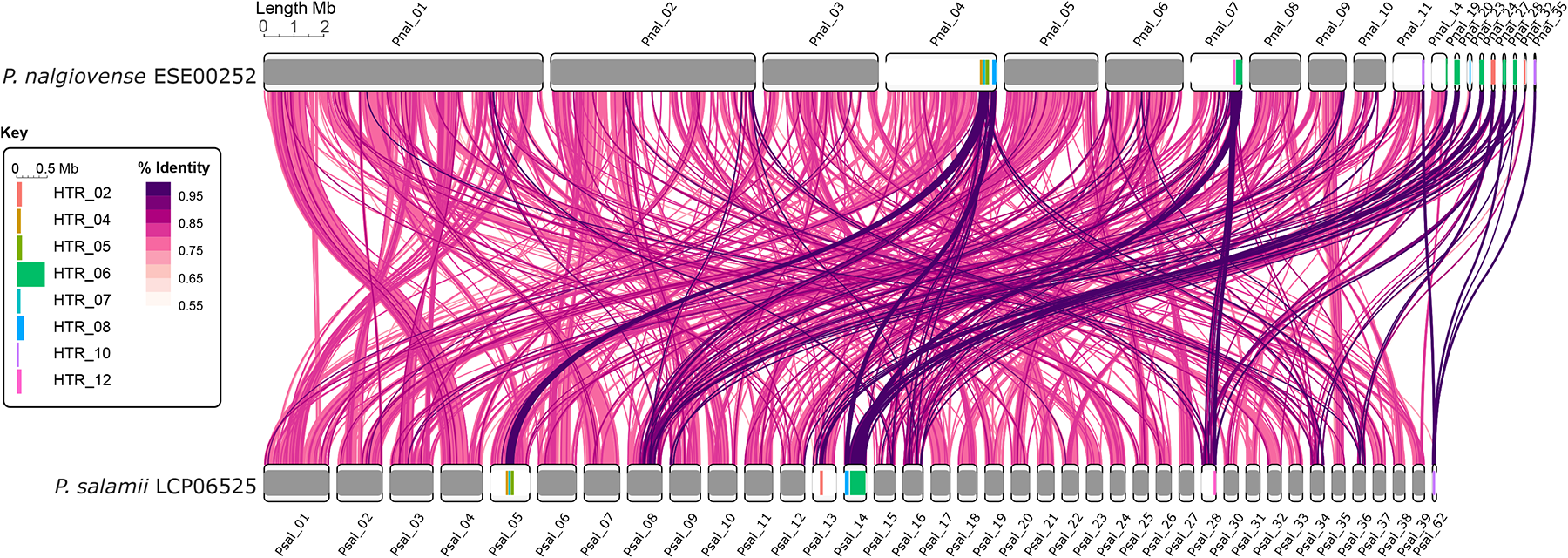
Synteny and similarity between the *Penicillium nalgiovense* and *P. salamii* reference genomes and location of the horizontally transferred regions (HTRs). Contigs based on long read assemblies for the genomes of *P. nalgiovense* ESE00252 are shown at the top and *P. salamii* LCP06525 at the bottom. Only contigs larger than 300 kbp or matching a HTR are shown. The links indicate matching blocks larger than 1 kbp, colored according to averaged nucleotide identity (%) between *P. salamii* genomes and *P. nalgiovense* ESE00252. Only links present in at least half of the pairwise comparisons are shown, or those matching HTRs and being at least half covered by non-overlapping contigs (to avoid taking repeated elements as evidence of presence). The HTRs are identified on the contigs by colors as indicated on the figure key, on which HTRs are magnified 2X for clarity.

We identified 283 coding genes in the putative HTRs remaining after repeat masking, 199 of which yielded hits in one of the egg-NOG databases, mostly with the fungal egg-NOG database (87.4%, 173 genes), suggesting that most of the putative HTRs identified originated from fungi. The other 25 genes had best hits in the eukaryote egg-NOG database (12.1%) or the principal egg-NOG database (0.5%). They could, therefore, also be of fungal origin, but better annotated in more phylogenetically distant groups (e.g. mammals), resulting in a best match at a higher taxonomical level.

All but the two most divergent *P. nalgiovense* strains contained the same putative HTRs, whereas there was more variability in terms of presence/absence of HTRs among *P. salamii* strains, with some strains even carrying none of the putative HTRs (Fig. 3; Figs. S6-8). The number and length of the putative HTRs largely accounted for the differences in genome size between *P. salamii* strains (adjusted R^2^: 0.67; Fig. S9). The fraction of each putative HTR present in genomes differed between strains (Fig. 3; Figs. S6-8) but the present sequences were always very similar to that of the reference putative HTR (99-100% identity; Fig. 3; Figs. S6-8 and Table S4). Among the eight largest HTRs, only HTR5 had around a third of its sequence with identity below 98% among some *P. nalgiovense* strains (Fig. 3). There were two different versions of the sequence in *P. nalgiovense* (with 89% identity between them), where the strains were in this region either 100% identical to one or the other. They may therefore correspond to recent HGTs of the same region from two different donor species or populations. In some strains of *P. nalgiovense*, HTRs 6, 8 and 10 did not show 100% identity along their entire sequences due to a large number of repeated sequences and therefore ambiguous mapping. We identified the same ∼6 kbp LTR-like sequence inserted in HTRs 7 and 8 of *P. nalgiovense* strain ESE00252 (Fig. 3). The finding of near 100% identity for all HTRs between strains and species (excluding ambiguous alignments due to repeats or repeat insertions) suggests HGTs during domestication by humans and preclude any dating attempt.

**Figure 3.**
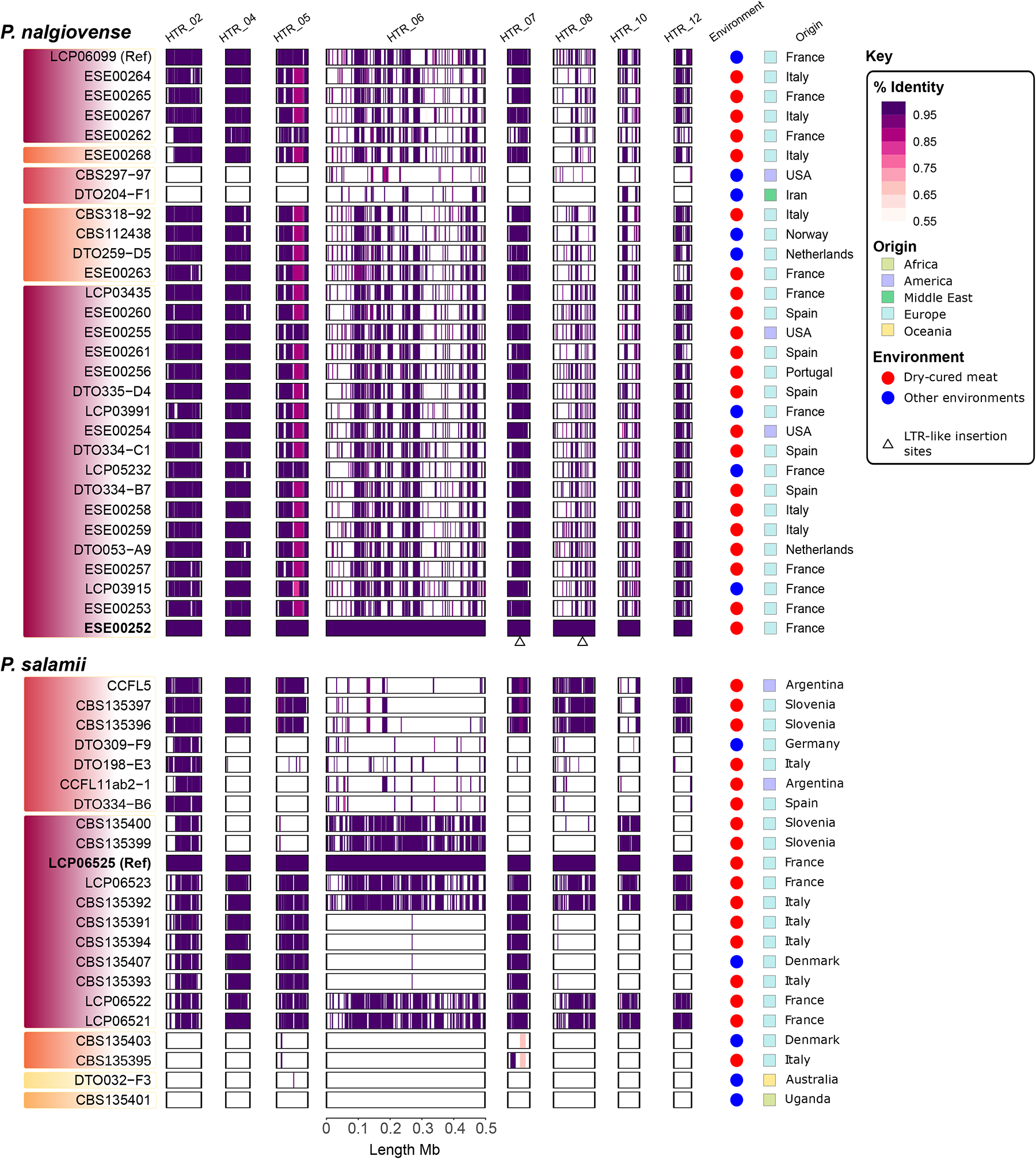
Presence and identity level of the eight largest putative horizontally transferred regions (HTRs) in *Penicillium nalgiovense* (top) and *P. salamii* (bottom) genomes. The color darkness on the contigs indicates the average nucleotide identity (%) of HTRs present in each strain compared to the reference genomes. White indicates complete lack of the corresponding region or ambiguous mapping (mostly due to repetitive elements). The length of each rectangle is proportional to the size indicated by the scale. The shades under the names of the strains on the left are colored according to the genetic clusters as in Figure 1. The strain geographical origins are shown on the right by the color of the squares (blue: Europe, purple: America, yellow: Australia, light green: Africa, green: Middle East). The color circles indicate environment of isolation, with dry-cured meat in red and other environments in blue. Strain codes in bold are the long-read based assemblies used for synteny analyses in Figure 2. Triangles below HTR_07 and HTR_08 in the ESE00252 strain indicate insertions of the very same LTR-like retrotransposon. See Figures S6 to S8 for the information on presence/absence of all the HTRs in all the strains and Table S4 for identity percentage and unambiguous alignment fraction.

We constructed a phylogenetic species tree of 68 genomes from other *Penicillium, Aspergillus* or *Monascus* species with 1102 single-copy orthologous genes. The resulting species tree (Fig. 4) was consistent with previous studies (Houbraken et al., 2014; Perrone et al., 2015; Steenwyk et al., 2019) and was well supported. The putative HTRs were absent from most of the other species analyzed (Fig. 4, Figs. S10-S12). Only 25 species carried parts of at least one putative HTR, and 19 species carried parts of more than one putative HTR. The species carrying the largest number of the 23 putative HTRs belonged to genus *Penicillium* (Fig. 4, Figs. S10-S12). Two *Penicillium* species had large fragments of seven putative HTRs: *P. biforme* and both varieties of *P. camemberti* (Fig. 4, Figs. S10-S12). Interestingly, these two species belong to a species complex found in dry-cured meat and sometimes even used for inoculation for dry-cured meat maturation, but mostly used for cheese production. The two varieties of *P. camemberti,* var. *camemberti* and var. *caseifulvum,* are two closely related clonal lineages displaying very little variability, having diverged recently from *P. biforme,* which harbors higher levels of genetic and phenotypic variability (Ropars et al., 2020). Most of the other *Penicillium* species harbored only small parts of the putative HTRs. HTR 6 and HTR 15 were present in more species than the other putative HTRs and had similar distributions across species (Fig. 4, Figs. S10-S12). The similarity between the putative HTRs present in the different species and the reference putative HTR was very high, despite the phylogenetic distances between the species harboring them (typically more than 95%, Fig. 4, Figs. S10-S12), providing further support for the view that they result from horizontal gene transfers. Among the 304 genes present in inferred HTRs in at least one of the 54 species studied here, only 25 had a tree topology compatible with the species tree, among which only 12 genes were present in both *P. salamii* and *P. nalgiovense*; these 12 genes were scattered across ten different HTRs. This means that all HTRs have multiple genes whose genealogies are topologically significantly different from the species phylogeny, and that no putative HTR carries only genes with genealogies congruent with the species phylogeny. We can therefore be highly confident that the putative HTRs are the result of non-vertical transmission.

**Figure 4.**
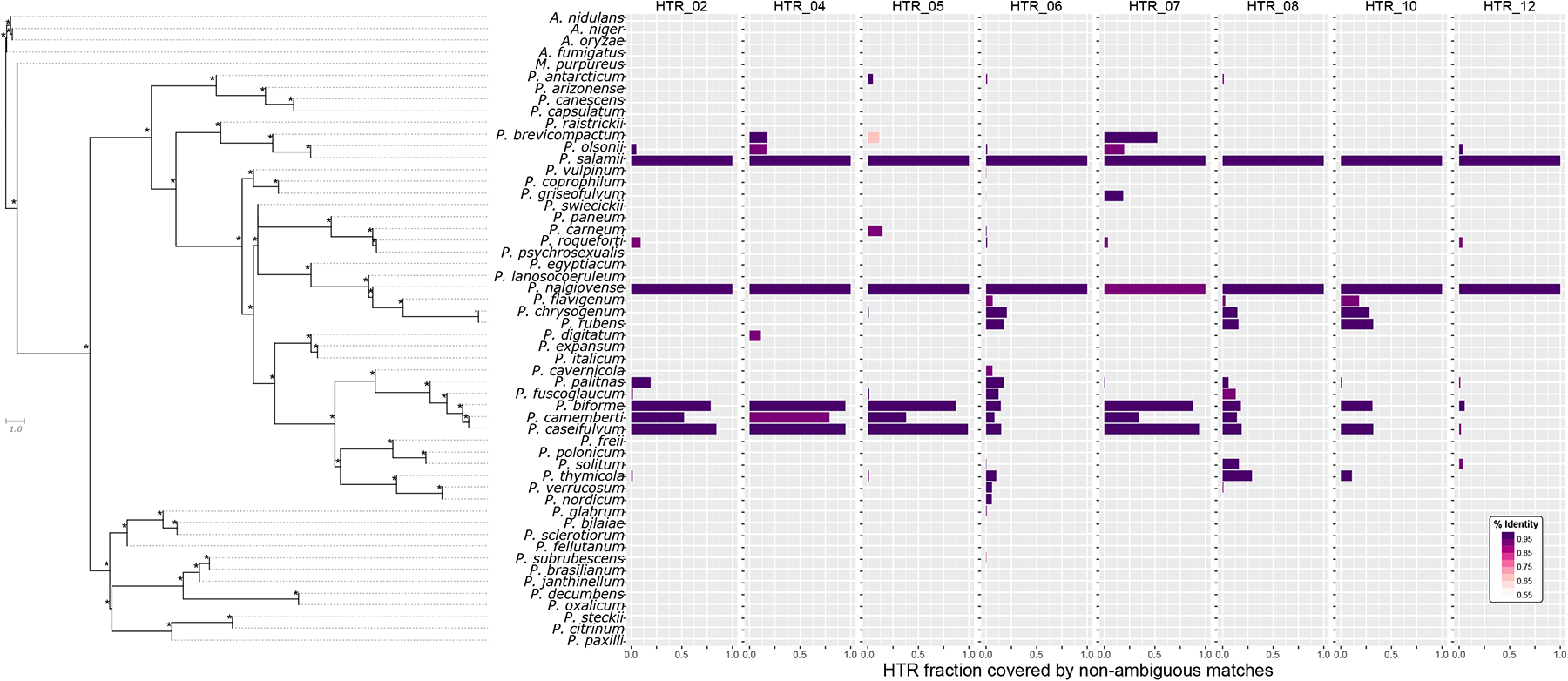
Presence and similarity of the eight largest HTR (horizontally transferred regions) across 54 *Penicillium*, *Aspergillus* or *Monascus* genomes. In each genome, the fraction of each HTR covered by non-ambiguous matches was obtained by dividing the total length of non-overlapping matches over the total length of the corresponding HTR. The percentages of identity (% identity) correspond to the number of identical bases to the consensus HTR sequences over the total matched length. The phylogenetic tree on the left was built from 1102 single-copy genes with Astral v5.6.3. (Zhang et al., 2018) and rooted on the *Aspergillus* clade. The distance in the internal branches is expressed in coalescent units. Stars on the right end of each branch correspond to a full support of the relationship between the four leaves surrounding this branch.

We, thus, identified genomic regions that were almost identical between distantly related species, absent in more closely related species, rich in TEs (accounting for a mean of 44% of the regions, ranging from 9 to 91%), including some firmly linked to known TE families, located in different regions of the genomes among species, or even the population, and having genealogical histories different from the species history. These findings provide compelling evidence that these regions resulted from horizontal gene transfers. The very high identity levels prevent using gene genealogies to identify the donor species for HTRs. The genetic diversity within species was not useful either to identify the donor species as we had only one or two genomes for each species, except *P. nalgiovense* and *P. salamii* in which they were nearly identical.

Given the distance separating species in our tree, the *P. nalgiovense* – *P. salamii* pair appeared to have a particularly high shared HTR content, in terms of total length. Only the *P. nalgiovense* – *P. biforme* pair had a longer total length of HTRs (Fig. S13), which, interestingly, also corresponds to a pair of species used in dry-cured meat production. More generally, all the species that were phylogenetically distantly related but had a large number of putative HTRs in common were known to be present and sometimes used in either cheese or dry-cured meat production (Fig. S13).

An analysis of functional categories in clusters of orthologous groups present in the HTRs identified eight categories present in most strains and four categories that were less present (Table S5). None of the *P. nalgiovense* or *P. salamii* strains displayed an enrichment in certain functions in the HTRs relative to the rest of the genome. The seven HTRs present in *P. biforme* and the two varieties of *P. camemberti* carried 42 and 47 genes, but only seven to 20 of them had predicted functions. These genes were involved in the cytoskeleton, transport and metabolism of carbohydrates, inorganic ions, lipids or secondary metabolites, post-translational modifications, or intracellular trafficking (Table S5). The two regions present in the largest number of species (HTR6 and HTR15) carried 62 and 11 genes, respectively, 45 and 10 of which, respectively, were annotated, and only 24 and seven, respectively, with predicted functions, belonging to 11 different COGs. The carbohydrate transport and metabolism category was the only one containing five genes, three being the maximum for the other categories. Therefore, some gene functions in the HTRs may be beneficial for dry-cured meat production as they are involved in the metabolism of meat nutrients, but the lack of significant enrichment renders conclusions challenging.

### Detection of positive selection

For *P*. *salamii*, we found no evidence of genome-wide positive or negative selection or a recent change in demography (Tajima’s *D* = 0.148, not significantly different from zero, 95% confidence interval [-1.657; 2.153]; *p*-value: 0.432). By contrast, for *P. nalgiovense*, we found evidence of widespread selective sweeps, background selection or recent population expansion (Tajima’s *D =* -2.834, 95% confidence interval [-4.641; -0.814]; *p*-value: 6e-08).

We looked for footprints of selection outside of HTRs with a Bayesian generalization of the McDonald and Kreitman test (McDonald & Kreitman, 1991), which identifies genes in which amino-acid changes are more or less frequent than expected under neutrality (inferring positive or purifying selection, respectively), by detecting higher or lower non-synonymous divergence from an outgroup than expected based on observed synonymous and nonsynonymous SNPs in the in-group. We analyzed 7,155 genes present in all, or all but one of the strains of *P. nalgiovense*, and 7,010 genes present in all, or all but one of the *P. salamii* strains. We found that 1,572 of these genes in *P. nalgiovense* and 466 in *P. salamii* were evolving under purifying selection, and that 71 of these genes were common to the two species (Table S6). We identified 751 genes in *P. salamii* and 30 in *P. nalgiovense* evolving under positive selection, with only three of these genes common to the two species (one with no predicted function, one predicted to encode a phosphotransferase and the third predicted to be involved in meiosis; Table S6). The genes under positive or purifying selection displayed no specific clustering within genomes. We detected a significant excess of positive selection in *P. nalgiovense* for genes involved in secondary metabolite biosynthesis, transport, and catabolism, and in *P. salamii* for genes involved in transcription, posttranslational modification, protein turnover, chaperones, signal transduction and defense mechanisms (Table S6). We detected a significant excess of purifying selection for functions involved in the transport and metabolism of carbohydrates and amino acids in *P. nalgiovense* and for functions involved in energy production and conversion, and in secondary metabolite biosynthesis, transport, and catabolism in *P. salamii* (Table S6).

### Phenotypic experiments to test the hypothesis of domestication

We analyzed several phenotypic traits, to test the hypothesis that dry-cured meat fungi are domesticated: growth and spore production on malt extract agar medium at different salt concentrations, colony color on malt extract agar medium, and lipolytic and proteolytic activities. If dry-cured meat fungi have been domesticated, then we would expect to see: 1) significant differences between dry-cured meat and non-dry-cured meat strains within each species, and 2) no significant differences between the non-dry-cured meat strains and the sister species not used for dry-cured meat production. These findings would indicate that only the fungi used for dry-cured meat production had evolved specific phenotypes related to adaptations to the dry-cured meat substrate. We found that *P. nalgiovense* and *P. salamii* contained horizontally transferred regions common to the *P. biforme* complex, which can also be used to inoculate dry-cured meat. We therefore also included *P. biforme* in phenotypic assays. We investigated the possibility that there were phenotypic differences between dry-cured meat and non-dry-cured meat strains within each of the species used for dry-cured meat maturation: *P. nalgiovense, P. salamii* and *P. biforme.* We also investigated whether there were differences between the non-dry-cured meat strains (the dry-cured meat strains were excluded from this analysis) and sister species not used for dry-cured meat production (i.e., *P. olsonii* for *P. salamii, P. chrysogenum* for *P. nalgiovense* and *P. palitans* for *P. biforme*). For the comparison of sister species, we included a “clade” effect in this test, with clades corresponding to sister-species pairs, to control for phylogenetic relatedness in the analysis.

### Salt tolerance: effect of salt concentration on growth and spore number

In comparisons of growth between non-dry-cured meat populations and their sister species in the three clades under various salt concentrations, we found significant effects of salt concentration, clade and an interaction between clade and salt concentration, but no effect of species identity within a clade (Table 2, Fig. 5). Thus, the non-dry-cured meat strains and their sister species had similar responses to salt. All pairs of species grew faster in the presence of 2% salt than without salt, but growth rates rapidly decreased with increasing salt concentration beyond 2% (Table 2, Fig. 5).

**Figure 5.**
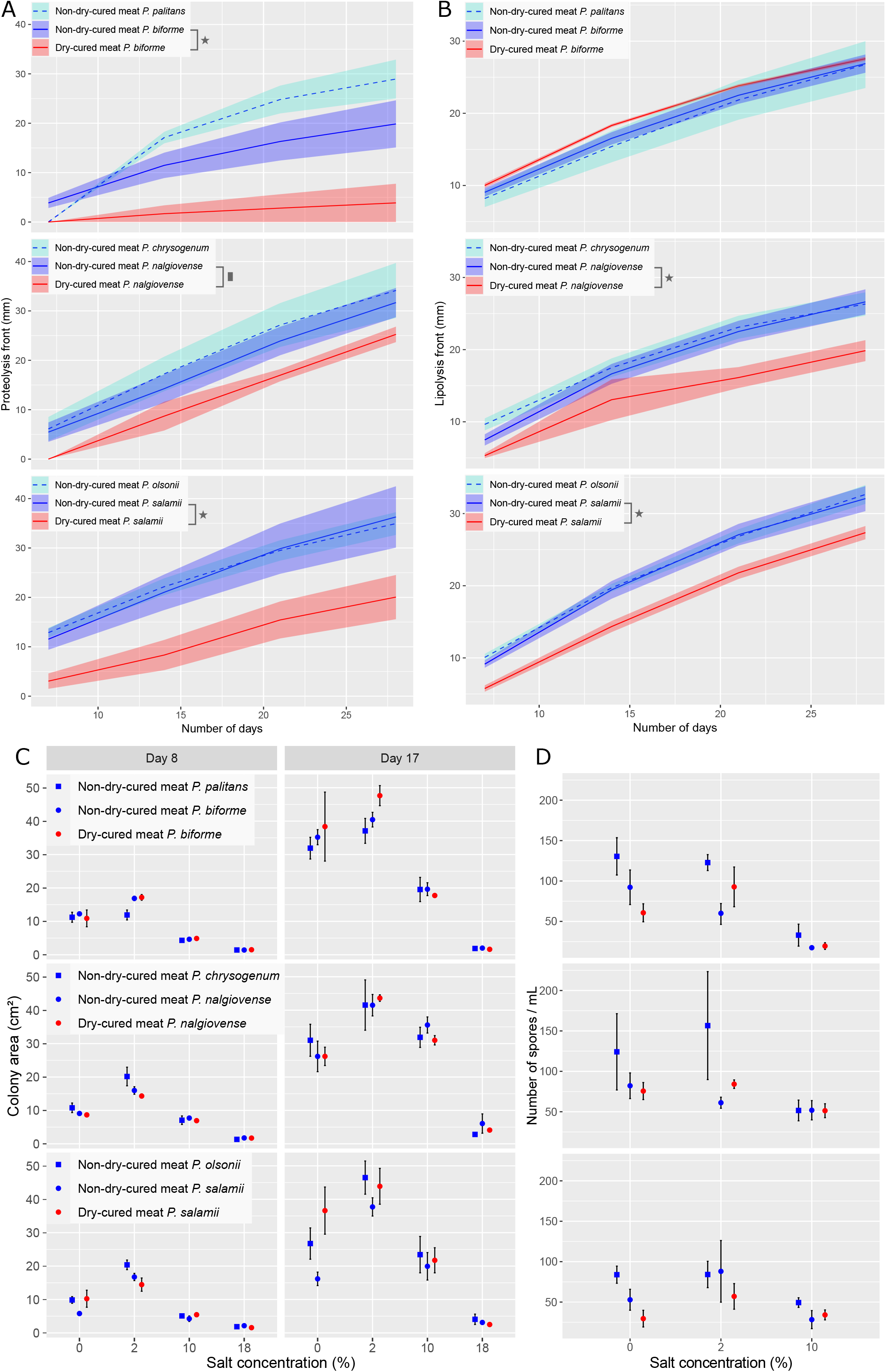
Proteolysis (A) and lipolysis (B) dynamics, and growth (C) and spore production (D) in response to different salt concentrations in *Penicillium biforme*, *P*. *nalgiovense* and *P*. *salamii* and their respective sister species, *P. palitans, P. chrysogenum* and *P*. *olsonii*. (A) and (B): The x-axis represents the number of days since the start of the experiment, the y-axis represents the front of lysis for proteolysis or lipolysis in mm in measure tubes. The colored area around each logistic line represents the standard error, and colors correspond to the environment of isolation (red=dry-cured meat, blue and turquoise=other environments excluding dry-cured meat). Stars indicate significant post-hoc tests (p-value < 0.05), squares marginally significant ones (0.05 < p-value < 0.1). (C) Growth rates represented as colony area in cm^2^ at days 8 (left panel) and 17 (right panel) post-inoculation at different salt concentrations (X axis). (D) Spore production is estimated at day 17 for the same species and the same salt concentrations. In (C) and (D), circles indicate species used for dry-cured meat production, i.e., *P. biforme*, *P. nalgiovense* and *P. salamii*, and squares indicate their respective sister species not used for producing dry-cured meat (*P. palitans*, *P. chrysogenum* and *P. olsonii*, respectively). The strains were collected either from dry-cured meat (red) or from other environments excluding dry-cured meat (blue). The error bars correspond to the standard error. We did not detect any significant difference between dry-cured meat and non-dry-cured meat samples in each species with post-hoc tests.

**Table 2.**
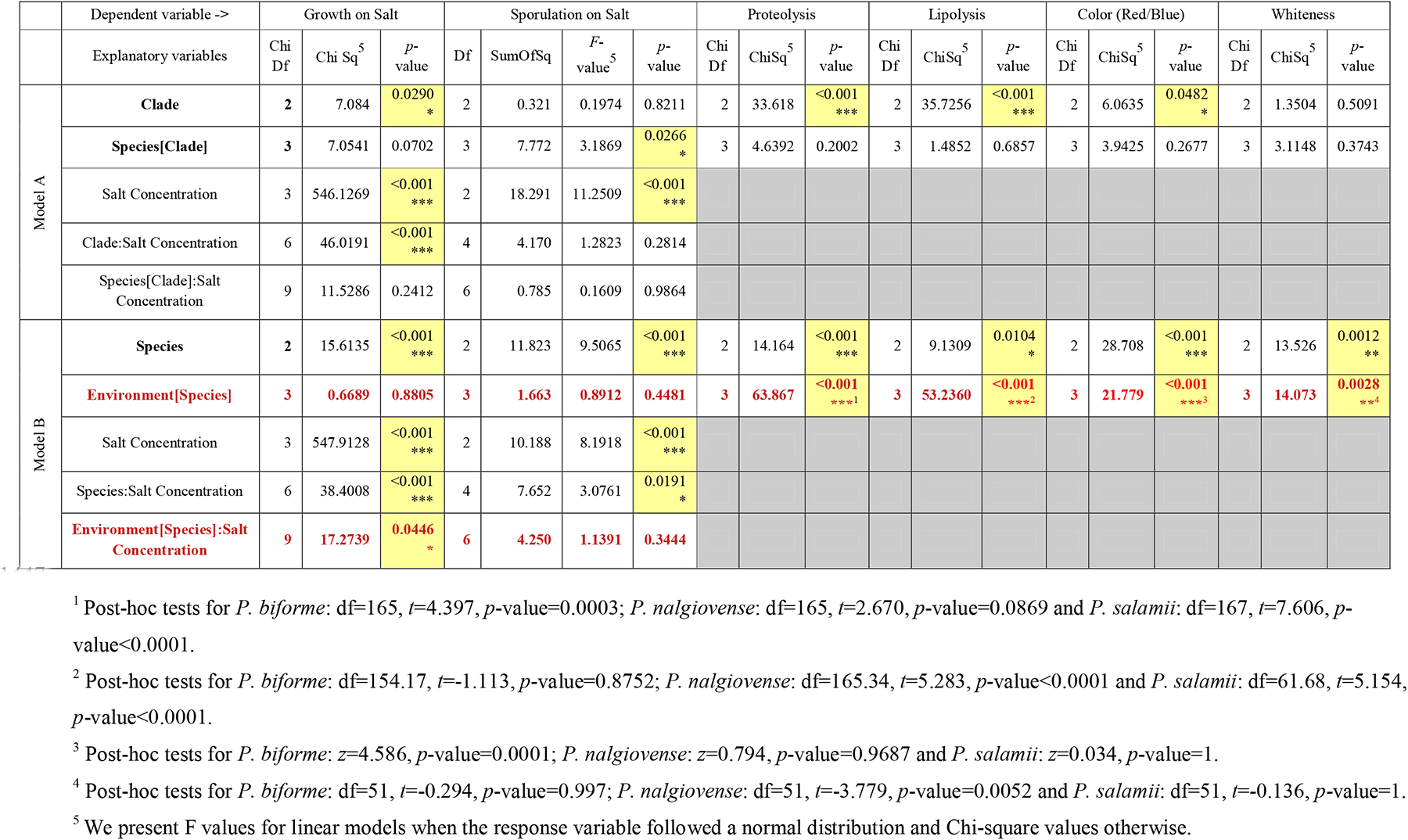
Statistics on the phenotypes measured in this study. Model A corresponds to the model testing whether non-dry-cured meat population in species used for dry-cured meat harbored different phenotypes than their sister species and Model B aimed at testing whether non-dry-cured samples showed significant differences relative to dry-cured meat samples. A clade corresponds to the species used in dry-cured-meat production and their sister species. Environment effect corresponds to either dry-cured meat or non-dry-cured meat. Some variables being non-independent (a species belong to a clade, and an environment is characteristic of a species), they are included as nested parameters and shown with squared brackets. Interactions are represented by colons. Cells with red bold font represent the main focus of the test, that is whether there is a difference between dry-cured meat and non-dry-cured meat populations. Cells with yellow background highlight significant differences with p-value < 0.05. Stars after the p-values correspond to a value between 0.05 and 0.01, between 0.01 and 0.001, and below 0.001 for one, two and three stars respectively.

In the analysis including dry-cured meat strains, we found no difference in growth between populations from dry-cured meat and populations from other environments (Table 2, Fig. 5). We found a significant interaction between isolation environment and salt concentration, a significant interaction between species and salt concentration and separate independent effects for salt concentration and species. This suggests that salt tolerance levels may differ between dry-cured meat and non-dry-cured meat strains (Table 2, Fig. 5). However, no post-hoc test of the impact of isolation environments was significant, in any species. The strongest difference between different isolation environments was for *P. salamii*, in which dry-cured meat strains grew better than non-dry-cured-meat strains in the absence of salt, although this difference was not significant in a post-hoc test (df =327; *t*=-3.285; *p*-value =0.1533).

We also measured spore production after 17 days of growth on malt extract agar medium containing various concentrations of salt. Spore numbers were very low on medium containing 18% salt. We therefore removed this category from the analysis. An analysis including only non-dry-cured meat strains and their sister species revealed a significant and strong effect of salt, with higher concentrations impairing spore production (Table 2, Fig. 5); we also found a weaker, but nevertheless significant within-clade species effect (Table 2, Fig. 5). In the analysis limited to dry-cured meat species, we found no difference between populations from dry-cured meat and those from other environments (Table 2, Fig. 5). We found a significant interaction between species identity and salt concentration (Table 2, Fig. 5). Significantly higher levels of sporulation were observed for *P. nalgiovense* than for *P. salamii,* but only in the total absence of salt (df=113, *t*=3.416, *p*-value=0.0240).

### Proteolysis and lipolysis activity

We investigated whether the dry-cured meat strains had evolved specific proteolysis and/or lipolysis activities in *P. nalgiovense, P. salamii* and *P. biforme.* A comparison of species without dry-cured meat strains revealed a significant effect of clade on both proteolysis and lipolysis (Table 2, Fig. 5). We detected no within-clade effect of species: the non-dry-cured meat populations and their sister species had similar levels of activity (Table 2, Fig. 5).

In the analysis including both dry-cured meat and non-dry-cured meat strains, the within-species effect of the isolation environment was significant for both proteolysis and lipolysis, and there was also a significant species effect (Table 2, Fig. 5). Post-hoc tests showed that, in *P. biforme* and *P. salamii,* proteolysis was significantly slower in strains from dry-cured meat than in strains from other environments (Table 2). We observed a similar tendency in *P. nalgiovense,* but it was not significant. Lipolysis rates were also lower in strains from dry-cured meat than in those from other environments in *P. nalgiovense* and in *P. salamii*, but no such effect was detected in *P. biforme* (Table 2).

The differences between dry-cured meat and non-dry-cured meat strains, together with the lack of difference between the non-dry-cured meat populations and their sister species suggest that dry-cured meat strains have evolved specific catabolic abilities, constituting domestication (Table 2, Fig. 5).

### Brightness and color

We tested whether colony color was closer to white in dry-cured meat populations, as this may have been a criterion of selection by humans or pigments may not be useful anymore in the human-made environment, where UV are less present (Lin & Xu, 2020). We found no significant species or clade effect in the analysis including only non-dry-cured meat strains and their sister species (Table 2, Fig. 6). In the analysis including all strains of the dry-cured meat species, we found a significant effect of species and of isolation environment within species on the distance from whiteness in the RGB (red, green and blue) space (Table 2, Fig. 6). Post-hoc tests showed that only *P. nalgiovense* colonies from dry-cured meat were significantly closer to white than colonies from other environments (Table 2). Visual inspection, indeed, suggested that the colonies on malt extract agar medium were typically white with a smooth surface in *P. nalgiovense* strains from dry-cured meat, whereas the colonies were more greenish, with a smooth or cotton-like surface in *P. salamii* (Fig. 6). These results indicate that colony color in the dry-cured meat population in *P. nalgiovense* has evolved toward greater whiteness.

**Figure 6.**
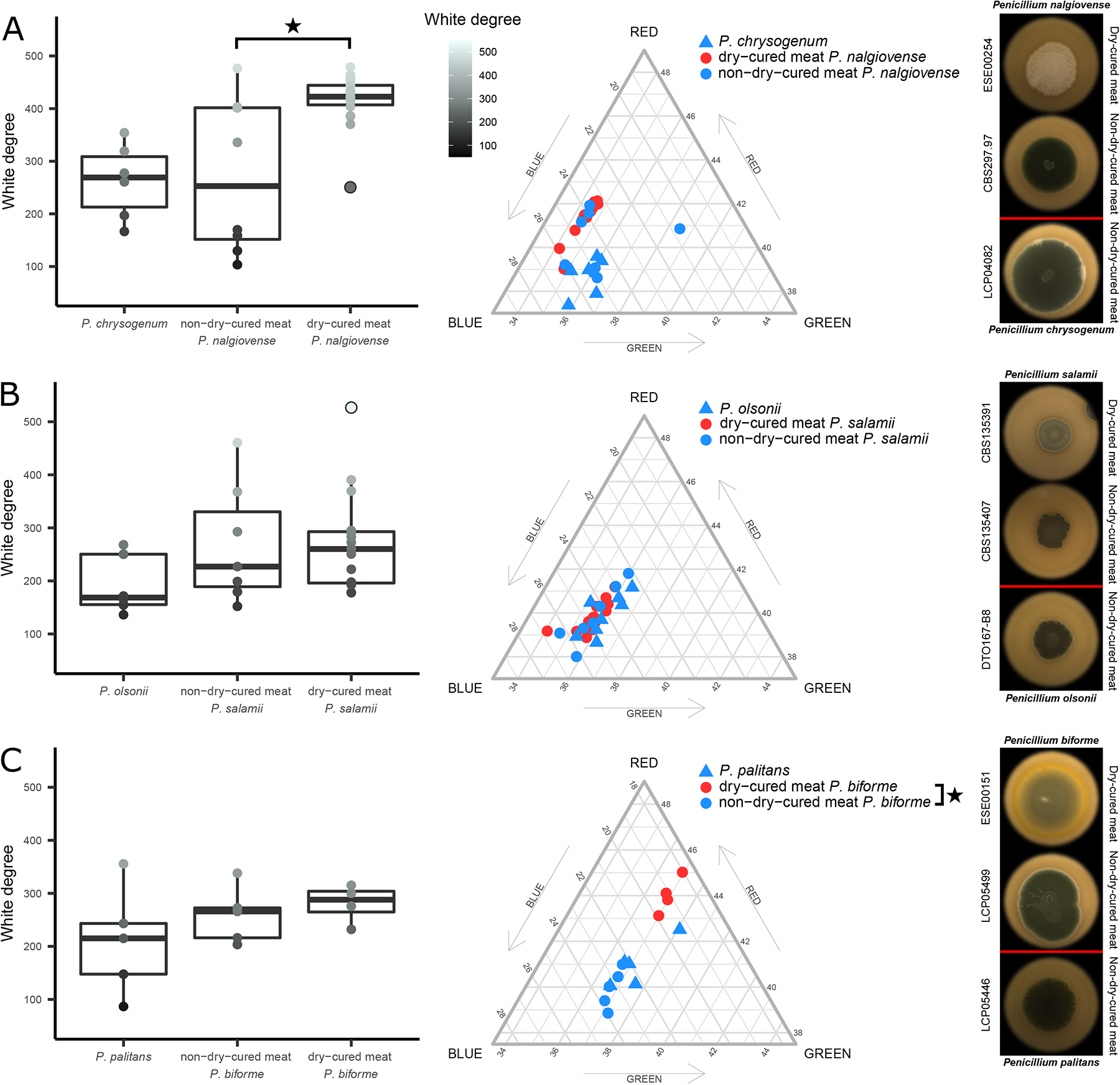
Degree of white, color composition and pictures on malt extract agar medium of *Penicillium nalgiovense* and *P. chrysogenum* (A)*, P. salamii* and *P. olsonii* (B)*, P*. *biforme* and *P*. *palitans* (C), separated by their environment of isolation. The left panel plots show the degree of white on a scale from 50 to 550 (represented as dark grey to light grey). The whitest point for *P. salamii* is circled in black for clarity. The middle panel shows the distribution among the red, blue and green values in the color of each *Penicillium* strain. The colors and shapes of the points correspond to the environment of isolation (red circle=dry-cured meat, blue circle=non-dry-cured meat, blue triangle=non-dry-cured meat sister species). Stars indicate significant post-hoc tests (p-value < 0.05). The right panel shows pictures of *P. nalgiovense, P. salamii, P. biforme* and their respective sister species on malt extract agar medium: the first row corresponds to strains collected on dry-cured meat, the second (and the sister species) to strains isolated from other environments.

We then investigated whether the dry-cured meat strains had evolved specific colony colors in terms of their red, green, and blue (RGB) components. In analyses limited to non-dry-cured meat strains and their sister species, the colors of fungal colonies were significantly affected by clade identity but not by the species within a clade (Table 2, Fig. 6). In analyses limited to dry-cured meat species, both isolation environment and species had significant effects on colony color (Table 2, Fig. 6). Post-hoc tests showed that *P. biforme* dry-cured meat strains were redder and/or less blue (Table 2), whereas the differences were not significant in the other two species. Thus, *P. biforme* dry-cured meat strains have evolved to have less blue colonies.

### Mycotoxins and penicillin production

We detected none of the mycotoxins or penicillin assessed (andrastin A, ermefortins A & B, (iso)-fumigaclavin A, meleagrin, mycophenolic acid, ochratoxin A, penicillic acid, penicillin G, penitrem A, roquefortine C and sterigmatocystin) in any of the *P. salamii* and *P. nalgiovense* dry-cured meat strains tested (Table S7). None of these extrolites were detected in *P. biforme* in a previous study (Ropars et al., 2020). All the sequenced strains from both *P. salamii* and *P. nalgiovense* harbored genes for the entire pathway for penicillin production in their genomes (*PCBAB, PCBC, penDE* and *phl* genes, and the *RFX* transcription factor gene).

None of the extrolites were detected in *P. olsonii*. We observed two distinct extrolite production profiles in *P. chrysogenum*, the sister species of *P. nalgiovense*. Five strains produced low levels of roquefortine C and no other extrolite (Table S7). The other nine *P. chrysogenum* strains produced high levels of andrastin A, meleagrin, roquefortine C and penicillin G.

### Possible feral strains

We tried to check whether some strains could be feral, i.e. escaped from dry-cured meat and isolated from other environment, with the rationale that they would be genetically and/or phenotypically close to dry-cured meat strains but isolated in another substrate. Among the six *P. nalgiovense* potential “feral” strains (i.e. closely related to dry-cured meat strains: CBS112438, DTO259.D5, LCP6099, LCP03915, LCP03991, LCP05232) four were tested for all the phenotypic traits (CBS112438, DTO259.D5, LCP03915, LCP05232) and one for the color (LCP03991). Four of them (CBS112438, LCP03915, LCP03991, LCP05232) exhibited phenotypes close to those of dry-cured meat strains (lower proteolysis and lipolysis, whiter color), and it was therefore not possible to rule out a feral origin. In contrast, the strain DTO259.D5 displayed phenotypes really similar to other non-dry-cured meat strains (high proteolysis and lipolysis, darker color), suggesting that it was a genuine wild strain rather than a recent escape from an industrial environment or that *P. nalgiovense* is able to revert back quickly to wild phenotypes. Two *P. salamii* strains were also genetically related to dry-cured meat strains despite having been collected in other environments (DTO309.F9 and CBS135407). Both were phenotyped and did not differ from the non-dry-cured strains, ruling out a potential “feral” origin.

## Discussion

The use of dry-curing as a method for conserving a perishable raw meat material can be traced back to Greek and Roman times (Frost, 1999). However, it remains unclear when the inoculation of dry-cured meat casings with *Penicillium* fungi began. *Penicillium nalgiovense* is the commonest species in the dry-cured meat food industry (Andersen & Frisvad, 1994; Larsen & Breinholt, 1999; Sunesen & Stahnke, 2003). *Penicillium salamii* was recently identified as an alternative ripening agent and has only been used or recovered from local dry-cured meat production sites, often producing naturally fermented meat products (Berwal & Dincho, 1994; M. Coton et al., 2021; Sunesen & Stahnke, 2003). This difference in usage was reflected in our samples from dry-cured meats, for which *P. nalgiovense* was the most frequently isolated species, followed by *P. salamii.* These two most abundant species have however not been found to co-occur in our samples despite six cases of species co-isolation, suggesting that they may be used as alternatives for dry-cured meat production. The wider use of *P. nalgiovense* may explain the difference in genetic diversity and the differentiation between the two species: the very low level of genetic diversity in *P. nalgiovense* may result from the selection of a single clonal lineage widely used in commercial dry-cured meat production, resulting in a strong bottleneck. Alternatively, the low level of genetic diversity may be explained by a possible relatively recent introduction of this species into Europe. Indeed, we had only four strains from outside Europe (USA and the Middle East), two of which were genetically different from the main lineage. *Penicillium nalgiovense* strains from non-food environments displayed no specific clustering, and most originated from human-related environments, such as the car upholstery, cheese, and dog bone. *Penicillium salamii* had a higher level of genetic diversity, consistent with the lack of wide-scale commercial use for this species (Magistà et al., 2016; Perrone et al., 2015). Genetic differentiation was observed in *P. salamii*, with clusters including different proportions of strains from dry-cured meats or other environments (e.g., sewage plant, tea leaves or soil), indicating that migration between dry-cured meat and other environments is still occurring.

We found no evidence of recombination in *P. nalgiovense*, with little LD decay, and only one mating type among the dry-cured meat strains, consistent with the use of asexual cultures of commercial starters. In the cheese fungi *P. camemberti* and *P. roqueforti*, cheese populations also have low levels of genetic diversity and clonal population structures (Dumas et al., 2020; Ropars et al., 2020). Non-cheese *P. roqueforti* populations have much higher levels of genetic diversity, with a larger number of recombination footprints, and higher levels of sexual fertility (Ropars et al., 2014, 2016), contrasting with our findings for *P. nalgiovense*. However, the alternative mating type was detected in the non-dry-cured meat strains. In *P. salamii*, we identified both mating types among the dry-cured meat strains, suggesting that sexual reproduction may occur in this environment. However, the LD patterns and the presence of only one mating type in each *P. salamii* population suggests an absence of recent sexual reproduction. The detection of both mating types in the two dry-cured meat species nevertheless opens up possibilities for future strain breeding.

Despite the lack of strong genetic differentiation between dry-cured meat strains and other strains, we found domestication footprints when comparing phenotypes relevant to dry-cured meat production. Indeed, dry-cured meat strains of both *P. salamii* and *P. nalgiovense* displayed slower proteolysis and lipolysis than strains from other environments. *Penicillium biforme* presented the same pattern for proteolysis. Excessive protein degradation is detrimental for dry-cured meat products, potentially rendering the product less firm and imparting bitter and metallic tastes (Toldrá, 1998), whereas excessive lipolysis would dry the product (Demyer et al., 1974) and have a negative impact on flavor. We found no major difference in lipolysis or proteolysis between the non-dry-cured meat strains of *P. salamii*, *P. nalgiovense* and *P. biforme* and their respective sister species, indicating that the dry-cured meat strains had evolved specific and convergent phenotypes, slower lipolysis and proteolysis, probably under selection by humans.

Fungal colonization of the surface of the meat casing has a direct impact on the observed color of the product, and this fungal coating has probably been subject to selection by humans for dry-cured meat fungal strains. Indeed, the visual appearance of the product is crucial for dry-cured meat-eaters, and a white color typically makes such products more attractive. *Penicillium nalgiovense* strains sampled from dry-cured meat were whiter than strains from other environments, consistent with the selection of dry-cured meat starters on the basis of color (Leistner, 1990). Dry-cured meat strains of *P. biforme* were less blue than non-dry-cured meat strains of this fungus. *Penicillium biforme* is widely used for cheese-making and is whiter than the closely related species *P. fuscoglaucum* on cheese media (Ropars et al., 2020). Whiter *P. biforme* strains may have been selected because it generates a more appealing aspect for both cheese and dry-cured meat production, especially as the strains isolated from these two types of product belong to the same population (Ropars et al., 2020). The clonal cheese fungus *P. camemberti* is known from historical records to have been selected as a white mutant to make Brie cheeses that looked less “moldy” (Ropars et al., 2020). White color in dry-cured meat fungi could alternatively be the result of relaxed selection in human-made environments because pigments would not be useful anymore to protect against UV or other stresses (Lin & Xu, 2020). By contrast, we observed no difference in color between *P. salamii* strains from different environments, probably due to weaker or more recent selection by humans, possibly even with colonization of the product principally from the environment rather than through active inoculation (Perrone et al., 2015), or due to contrasting selection pressures. Demand for local and traditional products is increasing, including that for dry-cured meats (Iaccarino et al., 2006), and this may have fostered an interest in less homogeneous and less white fungi on sausage casings. In the analysis limited to non-dry-cured meat strains, the lack of major color differences between the species used for dry-cured meat and their sister species indicated that the changes occurred specifically in dry-cured meat strains, further supporting the occurrence of domestication.

Phenotypic tests thus suggest that there may have been selection in *Penicillium* fungi for certain strains relevant to the production of dry-cured meat, with convergent evolution toward similar phenotypes, especially for lipolysis and proteolysis. Some of the strains isolated from non-dry-cured meat environments may be feral, but this would only make differences between strains from dry-cured meat and other environments more difficult to detect, this could not have generated or inflated the differences we found between strains from dry-cured meat and other environments, so our findings are robust. By contrast, we detected no specific traits relating to the response to salt concentration in dry-cured meat strains. *Penicillium* fungi, particularly those present in fermented food, are salt-tolerant and, therefore, able to grow in the salt-rich environments found in dry-cured meats and cheeses (Yadav et al., 2018); there may have been no further advantage of higher levels of salt tolerance or evolutionary constraints may have prevented the development of this tolerance.

None of the targeted extrolites, including mycotoxins and penicillin, were detected in *P. nalgiovense* or *P. salamii* dry-cured meat strains (M. Coton et al., 2021). However, the production of penicillin has been reported in some *P. nalgiovense* strains (Andersen & Frisvad, 1994) and the closely related species *P. chrysogenum* is known to produce mycotoxins (M. Coton et al., 2019; Frisvad et al., 2004). The lack of antibiotic or mycotoxin production in dry-cured meat strains may be due to relaxed selection in a human-made environment containing fewer competitors or to active selection for safe products. In *Penicillium* cheese fungi, some of the mycotoxin biosynthesis pathway genes harbor deletions, impairing these functions specifically in cheese populations and indicating selection for safe food production or relaxed selection (E. Coton et al., 2020; Gillot et al., 2017; Ropars et al., 2020). The absence of mycotoxin production in both *P. salamii* and its sister species *P. olsonii* suggests that this feature did not result from selection by humans in this clade. However, the culture medium is known to affect toxin production in *Penicillium* fungi (Frisvad et al., 2004).

We investigated genomic footprints potentially associated with adaptation to dry-cured meat production. Given the clonal structure of the dry-cured meat strains, especially in *P. nalgiovense*, it was not possible to perform genome-wide scans for the detection of selection based on either local decreases in genetic diversity or local genomic differentiation between populations. Indeed, under asexual reproduction, the selection of a beneficial variant results in a hitchhiking of the whole genome without the generation of typical local selective sweeps or islands of genomic differentiation. We therefore looked for footprints of horizontal gene transfers, which have been shown to be an important and frequent mechanism of rapid adaptation under selection by humans in fungi (Cheeseman et al., 2014; Marsit et al., 2015; Novo et al., 2013; Ropars et al., 2015). By comparing available *Penicillium* genomes from various environments, we found evidence of horizontal gene transfers, and, specifically, of horizontal transfers between *P. nalgiovense* and *P. salamii* despite the relatively large genetic distance between these species. The cumulative length of the HTRs shared between *P. nalgiovense* and *P. salamii* reached nearly 1.5 Mb. We could not generate genealogical trees of these regions to support the inference of horizontal transfer because most species completely lack these regions, but these absences, together with the very high levels of identity between these regions and their different locations within the genome, provide even stronger support for the inference of horizontal transfer. Indeed, introgression by interspecific hybridization would lead to the high-similarity regions being located at the same loci across species and being present in all species. In addition, given the very low collinearity (Fig. 2) and an estimate of 30 My divergence between *P. nalgiovense* and *P. salamii* (Houbraken et al., 2020; Perrone et al., 2015; Steenwyk et al., 2019), it is very unlikely that sexual reproduction could be successful. The inference of horizontal transfer is further supported by the presence of the DUF3435 domain in two large clusters of HTRs, as these domains are characteristic of the Starship mobile elements (Gluck-Thaler et al., 2022). By contrast to findings for cheese fungi, the HTRs were not specific to the dry-cured meat strains (Cheeseman et al., 2014; Ropars et al., 2015), which may be due to some non-dry-cured meat strains being feral strains, to gene flow between dry-cured meat and non-dry-cured meat populations, or to a horizontal transfer older than the divergence between these strains. In such cases, this could mean that humans chose fungi with desirable traits for dry-cured meat production, acquired by horizontal transfers before human selection. However, HTRs were nearly 100% identical between species and strains, indicating horizontal transfers in human times. Several HTRs also appeared to be present in other species, particularly in *P. biforme* but also in *P. camemberti*, which are also used commercially in the food industry, mostly for the inoculation of cheeses, but also in dry sausages (this study; Cheeseman et al., 2014; Ropars et al., 2015, 2020). The abundance of shared HTRs was higher in dry-cured meat species than in available *Penicillium* genome pairs, suggesting their possible selection in this rich, human-made environment, as previously reported for cheese *Penicillium* fungi (Cheeseman et al., 2014; Ropars et al., 2015) and *Saccharomyces* yeasts used in wine-making (Marsit et al., 2015). We were unable to identify any particular functions overrepresented in these regions relative to the rest of the genome, possibly due to the low proportion of genes with predicted functions or to the diversity of functions under selection. Other HTRs may be present in only *P. salamii* or only *P. nalgiovense* and involved in adaptation to the dry-cured meat environment but our approach cannot detect such HTRs. We were indeed mostly interested in parallel adaptation to dry-cured meat.

Despite the low level of genetic differentiation between strains from dry-cured meat and other environments, we were able to detect phenotypic differences, even in *P. nalgiovense,* in which overall genetic diversity was low. However, a few SNPs may have a strong phenotypic impact, and the insertion of transposable elements may also modify gene regulation. In order to detect genes under selection for adaptation to the dry-cured meat environment, we also ran tests to detect non-synonymous substitutions that were more or less frequent relative to synonymous substitutions than would be expected under neutrality. *Penicillium nalgiovense* had few genes under positive selection and many under purifying selection, consistent with a high frequency of asexual reproduction in this species and, thus, probable genome-wide hitchhiking effects and background selection. The number of genes under positive selection was 25 times greater in *P. salamii* than in *P. nalgiovense,* and the genes concerned were involved in different functions. This finding may reflect the lower adaptation ability of this species, with low levels of genetic polymorphism. A few categories of genes seemed to be overrepresented among the genes under positive or negative selection, including some that could be linked to the dry-cured meat environment (carbohydrate metabolism and transport, defense mechanisms and secondary metabolite production). It is therefore likely that, in addition to horizontal gene transfers, positive selection on genes already present in the genome has contributed to the adaptation to dry-cured meat in *P. salamii* and *P. nalgiovense*. Other types of adaptive changes may also have occurred that could not be studied here because of the low genetic diversity, such as selective sweeps in regulatory regions. Copy number variation could also be involved in adaptation (Heckel, 2022; Stalder et al., 2023; Weetman et al., 2018) but the use of short-read sequencing did not allow such analyses to be conducted here.

Overall, our findings suggest that selection by humans has induced convergent phenotypic changes in *P. biforme*, *P. salamii* and *P. nalgiovense*, and that *Penicillium* dry-cured meat fungi have acquired beneficial traits for dry sausage production. These fungi have, therefore, been domesticated. The low level of genetic differentiation between dry-cured meat and non-dry-cured meat strains is likely due to the domestication process being very recent. Currently, *P. nalgiovense* strains are cultivated and sold to be inoculated in dry-cured meat (e.g. https://www.lip-sas.fr/index.php/nos-produits/salaisons/nos-propositions), so that the control-over-reproduction component of domestication is clearly present. The convergence in the evolution of dry-cured meat populations was most striking for proteolysis, with lower levels of proteolysis in dry-cured meat populations for *P. nalgiovense*, *P. salamii* and *P. biforme*. Our findings, thus, add to the growing evidence of domestication in food fungi, such as in beer and wine *Saccharomyces* yeast, koji *Aspergillus oryzae* (Gibbons et al., 2012), and cheese *Penicillium* fungi (Dumas et al., 2020; Gallone et al., 2016; M. Gonçalves et al., 2016; Ropars et al., 2020; Sicard & Legras, 2011). Genomic analyses in *Saccharomyces cerevisiae* have revealed a differentiation of populations between wild and food environments, and even specific populations corresponding to different uses of yeast, for the fermentation of bread, beer, wine or cheese (Gallone et al., 2016; M. Gonçalves et al., 2016). The different clades have contrasting fermentation traits, such as the capacity to use malt, aroma production and ethanol production. In the blue cheese fungus *P. roqueforti*, population differentiation was also observed between cheese and other environments, with a more rapid growth of cheese strains on cheese medium, higher levels of lipolytic activity, more efficient cheese cavity colonization, more diverse and positive aromas, and a higher salt tolerance (Caron et al., 2021; Dumas et al., 2020; Ropars et al., 2015). In both *S. cerevisiae* and *Penicillium* cheese-making fungi, HGT events have been implicated in adaptation (Cheeseman et al., 2014; Ropars et al., 2015).

In conclusion, we found clear convergent adaptation in two distantly related dry-cured meat species of fungi, in terms of proteolysis and lipolysis, and in terms of shared horizontal gene transfers. Such studies on parallel adaptation to the same ecological niche are important for understanding the process of evolution and whether it is repeatable. However, only a few studies on a handful of models have investigated parallel adaptation. Studies on natural populations have shown that adaptation to similar ecological niches has led to the convergent evolution of phenotypes, such as size, color and form, in *Anolis* lizards, three-spined sticklebacks and cichlid fishes in African rift lakes, for example (Colosimo et al., 2004; Duboué et al., 2011; Elmer et al., 2014; Mahler et al., 2013; Salzburger, 2009). Dry-cured meat species of *Penicillium* therefore appear to be good models for investigating general questions about adaptation. We also found a substantial loss of diversity in dry-cured meat populations, as previously reported for cheese fungi (Dumas et al., 2020; Ropars et al., 2020). This lack of diversity may jeopardize future strain improvement and lead to degeneration. Our findings therefore also have implications for the food industry. The discovery of different phenotypes and the presence of two mating types in both species paves the way for strain improvement by conventional breeding, which could also allow studying the genomic bases of beneficial phenotypes for dry-cured meat making and generating further genetic and phenotypic diversity.

## Data accessibility

All sequencing data have been deposited to the European Nucleotide Archive under the accession PRJEB44534. The HTR sequences and their annotations are available under the publicly available gitlab project https://gitlab.com/abranca/consensus_htr.

## Acknowledgments

This work was supported by the ERC starting grant GenomeFun 309403 and Louis D foundation award to TG, a CNRS PEPS adaptation grant to AB and the FUNGADAPT ANRC19CCE20C0002 ANR grant to TG, AB and MC. YCL acknowledges a PhD grant from the France–Taiwan joint program Campus France. We thank Alice Feurtey, Fanny Hartmann and Fantin Carpentier for help with genome analyses. We thank MNHN for granting access to strains with the help of Manuela Lopez-Villavicencio and Sandrine Lacoste. We thank all the people who sent dry-cured meat casings from around the world. We thank the GenoToul platform for sequencing. We are also grateful to the GenoToul bioinformatics platform Toulouse Occitanie (Bioinfo Genotoul, doi: 10.15454/1.5572369328961167E12) for providing some computing resources. PacBio sequencing was conducted at the IGM Genomics Center, University of California, San Diego, La Jolla, CA. We thank Federico Laich and Giancarlo Perronne for providing *P. salamii* strains. Some sequence data (Table S2) were produced by the US Department of Energy Joint Genome Institute http://www.jgi.doe.gov/ in collaboration with the user community.

