## Supplementary file for "Domestication in dry-cured meat *Penicillium* fungi: convergent specific phenotypes and horizontal gene transfers without strong genetic subdivision"

**Supplementary Methods**

**Strain isolation, DNA extraction and species identification**

For strain isolation, dry-cured meat casing samples were collected from Europe, Asia, and America. To avoid mite contamination, we first put dry-cured meat casing samples in the freezer (-20°C) for three days. After freezing treatment, we deposited a piece of dry-cured meat casing onto a malt extract agar (MEA) medium (16g malt, 16g agar, 800 mL H20) for two to four days until fungal colonies were observed. For each Petri dish, we scraped the entire fungal material and placed it into 1 mL of 0.05% of Tween 20 (Sigma-Aldrich); we spread 100 µL of a diluted suspension (10^-3^ dilution) on MEA medium. This dilution step was used to isolate single-spore colonies on Petri dishes. We then selected colonies with *Penicillium*-like morphologies (Fig. 5) and purified them separately on MEA medium. All cultures were stored in 10% glycerol at -80°C. The Nagoya protocol on biodiversity and shared benefits did not apply to our strain collection, as all strains were isolated prior to law publications in their respective countries of collection.

For DNA extraction, monospore-derived *Penicillium* strains were cultured on MEA medium for seven days. Then 3-5 ml of 0.05 % Tween 20 was added to the Petri dishes to collect fungal tissues. DNA was extracted using the NucleoSpin soil kit for genomic DNA (Macherey-Nagel) following manufacturer instructions. DNA quality was assessed by measuring 260/230 and 280/260 nm ratios with a NanoDrop 2000 spectrophotometer (Thermo Fisher Scientific Inc), and DNA concentration was measured with a Qubit 2.0 fluorometer (Thermo Fisher Scientific Inc).

To identify the species isolated from dry-cured meat casings, *bt2a/bt2b* (Glass & Donaldson, 1995) primers were used to amplify a fragment of the β-tubulin gene, which is recognized as a discriminant marker for this purpose (Samson et al., 2004; Visagie et al., 2014). In addition, to identify the closely related species *P. biforme* and *P. fuscoglaucum*, previously grouped under the name *P.* *commune*, we used the PC4 and PC9 markers (Giraud et al., 2010). We distinguished the closely related species *P*. *chrysogenum* and *P*. *rubens* (Houbraken et al., 2011), by using the RPB2 (DNA-directed RNA polymerase II) marker (Visagie et al., 2014). PCRs were performed using a Bio-Rad thermal cycler with an initial denaturation of 2 min at 94°C, 40 cycles of 30 s denaturing at 94°C, 30 s annealing at 54°C and 40 s extension at 72°C with a final extension step at 72°C for 5 min. PCR fragments were sequenced at GENEWIZ Europe and sequences blasted (Blastn) against public databases (nucleotide collection nr/nt) for species identifications (Altschul et al., 1990; Sayers et al., 2019).

**Genome assemblies**

The complete long-read genome assemblies were performed using the SMRT analysis system v2.3.0 (Pacific Biosciences). Genome sequences were *de novo* assembled using the HGAP v3 (Chin et al., 2013) protocol, with a minimum seed read length of 1000, genome size of 30 Mb, target coverage of 20, and an overlap error rate of 0.03. Polished contigs were further error corrected using Quiver v1. A summary of raw data and assembly statistics is reported in Table S1. The protein-coding gene models were predicted with EuGene v4.2a trained for *Penicillium* (Cheeseman et al., 2014; Foissac et al., 2008; Sallet et al., 2019) and their putative function with eggNOG-mapper online (Huerta-Cepas et al., 2017) and the eggNOG 4.5.1. general database (Huerta-Cepas et al., 2015).

Short-read sequencing data were checked using FastQC (v0.11.6; Andrews, 2010). No adapter sequence could be detected. A first assembly round was performed using SOAPdenovo v2.04 (Luo et al., 2012) and a k-mer value of 71. To confirm species identity, we recovered part of the calmodulin and β-tubulin genes and of the ITS regions using blastn v2.2.29+ (Altschul et al., 1990) and the corresponding *P. nalgiovense* genes as references (accessions JX996974.1, KX928936.1 and KC009791.1), then identified species by comparing the partial genes obtained with the NCBI nucleotide database (nt) using blastn v2.2.29+ (Altschul et al., 1990; Sayers et al., 2019). We then compared all the assembled contigs and scaffolds to the NCBI nucleotide database (nt) using blastn v2.2.29+ (Altschul et al., 1990; Sayers et al., 2019) to identify those contaminated by other species.

Among the 55 genomes sequenced, we detected three *P. nalgiovense* samples (ESE00267, ESE00268 and ESE00262) as contaminated by *Debaryomyces hansenii*, a yeast almost systemically inoculated into dry-cured meats during production (Ravyts et al., 2012). Among *P. salamii* genomic data, one strain (LCP06522) was contaminated by *Geotrichum candidum*, and another one (CCFL11ab2.1) by a fungal species belonging to Phanerochaetaceae. By comparing each strain assembly with itself, we did not detect any large-scale duplication or assembly errors. We removed the reads corresponding to contaminants identified in the previous step with BBsplit from the BBmap package v38.22 (Bushnell, 2018), keeping reads mapping on both the expected reference genome and the identified contaminant one. We also kept reads mapping on neither the reference genome nor the contaminant one to detect regions that would be genuinely present in some *Penicillium* genomes but absent from the reference genomes, such as horizontally transferred regions that would be specific to some strains. It is a rather conservative approach, as the annotation of the potential horizontally transferred regions identified in this study did not match the contaminants identified at this step (see Results). With the reads cleaned from contaminants, we did a second assembly round with SPAdes v3.11.0 (Bankevich et al., 2012) and default k-mer parameters (21, 33, 55, 77), and then removed scaffolds shorter than 2,000 bp. We checked assembly quality with QUAST v5.0.2 (Gurevich et al., 2013). We performed pairwise genome alignments with NUCmer v3.1 and mummerplot v3.5 from the MUMmer3 package (Kurtz et al., 2004) with default parameters to detect large genomic rearrangements between strains within each species, and duplications within genomes. We predicted the genes as described for long-read assemblies.

**Detection of horizontal gene transfers**

In order to identify HGTs, we first searched for thresholds above which we could consider that regions were more similar across a longer length than expected given the species phylogenetic distance. All the following pairwise comparisons were performed using NUCmer v3.1 from the MUMmer3 package (Kurtz et al., 2004), keeping all anchored matches even in cases of multiple matches (--maxmatch option). We looked for blocks of 65-mer that were shared between sister species (i.e., 65 contiguous identical base pairs), which were later combined into longer regions if several 65 bp identical blocks were not separated by more than 90bp, and we retained such regions when they were at least 1,000 bp (default NUCmer parameters). We characterized the length and identity level of these genomic fragments that were nearly identical between sister species and obtained two distributions, one for the identity percentage and one for the length. We pooled the values of the comparisons between *P. olsonii* and *P. chrysogenum* on the one hand and *P. olsonii* and *P. rubens* on the other hand, as *P. chrysogenum* and *P. rubens* are so closely related that they were previously considered a single species. We considered the maximal identity percentage as the identity threshold for identifying HGTs. Because the longest sequence in the distribution was two times longer than the second longest one, we used the second maximal value of sequence length as the length threshold for identifying HGTs. These two threshold values for length and identity did not have to come from the same sequence and were considered independently. When comparing *P. salamii* and *P. nalgiovense* genomes, genomic regions longer and more similar than the two threshold cut-off values were therefore unexpected given their phylogenetic distance. The thresholds were highly conservative, to avoid false positives as much as possible, as we took the most extreme values of the distributions.

We then compared the genomes of each *P. nalgiovense* strain to each *P. salamii* strain pairwise with the same method as above and identified sequences shared between strains of the two species meeting the length and identity criteria, which we considered as putative horizontally transferred regions (HTRs). The sequences of the putative HTRs retrieved from the different genomes were mapped against each other using the “map to reference” algorithm in Geneious v9.1.8 (Biomatters Ltd., Auckland, New Zealand) and a consensus sequence was kept when they overlapped to obtain a set of reference HTRs.

To assess whether horizontal gene transfers were more frequent between *P. nalgiovense* and *P. salamii* than between two random pairs of *Penicillium* species, we compared 68 genomes from 54 species (four *Aspergillus* spp., *Monascus purpureus* and 49 other *Penicillium* spp.), using two different genomes for each species whenever possible (Table S1-S2). One genome of each of *P. biforme* and *P. egyptiacum,* and five genomes of *P. olsonii* were assembled from Illumina HiSeq reads for this analysis, with the method described above (Table S1). The other genomes were retrieved from public databases (Table S2). We compared the genomes in a pairwise fashion to identify HTRs using the same method as above. To obtain the phylogenetic distances between species pairs, we kept a single genome for each of the 54 species (Table S2), and we used the available coding sequences associated with the public sequences or detected the genes with EuGene v4.2a trained for *Penicillium* (Cheeseman et al., 2014; Foissac et al., 2008; Sallet et al., 2019) following the same procedure as for the *P. salamii* and *P. nalgiovense* genomes. To identify orthologs, we compared their protein sequences with blastp v2.6.0+ (Sayers et al., 2019) and grouped them by similarity using orthAgogue (Ekseth et al., 2014), choosing the markov clustering inflation parameter values that maximized the number of clusters containing 54 sequences (i.e one gene per species, I=2). We aligned the gene nucleotide sequences of single-copy orthologs using translatorX v1.1 and default parameters (alignment of the translated sequences with Muscle and cleaning with GBlocks and default parameters; Abascal et al., 2010). We then built a phylogenetic tree for each ortholog with RAxML v8.2.11, using the GTRGAMMAI model, starting from 20 independent trees and evaluated through 200 bootstraps (Stamatakis, 2014). We used these gene trees as input to build a species tree with Astral v5.6.3. (Zhang et al., 2018). We used the function cophenetic in the package *ape* (Paradis et al., 2004) run on R v3.6.2 (R Core Team, 2019) to extract the distance between species pairs. We plotted the cumulative lengths of the putative HTRs identified between species pairs as a function of the phylogenetic distance between species with the packages *ggplot* v3.3.0 and *ggrepel* v0.8.2 on R v3.6.2 (R Core Team, 2019).

As transposable elements (TEs) could be involved in horizontal gene transfers (Schaack et al., 2010), we checked whether particular transposable elements were present in the putative HTRs identified, and in particular at their margins. For this purpose, we identified TE present in *P. salamii* and *P. nalgiovense* reference genomes (LCP06525 and LCP06099-FM193) with RepeatModeler v1.0.11 (Smit & Hubley, 2017b). This software identifies TE families based on high frequency patterns but also relationship between the different elements and compare them to the Dfam library (v3.0; Storer et al., 2021), allowing some of the families to be formerly associated with known TE families in the five species present in the library (human, mouse, zebrafish, fruit fly, nematode), like LINE, LTR-Copia. We analyzed the putative HTRs with two databases: one based on all the TE identified in the *P. salamii* and *P. nalgiovense* reference genomes, and one based only on repeats associated to known TE families. Given the absence of fungi reference in the Dfam library, it is highly probable that the repeats identified but not associated with any known family are indeed TE, simply not annotated, but we cannot entirely rule out the possibility of any other duplicated sequences. We therefore looked at the 76 genes identified in these regions, none of them could be annotated with our protocol. To take this small uncertainty into consideration, we either used the entire HTRs or masked all the TE present in the first database with RepeatMasker v4.0.7 (Smit & Hubley, 2017a) for analyses not specifically focusing on TEs and mentioned those HTRs as “repeat-masked”. When the second database was used, masked sequences of the putative HTRs were identified as “TE-masked”.

To identify functions that could be enriched in the putative HTRs shared by *P. nalgiovense* and *P. salamii* compared to the rest of the genome, and therefore be potentially beneficial in dry-cured meat, the common method would be to perform an enrichment test in gene ontology (GO) function. However, because most genes present in these regions could not be annotated with GO terms, we used COG functional categories (clusters of orthologous groups) from eggNOG (Huerta-Cepas et al., 2015, 2017) and performed comparisons through Fisher’s exact tests given the small number of functions considered; we tested unilaterally for the over-representation of a particular function in HTR gene sets. To do so, we used the function fisher.test() in the package *stats* in R v3.6.2 (R Core Team, 2019).

We then searched for the presence of the reference set of HTRs in all *P. nalgiovense* and *P. salamii* genomes using blastn v2.2.29+ (Sayers et al., 2019), as well as in the 68 other genomes mentioned above. We used blastn because we aimed to detect fragments resulting from very recent horizontal transfers, so with nearly identical nucleotide sequences. When hits were obtained with the reference HTRs, we checked the proportion of each putative HTR present in the genome, its percentage of identity with the reference HTR sequence, its position in the genome by looking at flanking regions and whether it was fragmented on different scaffolds. Because the fragmentation of HTRs on different scaffolds could be due to incomplete assemblies, we considered a HTR to be split only when the region was present on two different scaffolds or when the additional sequence separating the two fragments had a length at least 10% higher than the sequence separating the two fragments on the reference HTR sequence. We also checked if the genomic region separating HTR fragments was only constituted of repeated sequences, in which case they could be TE insertions within HTRs following their transfer. This procedure led us to identify 23 putative HTRs. However, some groups of HTRs clustered close to each other in the reference genomes, like HTR_04, HTR_05 and HTR_07 on Pnal_04 and Psal_05 or HTR_06 and HTR_08 on Psal_14 (Figure 2) and could therefore actually share a common transfer history. No criterion appeared optimal to decide whether some HTRs were really independent or should be merged. The important finding is the existence and total content of HTRs and not their precise number. We therefore analyzed separately the 23 HTRs, or for some figures the eight largest ones, but it should be kept in mind that there may have been fewer horizontal transfer events.

To illustrate the presence, length, and similarity of HTRs in genomes, we clustered the regions based on the proportion of the HTRs present in each genome, using Euclidean distance and the default complete linkage clustering method. We plotted heatmaps using the heatmap.2 function of the *gplots* v3.0.1.1 package (Warnes et al., 2019) in R (R Core Team, 2019).

**Phenotypic tests: mycelium growth, salt tolerance, lipolysis, proteolysis, color, and spore numbers**

We compared various phenotypic traits between species and strains from different environments: growth rate at different salt concentrations, spore production, lipolysis, and proteolysis abilities. We first studied *P. nalgiovense* and *P. salamii*, the most common species inoculated for dry-cured meat production, with their respective sister species *P. chrysogenum* and *P. olsonii*. However, we found shared horizontally transferred regions between these dry-cured meat species and the *P. biforme* complex, including the two clonal lineages *P. camemberti* var. *camemberti* and *P. camemberti* var. *caseifulvum* (Ropars et al., 2020), and these can also be, even if more rarely, inoculated in dry-cured meat. We therefore also analyzed phenotypes in *P. biforme*, that display genetic and phenotypic variability, in contrast to the cheese clonal *P. camemberti* varieties (Ropars et al., 2020), and compared them to phenotypes of their sister species *P. palitans.* For all the experiments below, we used all available *P. nalgiovense*, *P. salamii* and *P*. *biforme* strains that we could find in public collections, from non-food environments, and similar numbers of strains, chosen at random, from dry-cured meat and from their closely related outgroup species. The Table S1 gives the number and identity of strains used for each species, type of environment and experiment.

For assessing salt tolerance, we used seven *P. chrysogenum* strains, 14 *P. nalgiovense* strains, eight *P. olsonii* strains, 17 *P. salamii* strains, 13 *P. biforme* strains and five *P. palitans* strains. Because the salt content of dry-cured meat products usually ranges between 2% to 5% of dry weight and can be much higher near the surface (Aaslyng et al., 2014; Corral et al., 2013), we measured growth rates on malt extract agar medium with various salt concentrations: 2% (2g NaCl/100 mL), 10% (10g NaCl/100 mL) and 18% (18g NaCl/100 mL). Malt extract agar medium was prepared with 16 g malt extract powder (Difal), 16g agar dissolved in 800 mL ddH_2_O and adjusted to pH 5.7 which is in the pH range of dry-cured meat products (pH 4.5 – 6; Aaslyng et al., 2014). We poured 30 mL of medium into Petri dishes (150 mm diameter). For inoculation, *Penicillium* spore suspensions were calibrated by counting spores in a Malassez hemocytometer cell using six squares (0.2 mm, two repeats) under the microscope. Spore number estimates were highly correlated between repetitions (r² = 0.94, df = 63, p-value < 2.2e-16). We then diluted spores to a concentration of 6 x10^5^ spores.mL^-1^ and deposited 10 μL in the center of each Petri dish before incubating plates at room temperature. We recorded growth by taking photos under constant conditions in terms of light and position of Petri dishes, at days 8 and 17. We used the Image J package Fuji (Guzmán et al., 2014; Samson et al., 2004) to measure colony area on the pictures using Trainable Weka Segmentation v3.2.23 (Arganda-Carreras et al., 2017). The software was trained to recognize background, medium and colony areas on the picture. The area estimates in pixel units were transformed into square centimeters using the size of the Petri dish. In the last step of the salt tolerance experiment, we collected all fungal material from each Petri dish in an Eppendorf tube filled with 0.05 % Tween® 20 water, which was then diluted (dilution 1:100). Spore numbers were estimated from these solutions under a microscope (Nikon ECLIPSE TS100, 40X) with a Malassez hemocytometer using the same protocol as above after 17 days of growth.

The lipolytic and proteolytic activities were tested with the same *Penicillium* strains as salt tolerance tests. Lysis activities were measured as follows: a suspension adjusted to 2,500 spores for each strain was inoculated on the top of a test tube containing tributyrin (triglyceride) agar for lipolytic activity measurements (10 mL.L^-1^, ACROS Organics, Belgium) or semi-skimmed milk for proteolytic activity measurements (40 g.L^-1^, from large retailers). Lipolytic and proteolytic activities were estimated on days 7, 14, 21 and 28 by determining extent of compound degradation as the media changes from opaque to translucent. Experiments were divided into three batches (N=26, 30 and 8 strains respectively for each test period).

To evaluate fungal colony color, spore suspensions of all available *Penicillium* strains (n=78; *P. chrysogenum* n=7, *P. nalgiovense* n=27, *P. olsonii* n=8, *P. salamii* n=22, *P. biforme* n=9 and *P. palitans* n=5; Table S1) were inoculated onto malt extract agar medium (70164 Sigma-Aldrich) and calibrated photos were taken after 18 days growth at 25°C using a ScanStation 100 (Interscience, Saint Nom, France). Images were analyzed using Image J packageFuji (Guzmán et al., 2014; Samson et al., 2004) to record the color decomposition among red, green and blue colors (RGB system). Levels of red, green, and blue are represented by a range of integers from 0 to 255 (256 levels for each color). White is 255,255,255 and black 0,0,0. Color was measured in three different circles on Petri dishes, each circle being of 0.5 cm diameter, located respectively in the center, edge of the colony and midway between these two points. RGB color values were averaged among pixels within each circle. As colony whiteness potentially represents a trait under selection by humans, we computed and analyzed the degree of whiteness by computing the distance to white, *i.e.,* the Euclidean distance in the 3-D orthogonal coordinates composed of red, blue and green values. We tested whether the colors of *P. biforme, P. nalgiovense* and *P*. *salamii* were closer to white than that of their sister species not used for dry-cured meat production, and whether, within each species, dry-cured meat strains were whiter than non-dry-cured meat strains.

**Mycotoxins and penicillin production**

For mycotoxin production, we grew fungal cultures in 24-well sterile microplates containing 2 mL agar medium per well. For each strain, 1 µL of a calibrated spore suspension (10^6^ spores.mL^-1^) prepared from a 7-day culture was inoculated in the center of yeast extract sucrose (YES) agar medium buffered at pH 4.5 with phosphate-citrate buffer and characterized by a high C/N ratio to favor mycotoxin production as already described by Gillot et al (Gillot et al., 2017). Six replicates per strain were performed, three for mycotoxin analyses and three for fungal dry-weight measurements. In the latter case, growth was performed on cellophane disks to collect fungal mycelium. The plates were incubated at 25°C in the dark for seven days and then stored at -20°C until mycotoxin analysis.

For mycotoxin extractions, an optimized “high-throughput” extraction method based on the one described by Gillot et al (Gillot et al., 2017) was used. Briefly, 2g-aliquots (the entire YES culture obtained from a well) were homogenized after thawing samples with a sterile flat spatula then 12.5 mL of acetonitrile (ACN) supplemented with 0.1% formic acid (v/v) was added, samples were vortexed for 30 sec followed by 15 min sonication. Then, extracts were again vortexed before being centrifuged for 10 min at 5000g at 4°C. The supernatants were directly collected and filtered through 0.45 µm PTFE membrane filters (GE Healthcare Life Sciences, UK) into amber vials and stored at -20°C until analyses.

Mycotoxin detection and quantification were performed using an Agilent 6530 Accurate-Mass Quadropole Time-of-Flight (Q-TOF) LC/MS system equipped with a Binary pump 1260 and degasser, well plate autosampler set to 10°C and a thermostated column compartment. Filtered 2µL aliquots were injected into a ZORBAX Extend C-18 column (2.1x50 mm and 1.8 µm, 600 bar) maintained at 35°C with a flow rate set to 0.3 mL.min^-1^. The mobile phase A contained milli-Q water + 0.1% formic acid (v/v) and 0.1% ammonium formate (v/v) while mobile phase B was ACN + 0.1% formic acid. Mobile phase B was maintained at 10% for 4 min followed by a gradient from 10 to 100% for 18 min. Then, mobile phase B was maintained at 100% for 5 min before a 5-min post-time. Samples were ionized in both positive (ESI+) and negative (ESI-) electrospray ionization modes in the mass spectrometer with the following parameters: capillary voltage 4 kV, source temperature 325°C, nebulizer pressure 50 psig, drying gas 12 L.min^-1^, ion range 100-1000 m/z. Target extrolite characteristics used for quantifications are given in Table S8 and included commercially available extrolites produced by *Penicillium* species: andrastin A, ermefortins A & B, (iso)-fumigaclavin A, meleagrin, mycophenolic acid, ochratoxin A, penicillic acid, penicillin G, penitrem A, roquefortin C, sterigmatocystin. Andrastin A, ermefortins A & B and (iso)-fumigaclavin A standards were obtained from Bioviotica (Goettingen, Germany), penitrem A from Cfm Oskar Tropitzsch (Marktredwitz, Germany), while all others were from Sigma-Aldrich (St Louis, MO, USA). All stock solutions were prepared in dimethyl sulfoxide (DMSO) at 1 mg.mL^-1^ in amber vials.

For these analyses, metabolite identification was performed using both the mean retention time ± 1 min and the corresponding ions listed in Table S8. We used a matrix matched calibration curve (R^2^ >0.99 for all extrolites except 2 >0.96) for reliable mycotoxin quantification with final concentrations ranging from 1 to 10000 ng.mL^-1^ according to the target metabolite and method performance was carried out as previously described by Gillot et al (Gillot et al., 2017). Mycotoxin concentrations were calculated from the equation y = mx+b, as determined by weighted (1/x^2^) linear regression of the matrix-matched calibration data and correlated to calculated fungal growth areas. Specific mycotoxin production was expressed as ng per g of fungal dry weight.

**Supplementary data:**

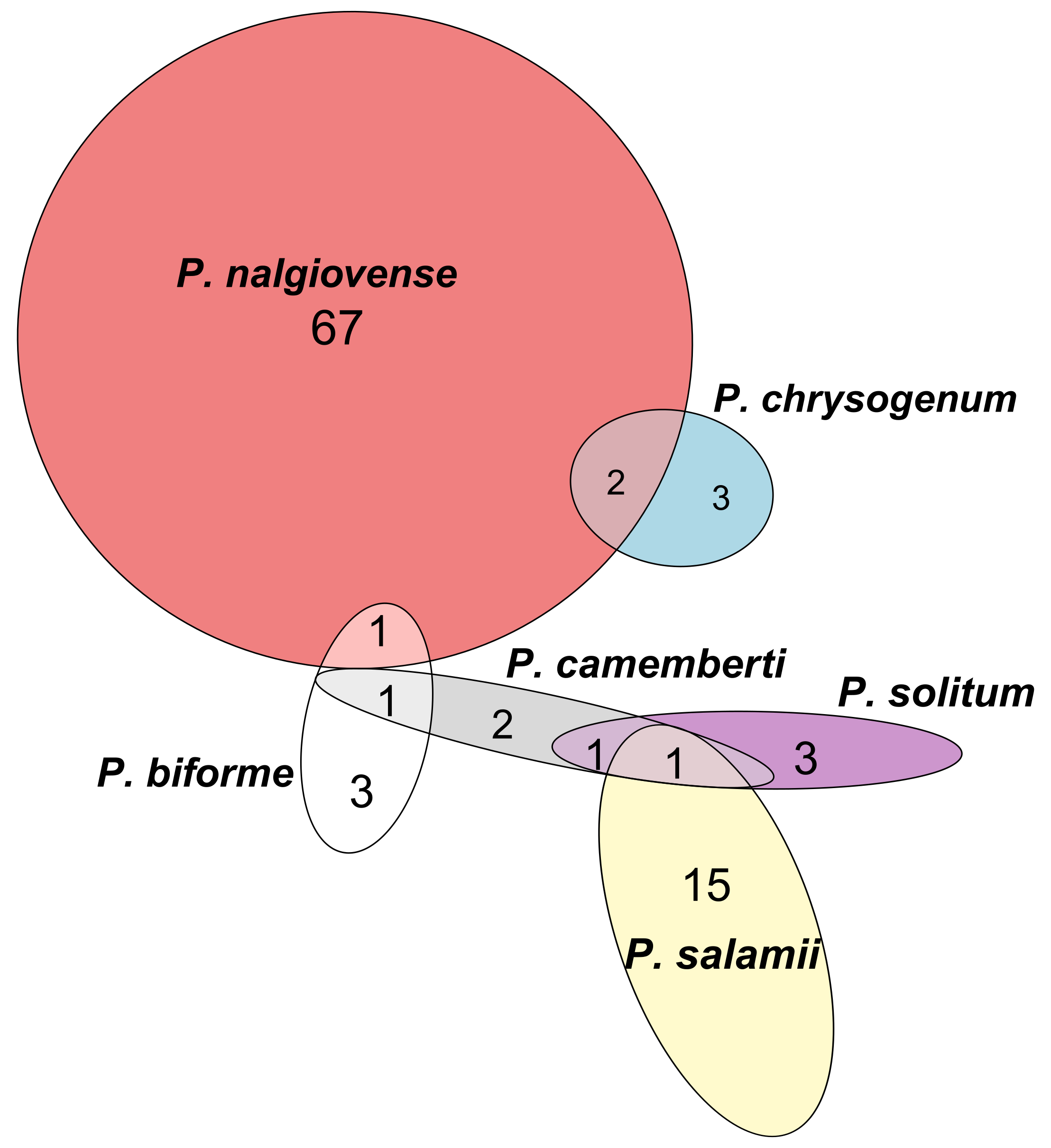

**Figure S1. Venn diagram showing the number of strains isolated belonging to the five most abundant *Penicillium* species found in dry-cured meat.** The overlaps between ellipses represent the number of strains isolated together with another species from the same dry-cured meat sample.

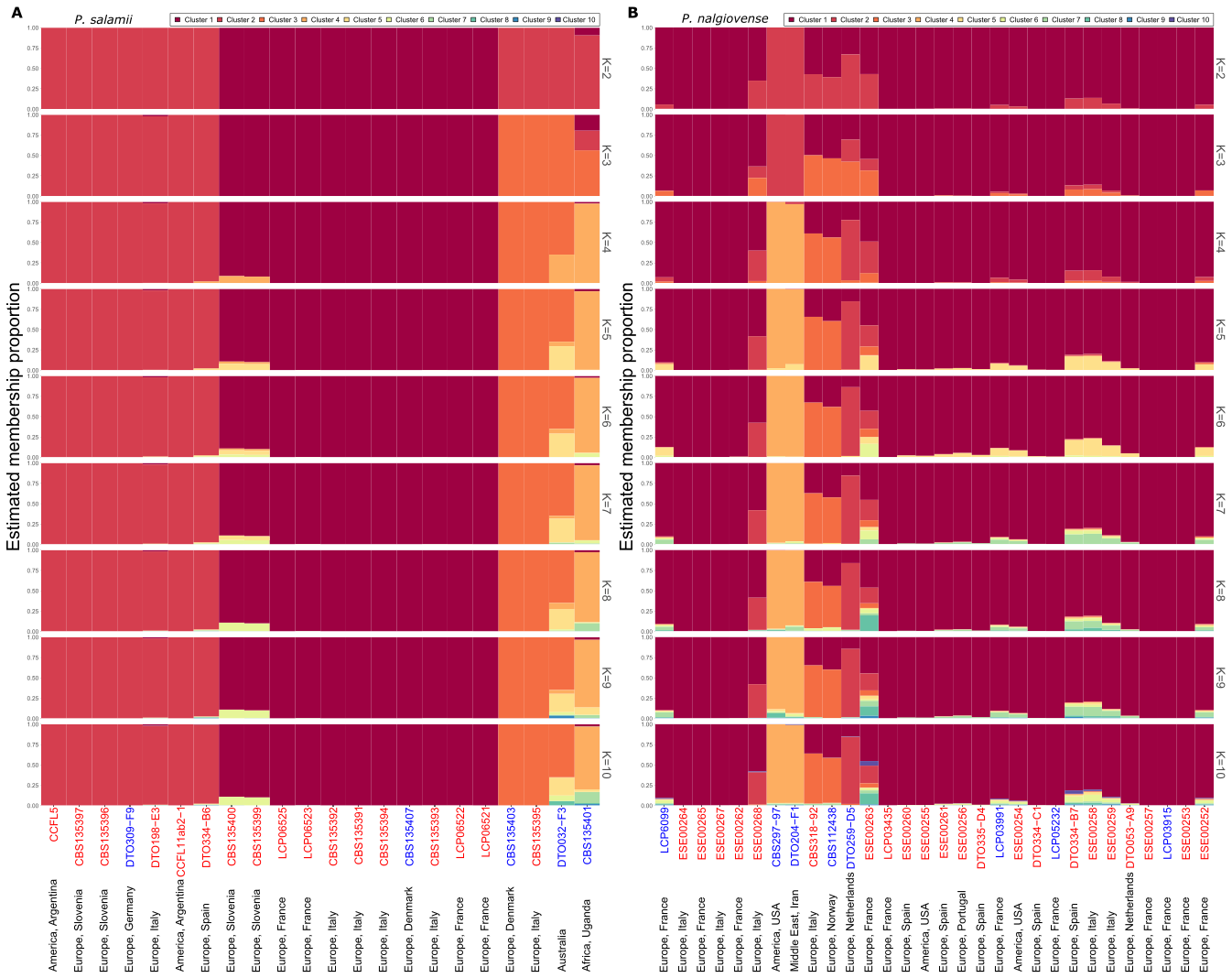

**Figure S2. Population structure inferred by STRUCTURE in *Penicillium* *salamii*** (A) **and *P*. *nalgiovense*** (B). Each bar represents an individual, with its estimated proportions of genetic information assigned to different clusters, represented by colors, for numbers of clusters ranging from K = 2 to K = 10. The environment of isolation is indicated at the bottom, with strains names in red when collected on dry-cured meat and in blue when collected in other environments. Strains are ordered following the populations trees from Fig. 1.

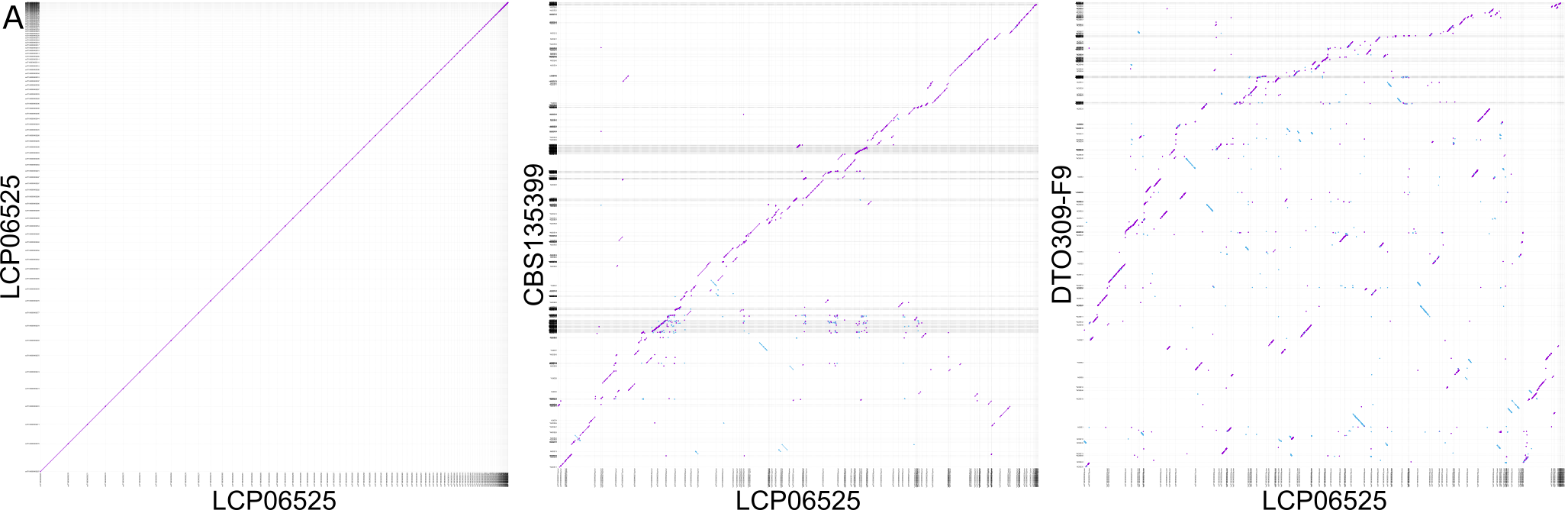

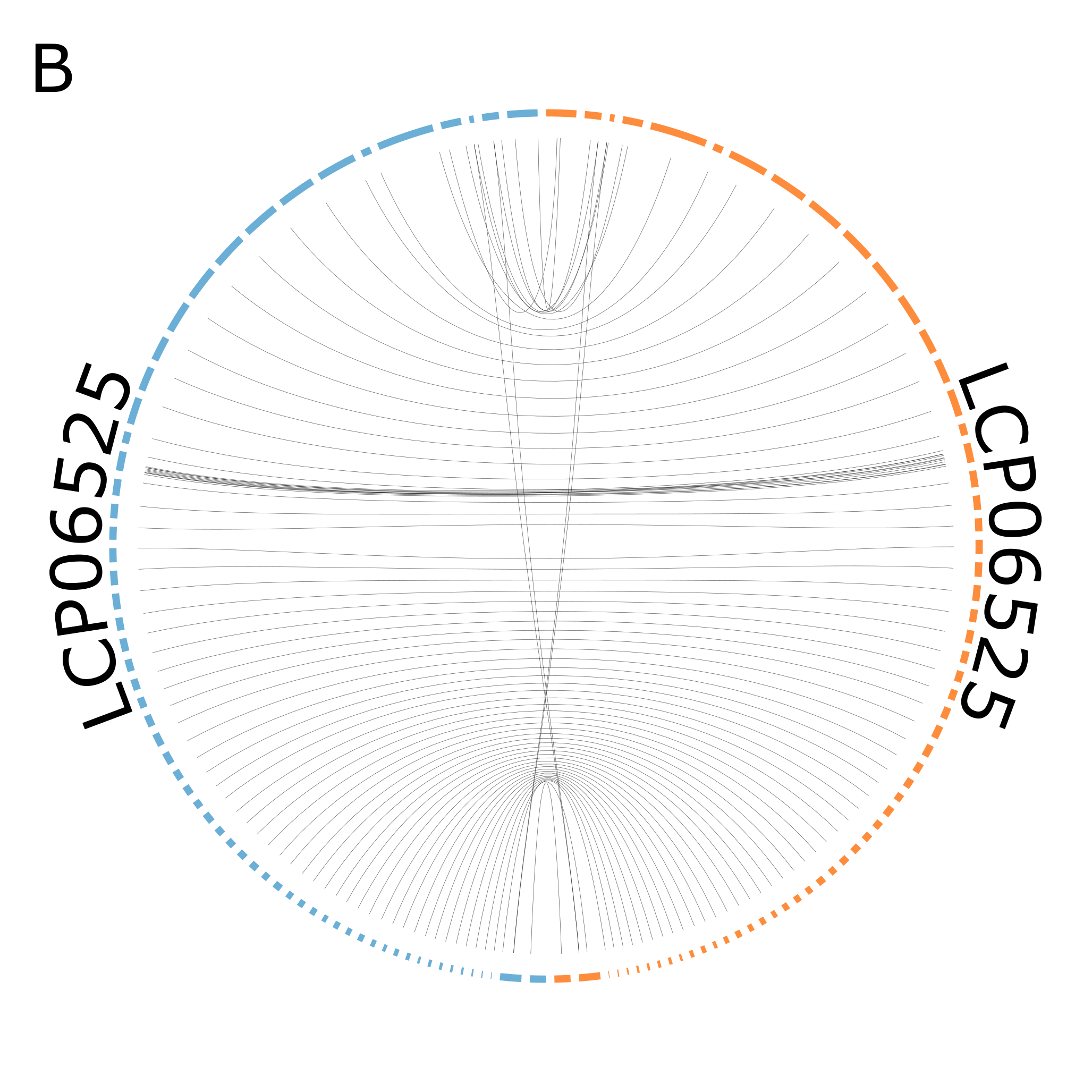

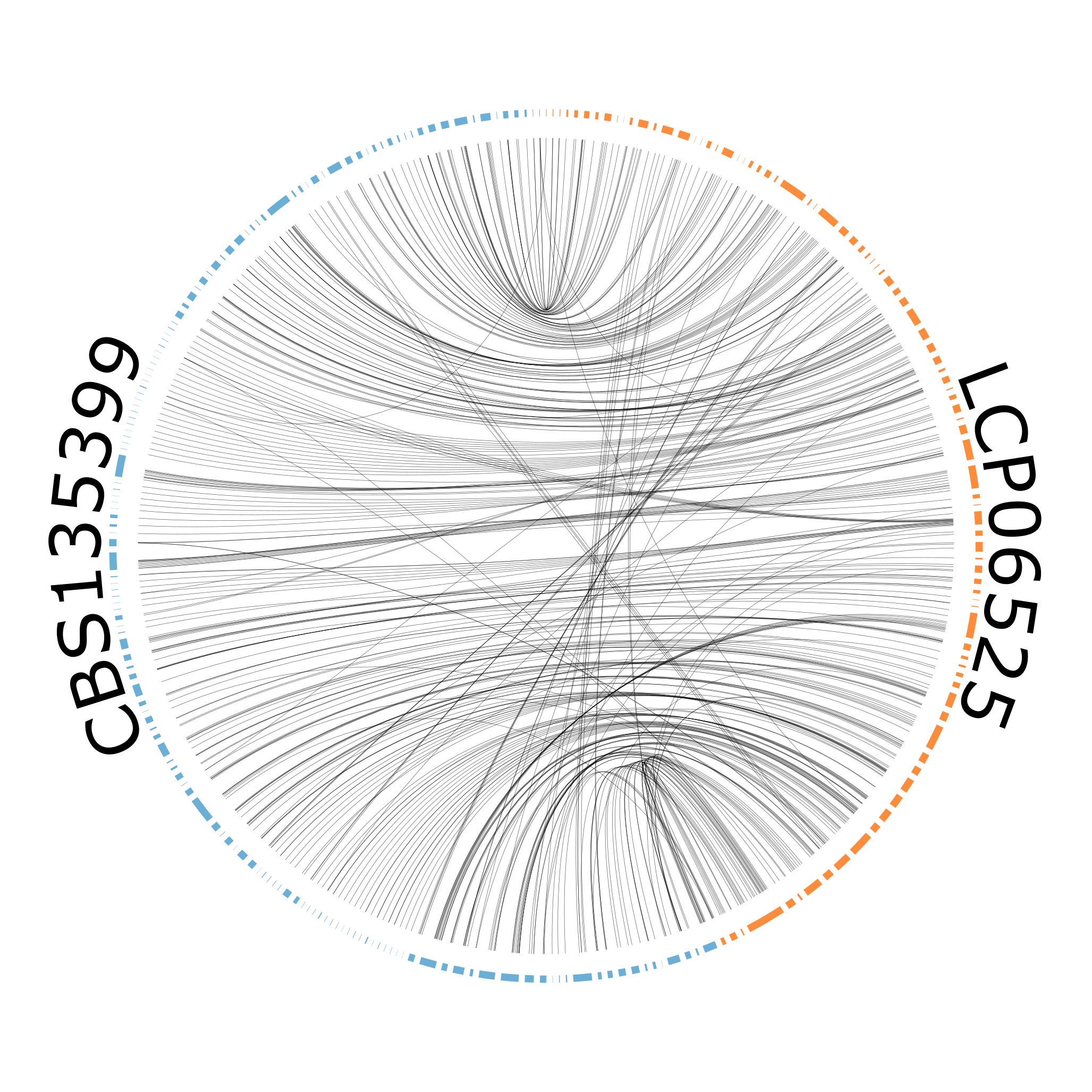

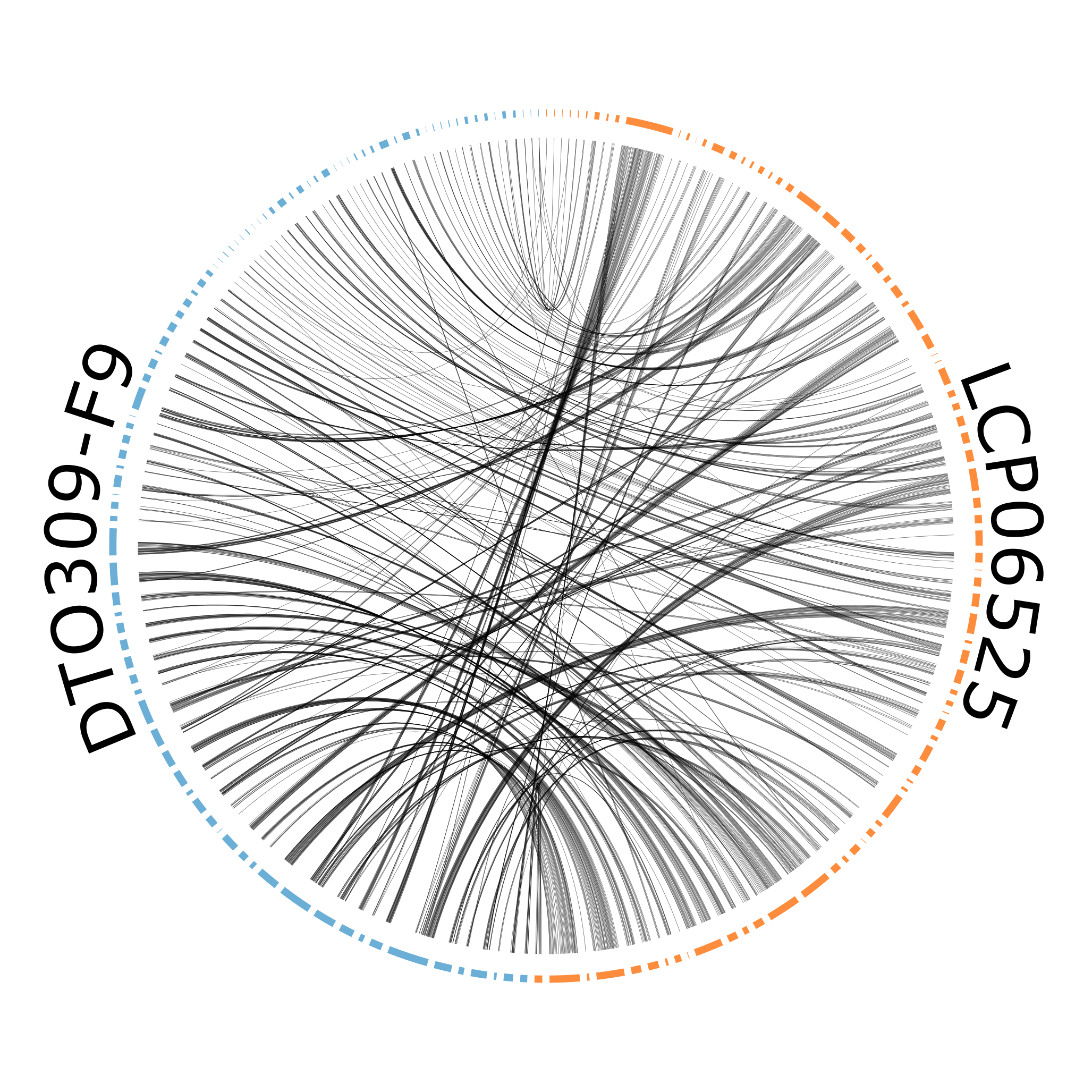

**Figure S3. Examples of pairwise comparisons of the genomic architectures among *Penicillium salamii* strains through mummerplots (A) and circos (B)**. In mummerplots, regions collinear between two strains are depicted in purple and in the direction bottom-left to upper-right, while inversions are shown in blue in an orthogonal direction. Two identical genomes should therefore present a clear purple diagonal (example on the left), while many rearrangements tend to blur the pattern (example on the right). Based on these plots, we qualitatively identified six groups. The example in the middle shows two genomes belonging to the same group. In circos plots, the scaffolds are ordered along a circle, with the reference LCP06525 in orange on the right, and the other strain in blue (LCP06525, CBS135399 and DTO309-F9 respectively). The lines in the middle correspond to regions with high similarity between the two genomes. Scaffolds were ordered following the mummerplots.

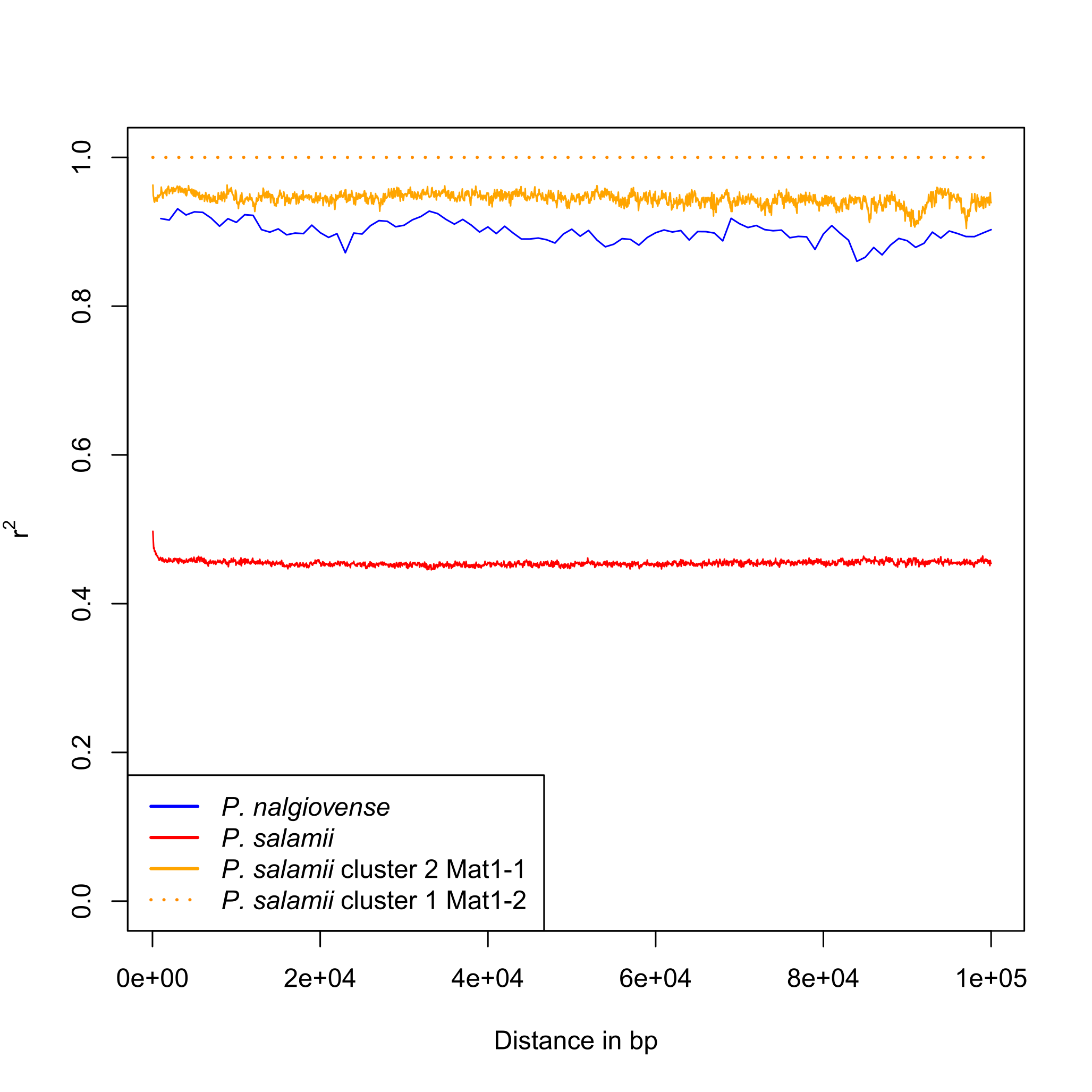

**Figure S4. Linkage disequilibrium (LD) decay in *Penicillium nalgiovense*, *P. salamii* as a whole and within the two main *P. salamii* clades (C).**

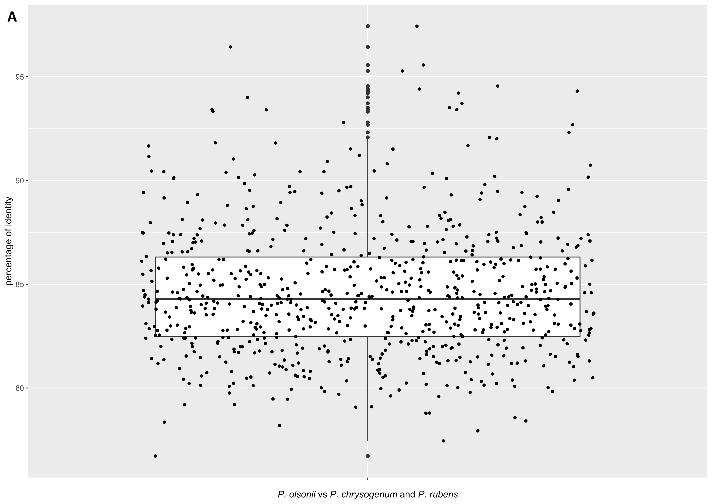

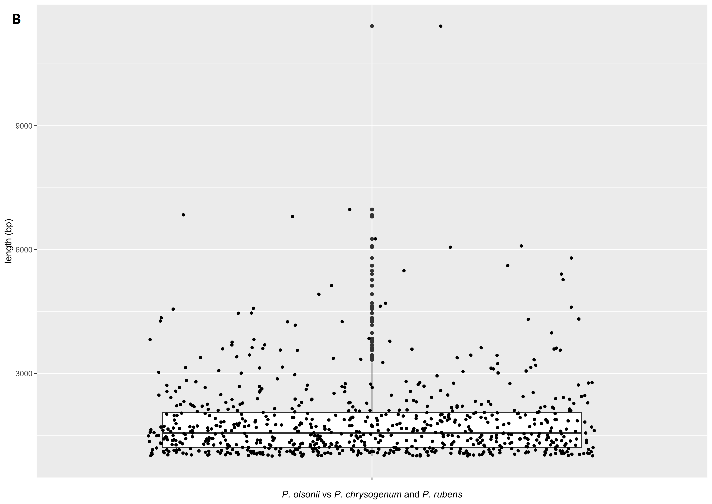

**Figure S5. Distribution of the percentage of identity (A) and length (B) of the sequences detected as similar between *Penicillium olsonii* and *P. chrysogenum* / *P. rubens*, sister species of *P. salamii* and *P. nalgiovense* respectively, used to establish threshold for the detection of horizontally transferred regions.** The sequences were considered similar when sharing at least one block of 65 identical consecutive bases; the blocks could be merged if not separated by more than 90 bp (default NUCmer parameters). Any sequence more similar and longer was not expected to occur by chance and was considered as potentially horizontally transferred.

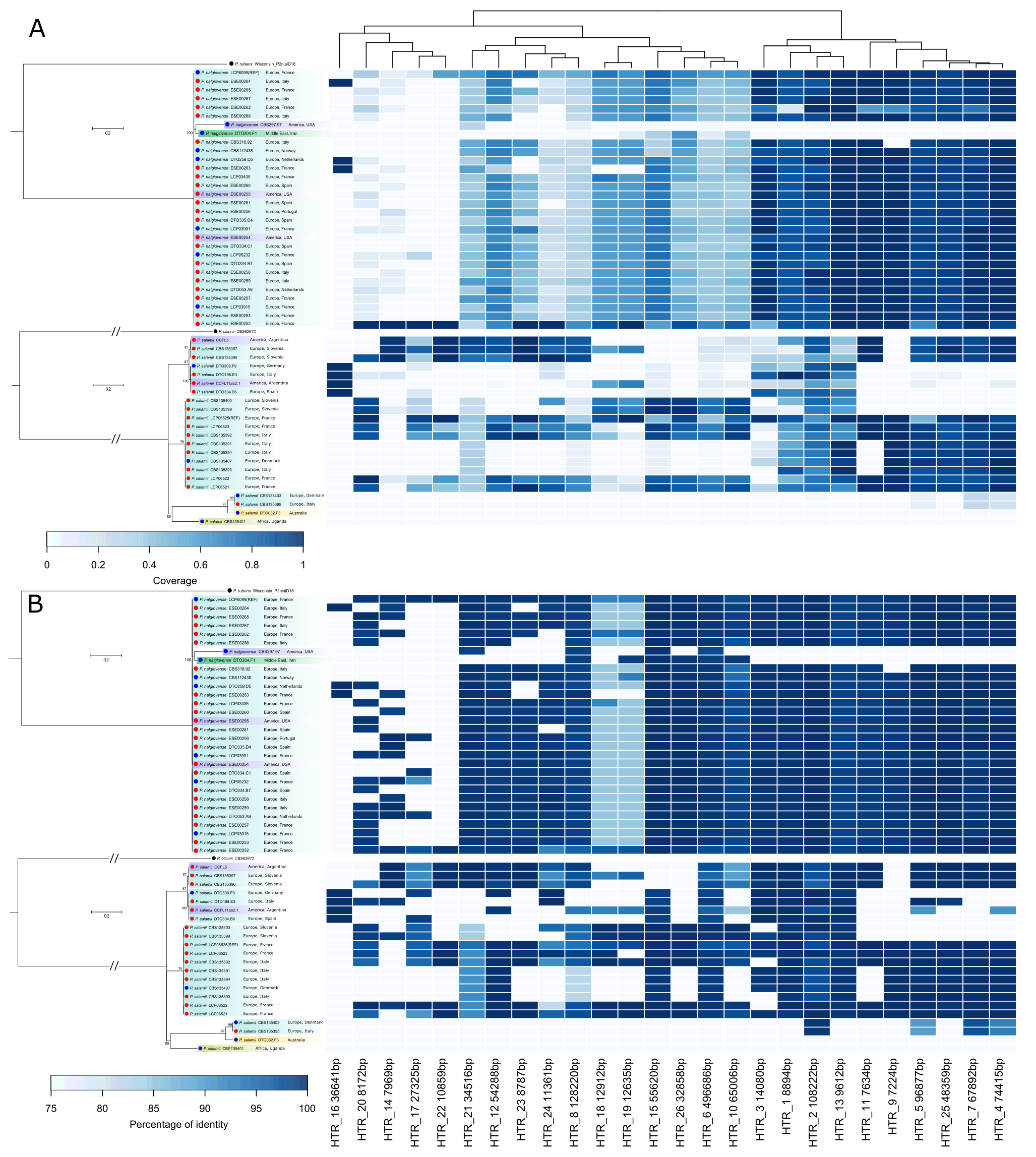

**Figure S6.** **Presence (dark blue rectangle) or absence (lightest blue rectangle) of putative horizontally-transferred regions (HTRs) without sequence masking, in *Penicillium nalgiovense* and *P. salamii* strains.** The blue darkness indicates the length percentage of HTRs present in each strain compared to the length of the reference consensus sequence (A), or the percentage of identity between the sequence in the focal strain and the reference consensus sequence (B). The regions are ordered based on their similarity in coverage.

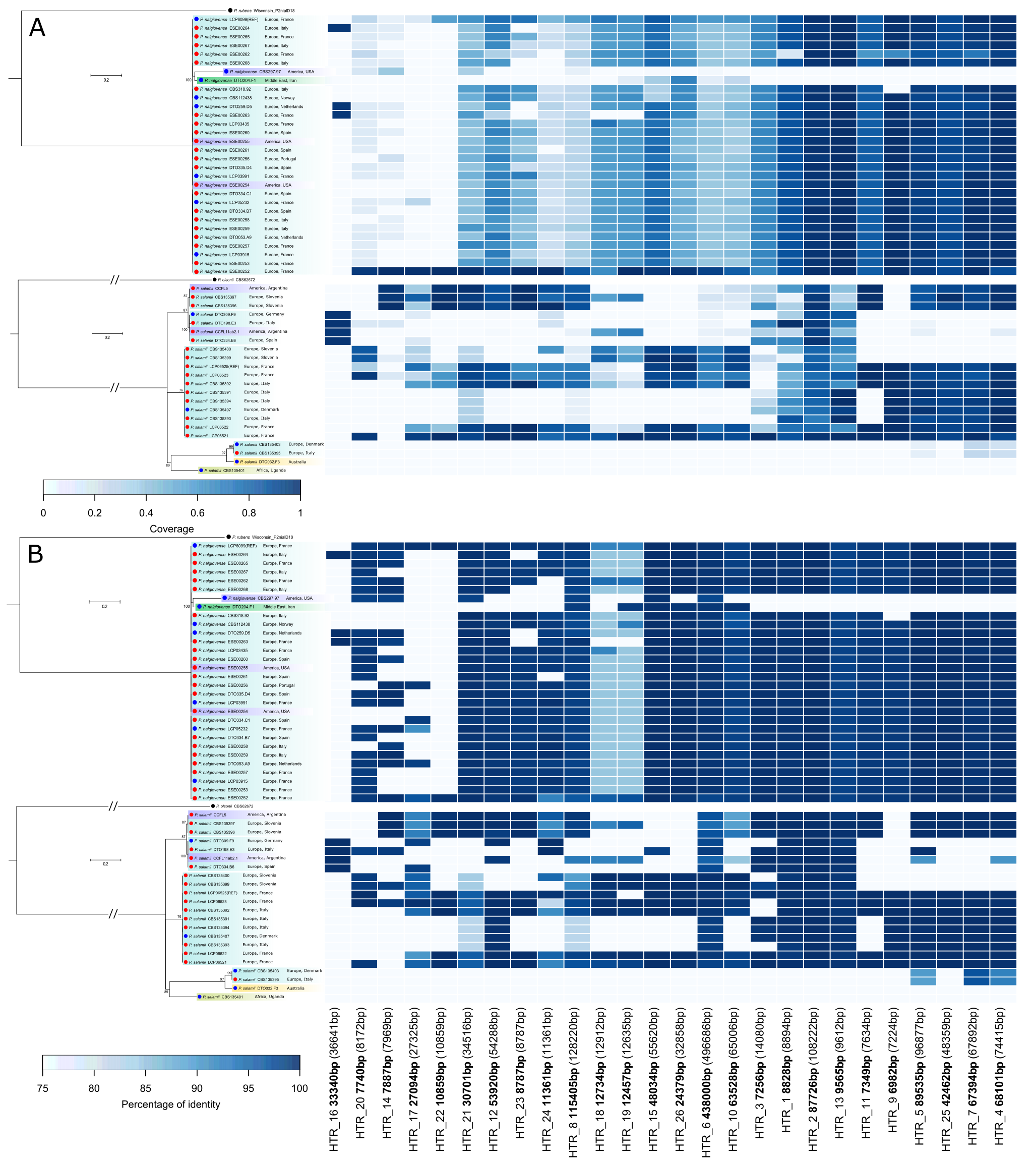

**Figure S7. Presence (dark blue rectangle) or absence (lightest blue rectangle) of putative horizontally transferred regions (HTRs) with transposable elements masked (“TE-masked” dataset), in *Penicillium nalgiovense* and *P. salamii* strains.** The blue darkness indicates the length percentage of HTRs present in each strain compared to the length of the reference consensus sequence (A), or the percentage of identity between the sequence in the focal strain and the reference consensus sequence (B). The lengths between brackets at the bottom correspond to the total lengths of the regions (without masking). The regions are ordered based on their similarity in coverage without masking (Fig. S6).

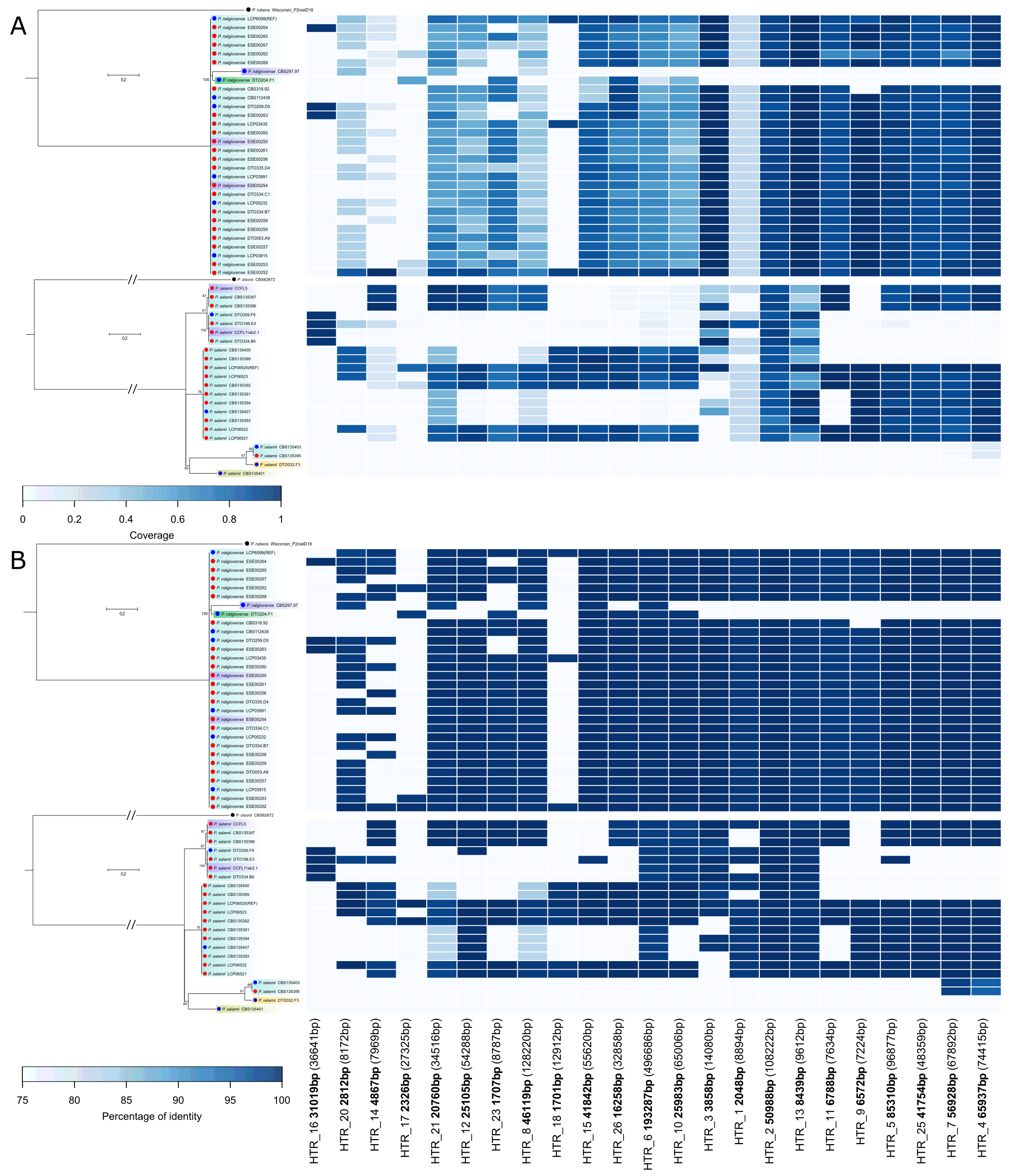

**Figure S8. Presence (dark blue rectangles) or absence (light blue rectangles) of putative horizontally-transferred regions (HTRs) in *Penicillium nalgiovense* and *P. salamii* strains.** Transposable elements were masked in the HTRs reference (“repeat-masked” scenario). The blue darkness indicating the length percentage of HTRs present in each strain compared to the length of the reference consensus region (A), or the percentage of identity between the sequence in the focal strain and the reference consensus region (B). The regions are ordered based on their similarity in coverage without masking (Fig. S6). The lengths between brackets at the bottom correspond to the total lengths of the regions (without masking).

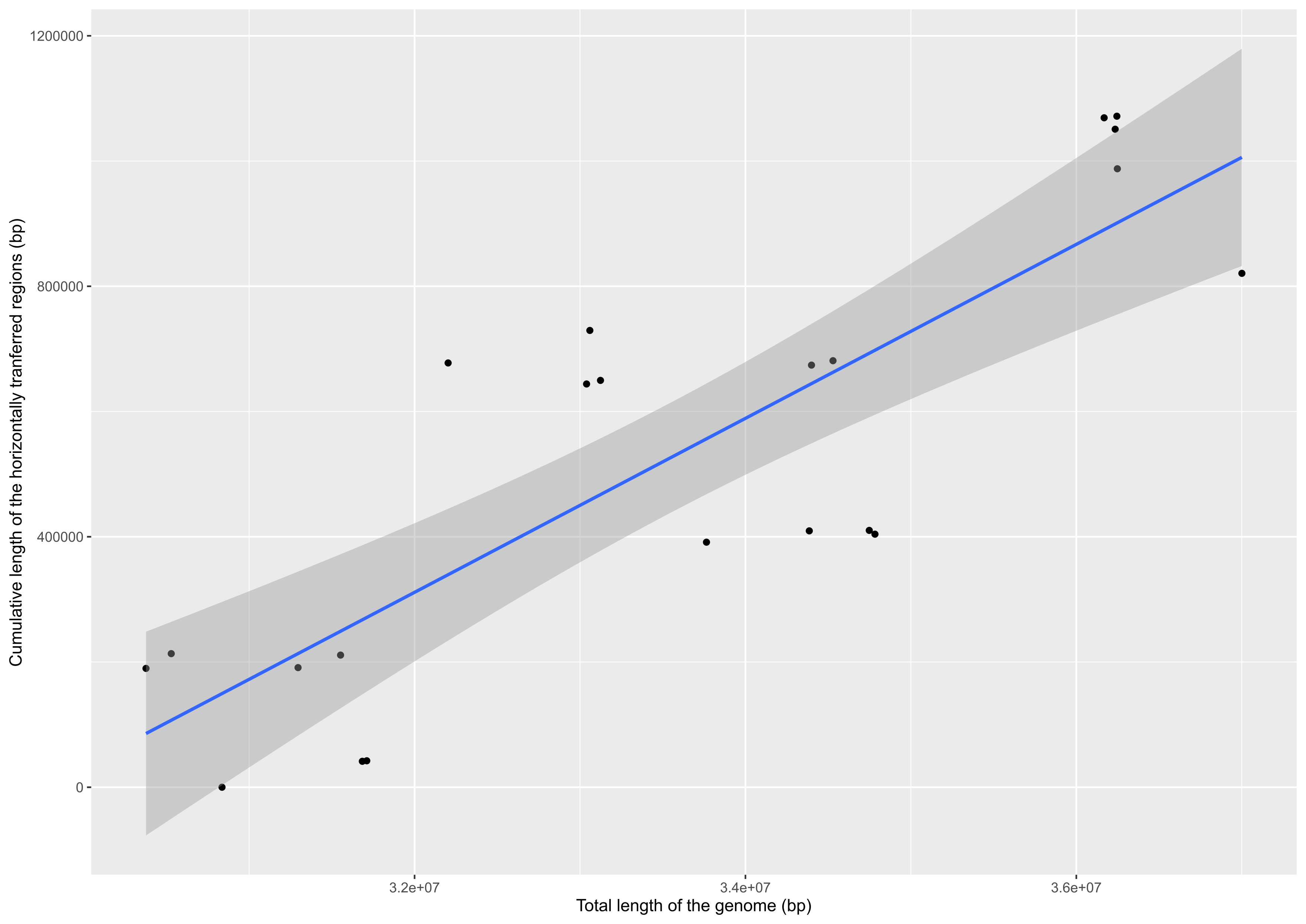

**Figure S9. Variation of the cumulative length of the horizontally transferred regions as a function of the total length of the genome across *Penicillium salamii* strains.** The blue line corresponds to the regression line, and the grey interval to the standard error.

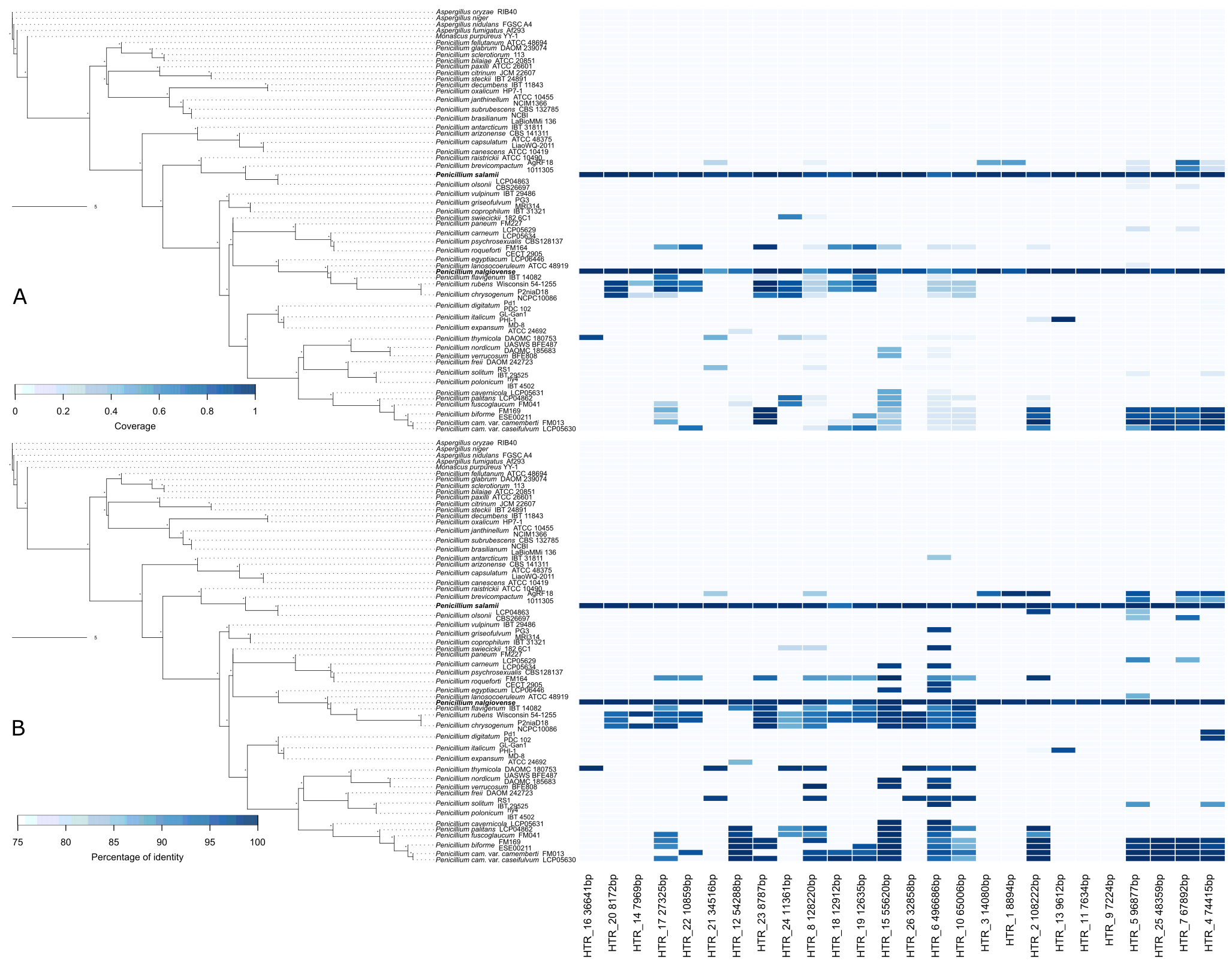

**Figure S10. Presence (dark blue rectangle) or absence (light blue rectangle) of putative horizontally transferred regions without sequence masking, in various *Aspergillus*, *Monascus* or *Penicillium* genomes.** The blue darkness indicates the length percentage of the horizontally transferred region present in each strain compared to the length of the reference consensus sequence (A), or the percentage of identity between the sequence in the focal strain and the reference consensus sequence (B).

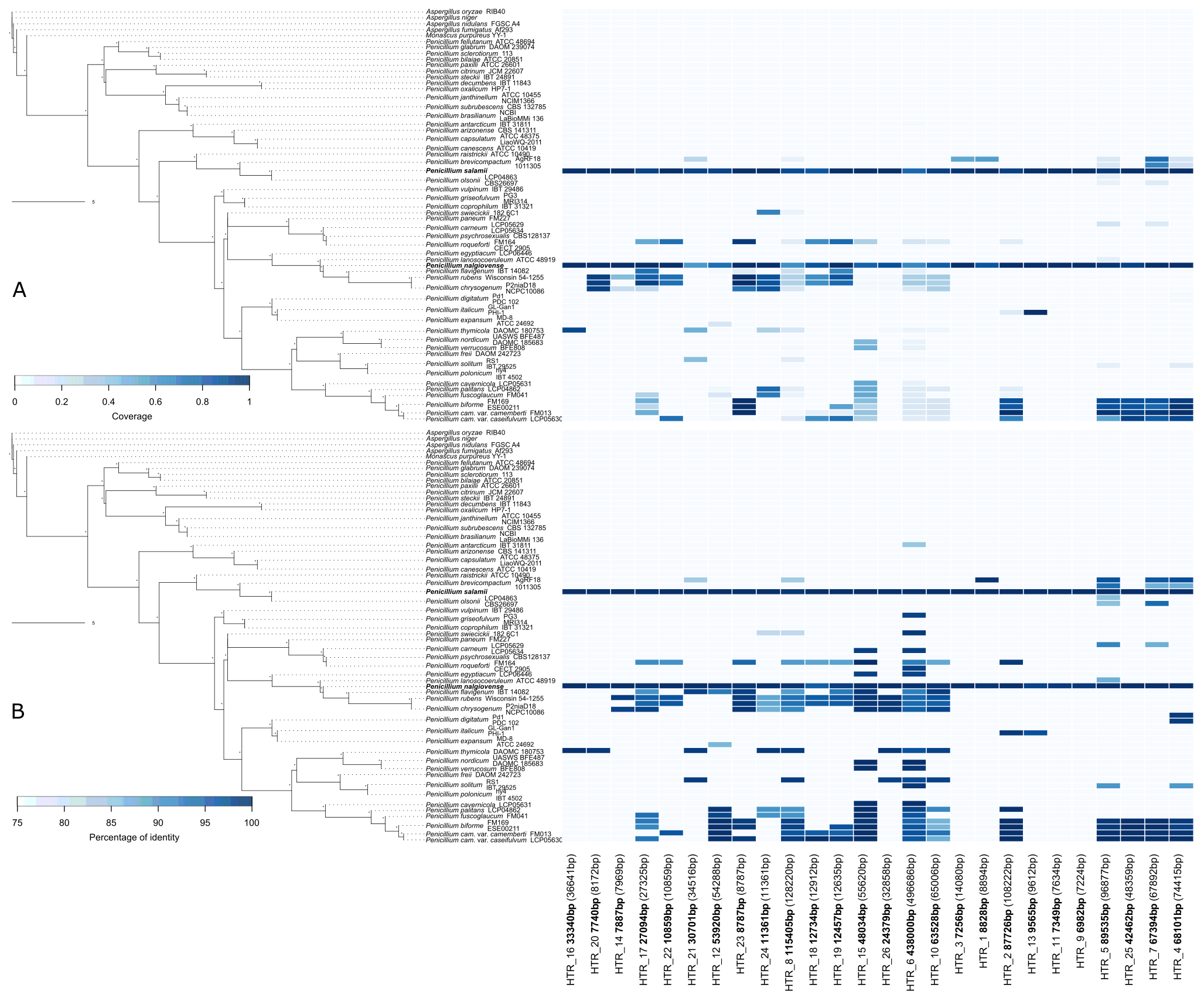

**Figure S11.** **Presence (dark blue rectangle) or absence (light blue rectangle) of putative horizontally transferred regions with transposable elements masked (“TE-masked” dataset), in various *Aspergillus*, *Monascus* or *Penicillium* genomes.** The blue darkness indicates the length percentage of the horizontally transferred region present in each strain compared to the length of the reference consensus sequence (A), or the percentage of identity between the sequence in the focal strain and the reference consensus sequence (B). The lengths between brackets correspond to the total lengths of the regions (without masking).

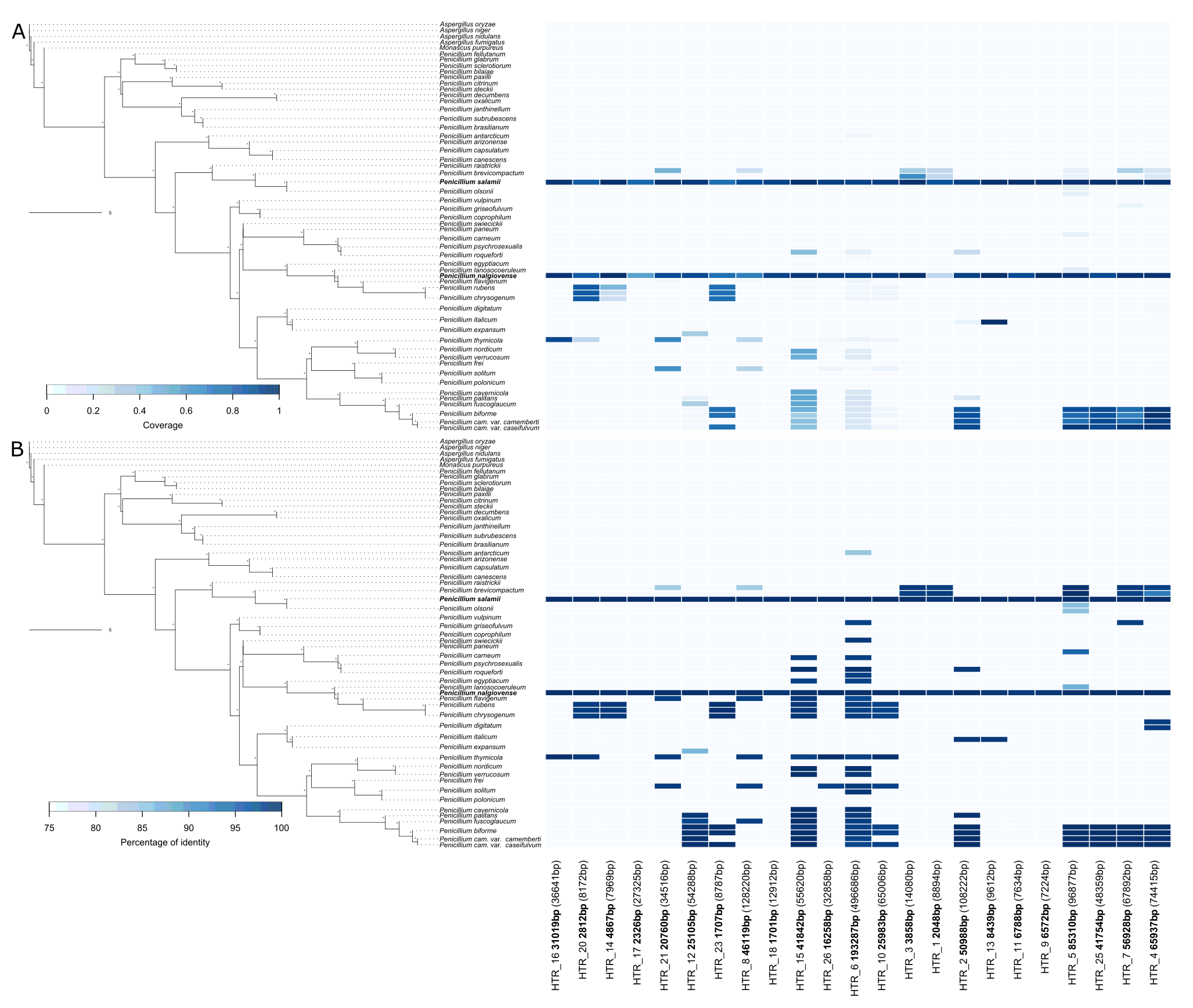

**Figure S12. Presence (dark blue rectangles) or absence (light blue rectangles) of putative horizontally-transferred regions (HTRs) in *Aspergillus*, *Monascus* and *Penicillium* strains.** Transposable elements were masked in the reference HTRs (“repeat-masked” scenario). The blue darkness indicating the length percentage of HTRs present in each strain compared to the length of the reference consensus region (A), or the percentage of identity between the sequence in the focal strain and the reference consensus region (B). The lengths between brackets correspond to the total lengths of the regions (without masking). The unrooted phylogenetic tree was built from 1102 single-copy genes with ASTRAL and the distance in the internal branches is expressed in coalescent units. Stars on the right end of each branch correspond to a full support of the relationship between the four leaves surrounding this branch.

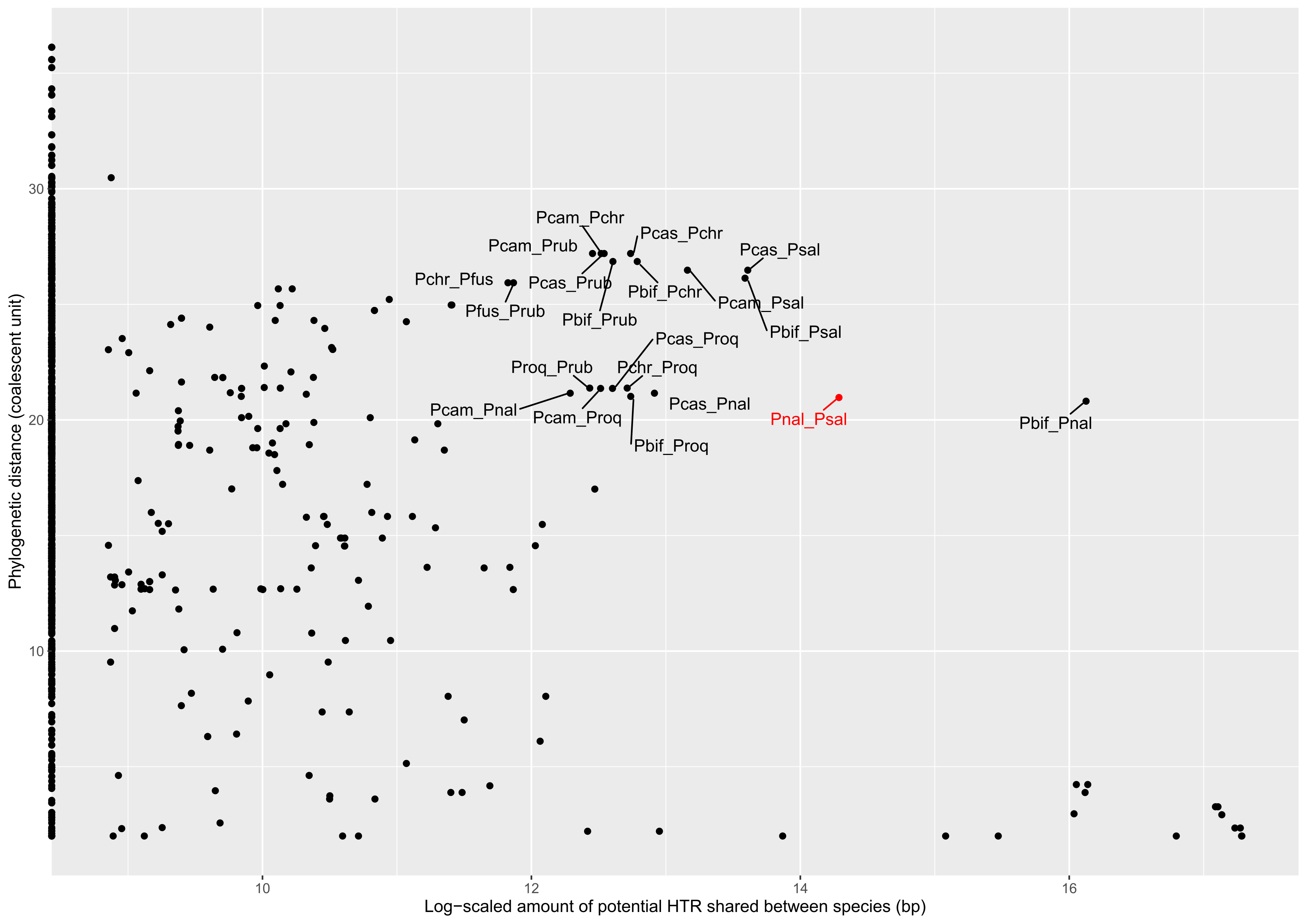

**Figure S13. Relation between the phylogenetic distance and the total length of putative horizontally transferred regions between pairs of Aspergillaceae species.** Pbif: *Penicillium biforme*, Pcam: *P. camemberti*, Pcas: *P. caseifulvum*, Pchr: *P. chrysogenum*, Pfus: *P. fuscoglaucum*, Pnal: *P. nalgiovense*, Proq: *P. roqueforti*, Prub: *P. rubens*, Psal: *P. salamii*.

**Table S1. *Penicillium* strains used in this study**; species, collection, sample number, substrate and country of sampling, and use in the study (genome sequencing, phenotypic tests). For the strains sequenced: number of contigs, total assembly length, GC content and N50 (sequence length of the shortest contig at 50% of the total genome length). ESE: Ecology, Systematic and Evolution Laboratory, Paris Saclay University, MNHN: Museum of Natural History, CBS: CBS-KNAW Culture Collections. ✓: salt tolerance test, protein metabolism test, lipid metabolism test; c: color test; m: mycotoxins test.

| Species | Collection | Sample number | Other collection names | Substrate of sampling | Country | Phenotypic test | Genome sequencing | Number of contigs | Total length (bp) | GC (%) | N50 (bp) | Genbank accession number |
| --- | --- | --- | --- | --- | --- | --- | --- | --- | --- | --- | --- | --- |
| *P. biforme* | ESE | PN025-2 |  | Sausage | France | ✓, c |  |  |  |  |  |  |
| *P. biforme* | ESE | ESE00207 |  | Sausage | France | ✓, c |  |  |  |  |  |  |
| *P. biforme* | ESE | ESE00151 |  | Sausage | n.a. | ✓, c |  |  |  |  |  |  |
| *P. biforme* | ESE | ESE00211 | PN023 | Sausage | France |  | Illumina | 503 | 35,896,733 | 47.55 | 205,634 | SRX9123590 |
| *P. biforme* | MNHN | LCP05496 |  | Air sampling | France | ✓ |  |  |  |  |  |  |
| *P. biforme* | MNHN | LCP05497 |  | Cheese | France | ✓ |  |  |  |  |  |  |
| *P. biforme* | MNHN | LCP05498 |  | Cheese | France | ✓ |  |  |  |  |  |  |
| *P. biforme* | MNHN | LCP05499 |  | Cheese | France | ✓, c |  |  |  |  |  |  |
| *P. biforme* | MNHN | LCP05500 |  | Food | France | ✓, c |  |  |  |  |  |  |
| *P. biforme* | MNHN | LCP05501 |  | Water | France | ✓ |  |  |  |  |  |  |
| *P. biforme* | MNHN | LCP05502 |  | Air sampling | France | ✓, c |  |  |  |  |  |  |
| *P. biforme* | MNHN | LCP05639 |  | Food | France | ✓, c |  |  |  |  |  |  |
| *P. biforme* | MNHN | LCP06101 |  | Cheese | France | ✓, c |  |  |  |  |  |  |
| *P. chrysogenum* | ESE | ESE00097 |  | Somport cheese (cow milk) | France | m |  |  |  |  |  |  |
| *P. chrysogenum* | ESE | ESE00106 |  | Ossau Iraty (raw sheep milk) | France | m |  |  |  |  |  |  |
| *P. chrysogenum* | ESE | ESE00128 |  | Sheep Tomme | France | m |  |  |  |  |  |  |
| *P. chrysogenum* | ESE | ESE00131 |  | Ossau Iraty (raw sheep milk) | France | m |  |  |  |  |  |  |
| *P. chrysogenum* | ESE | ESE00135 |  | Fontal cheese | Italy | m |  |  |  |  |  |  |
| *P. chrysogenum* | ESE | ESE00313 |  | Dry-cured meat Figavelle sausage | Kosovo | m |  |  |  |  |  |  |
| *P. chrysogenum* | ESE | ESE00352 |  | Molded chorizo | France | m |  |  |  |  |  |  |
| *P. chrysogenum* | ESE | ESE00361 |  | Edel salami | Germany | m |  |  |  |  |  |  |
| *P. chrysogenum* | ESE | ESE00434 |  | Dried sausage | Spain | m |  |  |  |  |  |  |
| *P. chrysogenum* | MNHN | LCP00456 |  | Seed | France | ✓, c, m |  |  |  |  |  |  |
| *P. chrysogenum* | MNHN | LCP00487 |  | Soil | France | ✓, c, m |  |  |  |  |  |  |
| *P. chrysogenum* | MNHN | LCP02616 |  | Unknown | The Bahamas | ✓, c, m |  |  |  |  |  |  |
| *P. chrysogenum* | MNHN | LCP03835 |  | Plant | Norway | ✓, c |  |  |  |  |  |  |
| *P. chrysogenum* | MNHN | LCP04047 |  | Water | Italy | ✓, c, m |  |  |  |  |  |  |
| *P. chrysogenum* | MNHN | LCP04082 |  | Food (not dry-cured meat) | Indonesia | ✓, c, m |  |  |  |  |  |  |
| *P. chrysogenum* | MNHN | LCP05427 |  | Refrigerator | France | ✓, c |  |  |  |  |  |  |
| *P. egyptiacum* | MNHN | LCP06446 |  | Soil | Egypt |  | Illumina | 909 | 32,674,410 | 48.88 | 1,200,179 | ERZ2879928 |
| *P. nalgiovense* | CBS | CBS109608 |  | Semi-hard white goat cheese | Greece | ✓, c |  |  |  |  |  |  |
| *P. nalgiovense* | CBS | CBS112438 |  | Ice | Norway | ✓, c | Illumina | 719 | 31,608,114 | 48.45 | 142,926 | ERZ2827838 |
| *P. nalgiovense* | CBS | CBS297.97 |  | Soil | USA | ✓, c, m | Illumina | 650 | 31,313,954 | 48.49 | 145,364 | ERZ2827884 |
| *P. nalgiovense* | CBS | CBS318.92 |  | Sausage | Italy (exported to Denmark) | c, m | Illumina | 674 | 31,474,098 | 48.45 | 149,785 | ERZ2827924 |
| *P. nalgiovense* | CBS | DTO053-A9 |  | Sausage | Netherlands | c, m | Illumina | 622 | 31,172,980 | 48.52 | 169,000 | ERZ2827962 |
| *P. nalgiovense* | CBS | DTO204-F1 |  | Soil | Iran | ✓, c, m | Illumina | 459 | 29,880,352 | 48.49 | 191,733 | ERZ2827989 |
| *P. nalgiovense* | CBS | DTO259-D5 |  | Dog bone | Netherlands | ✓, c, m | Illumina | 653 | 31,583,907 | 48.48 | 176,291 | ERZ2828028 |
| *P. nalgiovense* | CBS | DTO334-B7 |  | Fermented sausage | Spain | ✓, c, m | Illumina | 620 | 31,350,291 | 48.45 | 179,747 | ERZ2828065 |
| *P. nalgiovense* | CBS | DTO334-B8 |  | Fermented sausage | Spain | m |  |  |  |  |  |  |
| *P. nalgiovense* | CBS | DTO334-C1 |  | Fermented sausage | Spain | c | Illumina | 612 | 31,418,979 | 48.44 | 156,839 | ERZ2828108 |
| *P. nalgiovense* | CBS | DTO335-D4 |  | Fermented sausage | Spain | c, m | Illumina | 633 | 31,406,501 | 48.44 | 155,485 | ERZ2828150 |
| *P. nalgiovense* | MNHN | LCP03435 | PN134 | Sausage | France | c | Illumina | 512 | 31,422,740 | 48.46 | 210,949 | ERZ4121827 |
| *P. nalgiovense* | MNHN | LCP03915 | PN136 | Automotive interior lining | France | ✓, c | Illumina | 503 | 31,543,392 | 48.42 | 234,659 | ERZ2828750 |
| *P. nalgiovense* | MNHN | LCP03991 | PN115 | Seed | France | c | Illumina | 611 | 31,425,394 | 48.45 | 162,683 | ERZ2828774 |
| *P. nalgiovense* | MNHN | LCP05232 | PN140 | Air sample | France | ✓, c, m | Illumina | 624 | 31,334,022 | 48.48 | 154,267 | ERZ2828813 |
| *P. nalgiovense* | MNHN | LCP06099 (FM193) |  | Cheese | France |  | Illumina | 868 | 31,913,282 | 48.50 | 306,303 | GCA_000577395.2 |
| *P. nalgiovense* | ESE | ESE00252 | PN016 | Dry-cured meat | France |  | PacBio | 41 | 34,560,068 | 48.37 | 3,176,469 | ERZ2880275 |
| *P. nalgiovense* | ESE | ESE00253 | PN025 | Sausage | France | c | Illumina | 479 | 31,630,503 | 48.42 | 239,452 | ERZ2828191 |
| *P. nalgiovense* | ESE | ESE00254 | PN040 | Italy salami | America | ✓, c | Illumina | 618 | 31,381,282 | 48.45 | 179,781 | ERZ2828227 |
| *P. nalgiovense* | ESE | ESE00255 | PN044 | Dry-cured meat | America | c | Illumina | 645 | 31,057,889 | 48.59 | 166,006 | ERZ2828263 |
| *P. nalgiovense* | ESE | ESE00256 | PN054 | Chorizo sausage | Portugal | ✓, c | Illumina | 647 | 31,226,533 | 48.5 | 179,685 | ERZ2828305 |
| *P. nalgiovense* | ESE | ESE00257 | PN066 | Rocheblin - air dried sausage | France | c | Illumina | 476 | 31,547,343 | 48.42 | 261,417 | ERZ2828342 |
| *P. nalgiovense* | ESE | ESE00258 | PN072 | Sausage | Italy | c | Illumina | 619 | 31,097,831 | 48.54 | 165,240 | ERZ2828381 |
| *P. nalgiovense* | ESE | ESE00259 | PN075 | Sausage | Italy | ✓, c | Illumina | 647 | 31,359,185 | 48.49 | 161,865 | ERZ2828419 |
| *P. nalgiovense* | ESE | ESE00260 | PN076 | Sausage | Spain | ✓, c | Illumina | 628 | 31,236,137 | 48.53 | 175,999 | ERZ2828458 |
| *P. nalgiovense* | ESE | ESE00261 | PN077 | Sausage | Spain | c | Illumina | 627 | 31,120,043 | 48.5 | 165,263 | ERZ2828490 |
| *P. nalgiovense* | ESE | ESE00262 | PN127 | Sausage | France | ✓, c | Illumina | 2,542 | 29,365,678 | 48.77 | 18,328 | ERZ2828520 |
| *P. nalgiovense* | ESE | ESE00263 | PN130 | Sausage | France | ✓, c | Illumina | 663 | 31,809,620 | 48.49 | 179,630 | ERZ2828558 |
| *P. nalgiovense* | ESE | ESE00264 | PN142 | Sausage | Italy | c | Illumina | 616 | 31,779,405 | 48.53 | 236,301 | ERZ2828596 |
| *P. nalgiovense* | ESE | ESE00265 | PN143 | Sausage | France | c | Illumina | 532 | 31,250,068 | 48.55 | 245,520 | ERZ2828633 |
| *P. nalgiovense* | ESE | ESE00267 | PN073 | Sausage | Italy |  | Illumina | 692 | 30,918,143 | 48.53 | 153,654 | ERZ2828670 |
| *P. nalgiovense* | ESE | ESE00268 | PN074 | Sausage | Italy |  | Illumina | 735 | 31,585,821 | 48.4 | 174,484 | ERZ2828711 |
| *P. olsonii* | CBS | CBS193.88 |  | Peatmoss soil under *Diffenbachia* sp. | Denmark | m |  |  |  |  |  |  |
| *P. olsonii* | CBS | CBS266.97 |  | Barley | Denmark | ✓, c | Illumina | 100 | 28,772,805 | 49.96 | 584,105 | ERZ2827456 |
| *P. olsonii* | CBS | CBS626.72 |  | Soil | Russia | ✓, c, m | Illumina | 145 | 29,664,249 | 49.82 | 536,054 | ERZ2827499 |
| *P. olsonii* | CBS | CBS833.88 |  | Cactus pot soil | Denmark | ✓, c, m |  |  |  |  |  |  |
| *P. olsonii* | CBS | DTO032-C5 |  | Air sample in bakery | Netherlands | ✓, c |  |  |  |  |  |  |
| *P. olsonii* | CBS | DTO056-H4 |  | Swab sample, house | Netherlands | m |  |  |  |  |  |  |
| *P. olsonii* | CBS | DTO085-E6 |  | Air sample, veranda | Netherlands | ✓, c |  |  |  |  |  |  |
| *P. olsonii* | CBS | DTO167-B8 |  | Indoor air in poultry house | Poland | ✓, c, m |  |  |  |  |  |  |
| *P. olsonii* | CBS | DTO273-A7 |  | Indoor environment | Germany | ✓, c, m |  |  |  |  |  |  |
| *P. olsonii* | ESE | ESE00149 | PN024 | Dry-cured meat | France |  | Illumina | 184 | 29,814,039 | 49.54 | 617,256 | ERZ2827545 |
| *P. olsonii* | MNHN | LCP04863 | PN0105 | Mummy | France | ✓, c | Illumina | 104 | 29,440,668 | 49.41 | 829,514 | ERZ2827586 |
| *P. olsonii* | MNHN | LCP05357 | PN0109 | Refrigerator | France |  | Illumina | 95 | 29,373,279 | 49.86 | 1,148,184 | ERZ2827627 |
| *P. palitans* | MNHN | LCP01628 |  | Cheese | France | ✓, c |  |  |  |  |  |  |
| *P. palitans* | MNHN | LCP04862 |  | Rice flour | France | ✓, c |  |  |  |  |  |  |
| *P. palitans* | MNHN | LCP05446 |  | Inner fridge wall | France | ✓, c |  |  |  |  |  |  |
| *P. palitans* | MNHN | LCP06125 |  | Food | n.a. | ✓, c |  |  |  |  |  |  |
| *P. palitans* | MNHN | LCP06126 |  | Food | n.a. | ✓, c |  |  |  |  |  |  |
| *P. salamii* | CBS | CBS135391 | PN047 | Salami | Italy | ✓, c , m | Illumina | 511 | 34,385,921 | 47.86 | 367,415 | ERZ2829681 |
| *P. salamii* | CBS | CBS135392 | PN048 | Salami named soppressata | Italy | ✓, c | Illumina | 414 | 36,168,087 | 47.4 | 572,850 | ERZ2829727 |
| *P. salamii* | CBS | CBS135393 |  | Capocollo (2 weeks ageing) | Italy | ✓, c, m | Illumina | 557 | 33,764,403 | 48.39 | 308,640 | ERZ2829774 |
| *P. salamii* | CBS | CBS135394 | PN049 | Capocollo (60 days ageing) | Italy | ✓, c | Illumina | 337 | 34,782,843 | 47.45 | 595,190 | ERZ2829802 |
| *P. salamii* | CBS | CBS135395 |  | Hot salami | Italy | c, m | Illumina | 385 | 31,711,110 | 49.26 | 328,839 | ERZ2829851 |
| *P. salamii* | Giancarlo Perrone | CBS135396 | PN059 | Air-dried ham (Prsut) | Slovenia | ✓, c | Illumina | 260 | 33,123,803 | 48.62 | 639,039 | ERZ2829899 |
| *P. salamii* | CBS | CBS135397 |  | Air-dried ham (Prsut) | Slovenia | c | Illumina | 279 | 33,059,587 | 48.61 | 471,156 | ERZ2829950 |
| *P. salamii* | CBS | CBS135399 |  | Air-dried ham (Prsut) | Slovenia | c, m | Illumina | 382 | 34,529,233 | 47.48 | 432,008 | ERZ2830001 |
| *P. salamii* | CBS | CBS135400 |  | Air-dried ham (Prsut) | Slovenia | c | Illumina | 417 | 34,399,282 | 47.48 | 429,365 | ERZ2830046 |
| *P. salamii* | Giancarlo Perrone | CBS135401 | PN060 | Tea leaves | Uganda | ✓, c, m | Illumina | 1,042 | 34,980,374 | 48.9 | 438,403 | ERZ2830088 |
| *P. salamii* | CBS | CBS135402 |  | Tea leaves | Uganda | ✓, c |  |  |  |  |  |  |
| *P. salamii* | CBS | CBS135403 |  | Sewage plant | Denmark | ✓, c, m | Illumina | 415 | 31,684,316 | 49.27 | 317,974 | ERZ2830133 |
| *P. salamii* | CBS | CBS135407 |  | Sewage plant | Denmark | ✓, c | Illumina | 404 | 34,748,438 | 47.42 | 365,854 | ERZ2830175 |
| *P. salamii* | Federico Laich | CCFL11ab1 |  | Sausage | Argentina | ✓, c |  |  |  |  |  |  |
| *P. salamii* | Federico Laich | CCFL11ab2.1 |  | Sausage | Argentina | ✓, c | Illumina | 261 | 31,552,640 | 48.77 | 599,400 | ERZ2830215 |
| *P. salamii* | Federico Laich | CCFL4 |  | Sausage | Argentina | ✓ |  |  |  |  |  |  |
| *P. salamii* | Federico Laich | CCFL5 |  | Sausage | Argentina | ✓, c | Illumina | 302 | 32,202,828 | 48.69 | 458,588 | ERZ2830260 |
| *P. salamii* | CBS | DTO032-F3 |  | Soil | Australia | c, m | Illumina | 304 | 30,836,126 | 49.41 | 392,125 | ERZ2830308 |
| *P. salamii* | CBS | DTO198-E3 |  | Salami | Italy |  | Illumina | 205 | 30,529,291 | 48.87 | 565,591 | ERZ2830333 |
| *P. salamii* | CBS | DTO225-B7 |  | Sewage plant | Denmark | ✓, c |  |  |  |  |  |  |
| *P. salamii* | CBS | DTO309-F9 |  | Indoor environment of a factory | Germany | ✓, c, m | Illumina | 164 | 31,295,745 | 48.86 | 652,201 | ERZ2830377 |
| *P. salamii* | CBS | DTO334-B6 |  | Fermented sausage | Spain |  | Illumina | 157 | 30,376,733 | 48.87 | 602,877 | ERZ2830420 |
| *P. salamii* | MNHN | LCP06521 | PN001 | Smoked sausage | France | ✓, c, m | Illumina | 410 | 36,245,078 | 47.38 | 559,456 | ERZ2830465 |
| *P. salamii* | MNHN | LCP06522 | PN002 | Smoked sausage | France | ✓, c, m | Illumina | 711 | 36,247,989 | 47.33 | 257,451 | ERZ2832335 |
| *P. salamii* | MNHN | LCP06523 | PN006 | Dry-cured meat | France | c, m | Illumina | 477 | 36,234,877 | 47.38 | 428,106 | ERZ2830552 |
| *P. salamii* | MNHN | LCP06525 | PN007, M1CC1 | Dry-cured meat | France |  | PacBio | 99 | 37,000,735 | 48.75 | 639,522 | ERZ2880307 |

**Table S2. Genbank accessions of the genomes used in the phylogeny, in the detection of horizontal gene transfers and in the detection of selection.** Some of these genomes were produced by the US Department of Energy Joint Genome Institute http://www.jgi.doe.gov/ in collaboration with the user community.

| Species | Strain | GenBank accession | Sequenced and assembled by the JGI | Phylogeny | HGT detection | Selection detection |
| --- | --- | --- | --- | --- | --- | --- |
| *Aspergillus fumigatus* | Af293 | GCA_000002655.1 |  |  | X |  |
| *Aspergillus nidulans* | FGSC A4 | GCA_000011425.1 |  |  | X |  |
| *Aspergillus niger* | NA | GCA_000002855.2 |  |  | X |  |
| *Aspergillus oryzae* | RIB40 | GCA_000184455.3 |  |  | X |  |
| *Monascus purpureus* | YY-1 | GCA_003184285.1 |  |  | X |  |
| *Penicillium antarcticum* | IBT 31811 | GCA_002072345.1 |  | X | X |  |
| *Penicillium arizonense* | CBS 141311 | GCA_001773325.1 |  | X | X |  |
| *Penicillium biforme* | FM169 | GCA_000577785.1 |  |  | X |  |
| *Penicillium bilaiae* | ATCC 20851 | SRX872072, SRX872073 | X | X | X |  |
| *Penicillium brasilianum* | NA | GCA_001048715.1 |  |  | X |  |
| *Penicillium brasilianum* | LaBioMMi 136 | GCA_002016555.1 |  |  | X |  |
| *Penicillium brevicompactum* | AgRF18 | SRX712419, SRX712420, SRX1726660 | X |  | X |  |
| *Penicillium brevicompactum* | 1011305 | PRJNA346833 | X |  | X |  |
| *Penicillium camemberti* | FM013 | GCA_000513335.1 |  |  | X |  |
| *Penicillium canescens* | ATCC 10419 | SRX872091, SRX872092 | X | X | X |  |
| *Penicillium capsulatum* | ATCC 48375 | GCA_000943765.1 |  |  | X |  |
| *Penicillium capsulatum* | LiaoWQ-2011 | GCA_000943775.1 |  |  | X |  |
| *Penicillium carneum* | LCP05629 | ERS1628900 |  |  | X |  |
| *Penicillium carneum* | LCP05634 | GCA_000577495.1 |  |  | X |  |
| *Penicillium caseifulvum* | LCP05630 | SRX9123553 |  |  | X |  |
| *Penicillium cavernicola* | LCP05631 | SRX9123564 |  |  | X |  |
| *Penicillium chrysogenum* | P2niaD18 | GCA_000710275.1 |  |  | X | X |
| *Penicillium chrysogenum* | HKF42 | GCA_002080375.1 |  |  |  | X |
| *Penicillium chrysogenum* | IB 08/921 | GCA_000801355.1 |  |  |  | X |
| *Penicillium chrysogenum* | KF25 | GCA_000816005.1 |  |  |  | X |
| *Penicillium chrysogenum* | NCPC10086 | GCA_000523475.1 |  |  | X | X |
| *Penicillium citrinum* | JCM 22607 | GCA_001950535.1 |  |  | X |  |
| *Penicillium coprophilum* | IBT 31321 | GCA_002072405.1 |  | X | X |  |
| *Penicillium decumbens* | IBT 11843 | GCA_002072245.1 |  |  | X |  |
| *Penicillium digitatum* | Pd1 | GCA_000315645.2 |  |  | X |  |
| *Penicillium digitatum* | PDC 102 | GCA_001307865.1 |  |  | X |  |
| *Penicillium expansum* | MD-8 | GCF_000769745.1 |  |  | X |  |
| *Penicillium expansum* | ATCC 24692 | SRX872070, SRX872071 | X |  | X |  |
| *Penicillium fellutanum* | ATCC 48694 | SRX872095, SRX872096 | X | X | X |  |
| *Penicillium flavigenum* | IBT 14082 | GCA_002072365.1 |  | X | X |  |
| *Penicillium freii* | DAOM 242723 | GCA_001513925.1 |  |  | X |  |
| *Penicillium fuscoglaucum* | FM041 | GCA_000576735.1 |  |  | X |  |
| *Penicillium glabrum* | DAOM 239074 | SRX711997, SRX711998, SRX1726073, SRX1726074 | X |  | X |  |
| *Penicillium griseofulvum* | PG3 | GCA_001561935.1 |  |  | X |  |
| *Penicillium griseofulvum* | MRI314 | GCA_001735785.1 |  | X | X |  |
| *Penicillium italicum* | GL-Gan1 | GCA_002116305.1 |  |  | X |  |
| *Penicillium italicum* | PHI-1 | GCA_000769765.1 |  |  | X |  |
| *Penicillium janthinellum* | ATCC 10455 | SRX875226, SRX875227 |  |  | X |  |
| *Penicillium janthinellum* | NCIM1366 | GCA_002369805.1 |  |  | X |  |
| *Penicillium lanosocoeruleum* | ATCC 48919 | SRX711999, SRX712000, SRX1726071 | X | X | X |  |
| *Penicillium nalgiovense* | FM193 (LCP06099) | GCA_000577395.2 |  | X | X | X |
| *Penicillium nordicum* | UASWS BFE487 | GCA_000733025.2 |  |  | X |  |
| *Penicillium nordicum* | DAOMC 185683 | GCA_001278595.1 |  |  | X |  |
| *Penicillium oxalicum* | HP7-1 | GCA_001723175.2 |  | X | X |  |
| *Penicillium palitans* | LCP04862 | SRX9123535 |  |  | X |  |
| *Penicillium paneum* | FM227 | GCA_000577715.1 |  |  | X |  |
| *Penicillium paxilli* | ATCC 26601 | GCA_000347475.1 |  |  | X |  |
| *Penicillium polonicum* | hy4 | GCA_003344595.1 |  |  | X |  |
| *Penicillium polonicum* | IBT 4502 | GCA_002072265.1 |  | X | X |  |
| *Penicillium psychrosexualis* | CBS128137 | ERS1628901 |  |  | X |  |
| *Penicillium raistrickii* | ATCC 10490 | SRX712423, SRX712424, SRX1726661 | X | X | X |  |
| *Penicillium roqueforti* | FM164 | GCA_000513255.1 |  |  | X |  |
| *Penicillium roqueforti* | CECT 2905 | GCA_001939915.1 |  |  | X |  |
| *Penicillium rubens* | Wisconsin 54-1255 | GCF_000226395.1 |  |  | X |  |
| *Penicillium sclerotiorum* | 113 | GCA_001750025.1 |  |  | X |  |
| *Penicillium solitum* | RS1 | GCA_000952775.2 |  |  | X |  |
| *Penicillium solitum* | IBT 29525 | GCA_002072235.1 |  |  | X |  |
| *Penicillium steckii* | IBT 24891 | GCA_002072375.1 |  | X | X |  |
| *Penicillium subrubescens* | CBS 132785 | GCA_001908125.1 |  | X | X |  |
| *Penicillium swiecickii* | 182 6C1 | SRX3959626 | X | X | X |  |
| *Penicillium thymicola* | DAOMC 180753 | SRX950541 | X |  | X |  |
| *Penicillium verrucosum* | BFE808 | GCA_000970515.2 |  |  | X |  |
| *Penicillium vulpinum* | IBT 29486 | GCA_002072255.1 |  | X | X |  |

**Table S3. Length of the putative horizontally transferred regions (HTR) identified in the *Penicillium nalgiovense* and *P. salamii* strains.** In the “repeat-masked” dataset, all repeated regions are masked, identified or not as belonging to known transposable element (TE) families. In the “TE-masked” dataset, only TE belonging to known families are masked.

| HTR | "repeat-masked" dataset (length in bp) | "TE-masked" dataset (length in bp) | complete dataset (length in bp) |
| --- | --- | --- | --- |
| HTR_01 | 2048 | 8828 | 8894 |
| HTR_02 | 50988 | 87726 | 108222 |
| HTR_03 | 3858 | 7256 | 14080 |
| HTR_04 | 65937 | 68101 | 74415 |
| HTR_05 | 85310 | 89535 | 96877 |
| HTR_06 | 193287 | 438000 | 496686 |
| HTR_07 | 56928 | 67394 | 67892 |
| HTR_08 | 46119 | 115405 | 128220 |
| HTR_09 | 6572 | 6982 | 7224 |
| HTR_10 | 25983 | 63528 | 65006 |
| HTR_11 | 6788 | 7349 | 7634 |
| HTR_12 | 25105 | 53920 | 54288 |
| HTR_13 | 8439 | 9565 | 9612 |
| HTR_14 | 4867 | 7887 | 7969 |
| HTR_15 | 41842 | 48034 | 55620 |
| HTR_16 | 31019 | 33340 | 36641 |
| HTR_17 | 2326 | 27094 | 27325 |
| HTR_18 | 1701 | 12734 | 12912 |
| *HTR_19* | *284* | *12457* | *12635* |
| HTR_20 | 2812 | 7740 | 8172 |
| HTR_21 | 20760 | 30701 | 34516 |
| *HTR_22* | *485* | *10859* | *10859* |
| HTR_23 | 1707 | 8787 | 8787 |
| *HTR_24* | *0* | *11361* | *11361* |
| HTR_25 | 41754 | 42462 | 48359 |
| HTR_26 | 16258 | 24379 | 32858 |

**Table S4. Coverage and identity percentage of each HTR present in each *P. nalgiovense* and *P. salamii* strain.**

| HTR | Assembly | %Id | Coverage |
| --- | --- | --- | --- |
| HTR_02 | Pnal-CBS112438 | 0.984 | 0.795 |
| HTR_02 | Pnal-CBS297-97 | 0.917 | 0.032 |
| HTR_02 | Pnal-CBS318-92 | 0.985 | 0.783 |
| HTR_02 | Pnal-DTO053-A9 | 0.983 | 0.816 |
| HTR_02 | Pnal-DTO204-F1 | 0.954 | 0.038 |
| HTR_02 | Pnal-DTO259-D5 | 0.949 | 0.800 |
| HTR_02 | Pnal-DTO334-B7 | 0.978 | 0.796 |
| HTR_02 | Pnal-DTO334-C1 | 0.980 | 0.807 |
| HTR_02 | Pnal-DTO335-D4 | 0.981 | 0.791 |
| HTR_02 | Pnal-ESE00253 | 0.976 | 0.804 |
| HTR_02 | Pnal-ESE00254 | 0.980 | 0.803 |
| HTR_02 | Pnal-ESE00255 | 0.984 | 0.800 |
| HTR_02 | Pnal-ESE00256 | 0.975 | 0.803 |
| HTR_02 | Pnal-ESE00257 | 0.970 | 0.796 |
| HTR_02 | Pnal-ESE00258 | 0.982 | 0.795 |
| HTR_02 | Pnal-ESE00259 | 0.979 | 0.790 |
| HTR_02 | Pnal-ESE00260 | 0.981 | 0.795 |
| HTR_02 | Pnal-ESE00261 | 0.977 | 0.791 |
| HTR_02 | Pnal-ESE00262 | 0.987 | 0.785 |
| HTR_02 | Pnal-ESE00263 | 0.978 | 0.784 |
| HTR_02 | Pnal-ESE00264 | 0.979 | 0.732 |
| HTR_02 | Pnal-ESE00265 | 0.982 | 0.802 |
| HTR_02 | Pnal-ESE00267 | 0.988 | 0.785 |
| HTR_02 | Pnal-ESE00268 | 0.981 | 0.813 |
| HTR_02 | Pnal-LCP03435 | 0.980 | 0.789 |
| HTR_02 | Pnal-LCP03915 | 0.981 | 0.765 |
| HTR_02 | Pnal-LCP03991 | 0.974 | 0.800 |
| HTR_02 | Pnal-LCP05232 | 0.975 | 0.798 |
| HTR_02 | Psal-CBS135391 | 0.976 | 0.763 |
| HTR_02 | Psal-CBS135392 | 0.973 | 0.754 |
| HTR_02 | Psal-CBS135393 | 0.970 | 0.714 |
| HTR_02 | Psal-CBS135394 | 0.980 | 0.743 |
| HTR_02 | Psal-CBS135395 | 0.971 | 0.013 |
| HTR_02 | Psal-CBS135396 | 0.974 | 0.836 |
| HTR_02 | Psal-CBS135397 | 0.977 | 0.818 |
| HTR_02 | Psal-CBS135399 | 0.980 | 0.756 |
| HTR_02 | Psal-CBS135400 | 0.979 | 0.740 |
| HTR_02 | Psal-CBS135401 | 0.948 | 0.014 |
| HTR_02 | Psal-CBS135403 | 0.987 | 0.028 |
| HTR_02 | Psal-CBS135407 | 0.981 | 0.723 |
| HTR_02 | Psal-CCFL11ab2-1 | 0.984 | 0.812 |
| HTR_02 | Psal-CCFL5 | 0.982 | 0.810 |
| HTR_02 | Psal-DTO198-E3 | 1.000 | 0.727 |
| HTR_02 | Psal-DTO309-F9 | 0.990 | 0.718 |
| HTR_02 | Psal-DTO334-B6 | 0.997 | 0.737 |
| HTR_02 | Psal-LCP06521 | 0.972 | 0.753 |
| HTR_02 | Psal-LCP06522 | 0.976 | 0.757 |
| HTR_02 | Psal-LCP06523 | 0.971 | 0.795 |
| HTR_04 | Pnal-CBS112438 | 0.995 | 0.916 |
| HTR_04 | Pnal-CBS297-97 | 0.899 | 0.033 |
| HTR_04 | Pnal-CBS318-92 | 0.997 | 0.910 |
| HTR_04 | Pnal-DTO053-A9 | 0.995 | 0.914 |
| HTR_04 | Pnal-DTO259-D5 | 0.994 | 0.995 |
| HTR_04 | Pnal-DTO334-B7 | 0.996 | 0.998 |
| HTR_04 | Pnal-DTO334-C1 | 0.996 | 0.997 |
| HTR_04 | Pnal-DTO335-D4 | 0.993 | 1.000 |
| HTR_04 | Pnal-ESE00253 | 0.997 | 0.991 |
| HTR_04 | Pnal-ESE00254 | 0.995 | 1.000 |
| HTR_04 | Pnal-ESE00255 | 0.995 | 0.998 |
| HTR_04 | Pnal-ESE00256 | 0.996 | 0.998 |
| HTR_04 | Pnal-ESE00257 | 0.997 | 0.997 |
| HTR_04 | Pnal-ESE00258 | 0.996 | 1.000 |
| HTR_04 | Pnal-ESE00259 | 0.995 | 1.000 |
| HTR_04 | Pnal-ESE00260 | 0.995 | 0.908 |
| HTR_04 | Pnal-ESE00261 | 0.996 | 0.998 |
| HTR_04 | Pnal-ESE00262 | 0.994 | 0.950 |
| HTR_04 | Pnal-ESE00263 | 0.997 | 0.998 |
| HTR_04 | Pnal-ESE00264 | 0.997 | 0.997 |
| HTR_04 | Pnal-ESE00265 | 0.998 | 1.000 |
| HTR_04 | Pnal-ESE00267 | 0.995 | 0.995 |
| HTR_04 | Pnal-ESE00268 | 0.996 | 0.993 |
| HTR_04 | Pnal-LCP03435 | 0.996 | 0.997 |
| HTR_04 | Pnal-LCP03915 | 0.994 | 1.000 |
| HTR_04 | Pnal-LCP03991 | 0.998 | 0.991 |
| HTR_04 | Pnal-LCP05232 | 0.996 | 0.915 |
| HTR_04 | Psal-CBS135391 | 0.991 | 0.990 |
| HTR_04 | Psal-CBS135392 | 0.990 | 0.882 |
| HTR_04 | Psal-CBS135393 | 0.989 | 0.989 |
| HTR_04 | Psal-CBS135394 | 0.983 | 0.906 |
| HTR_04 | Psal-CBS135395 | 0.943 | 0.319 |
| HTR_04 | Psal-CBS135396 | 0.980 | 0.946 |
| HTR_04 | Psal-CBS135397 | 0.977 | 0.979 |
| HTR_04 | Psal-CBS135399 | 0.944 | 0.268 |
| HTR_04 | Psal-CBS135400 | 0.941 | 0.265 |
| HTR_04 | Psal-CBS135401 | 0.933 | 0.071 |
| HTR_04 | Psal-CBS135403 | 0.944 | 0.304 |
| HTR_04 | Psal-CBS135407 | 0.989 | 1.000 |
| HTR_04 | Psal-CCFL11ab2-1 | 0.922 | 0.269 |
| HTR_04 | Psal-CCFL5 | 0.978 | 0.992 |
| HTR_04 | Psal-DTO032-F3 | 0.919 | 0.102 |
| HTR_04 | Psal-DTO198-E3 | 0.931 | 0.202 |
| HTR_04 | Psal-DTO309-F9 | 0.923 | 0.061 |
| HTR_04 | Psal-DTO334-B6 | 0.921 | 0.186 |
| HTR_04 | Psal-LCP06521 | 0.989 | 1.000 |
| HTR_04 | Psal-LCP06522 | 0.977 | 0.949 |
| HTR_04 | Psal-LCP06523 | 0.987 | 0.902 |
| HTR_05 | Pnal-CBS112438 | 0.959 | 0.903 |
| HTR_05 | Pnal-CBS297-97 | 0.992 | 0.005 |
| HTR_05 | Pnal-CBS318-92 | 0.959 | 0.921 |
| HTR_05 | Pnal-DTO053-A9 | 0.905 | 0.923 |
| HTR_05 | Pnal-DTO259-D5 | 0.955 | 0.862 |
| HTR_05 | Pnal-DTO334-B7 | 0.956 | 0.844 |
| HTR_05 | Pnal-DTO334-C1 | 0.956 | 0.849 |
| HTR_05 | Pnal-DTO335-D4 | 0.958 | 0.911 |
| HTR_05 | Pnal-ESE00253 | 0.958 | 0.911 |
| HTR_05 | Pnal-ESE00254 | 0.899 | 0.866 |
| HTR_05 | Pnal-ESE00255 | 0.932 | 0.857 |
| HTR_05 | Pnal-ESE00256 | 0.904 | 0.925 |
| HTR_05 | Pnal-ESE00257 | 0.899 | 0.885 |
| HTR_05 | Pnal-ESE00258 | 0.898 | 0.857 |
| HTR_05 | Pnal-ESE00259 | 0.957 | 0.942 |
| HTR_05 | Pnal-ESE00260 | 0.904 | 0.928 |
| HTR_05 | Pnal-ESE00261 | 0.956 | 0.842 |
| HTR_05 | Pnal-ESE00262 | 0.996 | 0.773 |
| HTR_05 | Pnal-ESE00263 | 0.955 | 0.871 |
| HTR_05 | Pnal-ESE00264 | 0.900 | 0.886 |
| HTR_05 | Pnal-ESE00265 | 0.900 | 0.875 |
| HTR_05 | Pnal-ESE00267 | 0.959 | 0.905 |
| HTR_05 | Pnal-ESE00268 | 0.946 | 0.869 |
| HTR_05 | Pnal-LCP03435 | 0.903 | 0.896 |
| HTR_05 | Pnal-LCP03915 | 0.906 | 0.942 |
| HTR_05 | Pnal-LCP03991 | 0.900 | 0.869 |
| HTR_05 | Pnal-LCP05232 | 0.939 | 0.929 |
| HTR_05 | Psal-CBS135391 | 0.983 | 0.904 |
| HTR_05 | Psal-CBS135392 | 0.992 | 0.943 |
| HTR_05 | Psal-CBS135393 | 0.985 | 0.881 |
| HTR_05 | Psal-CBS135394 | 0.983 | 0.950 |
| HTR_05 | Psal-CBS135395 | 0.941 | 0.235 |
| HTR_05 | Psal-CBS135396 | 0.915 | 0.964 |
| HTR_05 | Psal-CBS135397 | 0.913 | 0.945 |
| HTR_05 | Psal-CBS135399 | 0.932 | 0.127 |
| HTR_05 | Psal-CBS135400 | 0.935 | 0.126 |
| HTR_05 | Psal-CBS135401 | 0.923 | 0.022 |
| HTR_05 | Psal-CBS135403 | 0.939 | 0.246 |
| HTR_05 | Psal-CBS135407 | 0.989 | 0.919 |
| HTR_05 | Psal-CCFL11ab2-1 | 0.938 | 0.185 |
| HTR_05 | Psal-CCFL5 | 0.909 | 0.917 |
| HTR_05 | Psal-DTO032-F3 | 0.965 | 0.047 |
| HTR_05 | Psal-DTO198-E3 | 0.943 | 0.217 |
| HTR_05 | Psal-DTO309-F9 | 0.911 | 0.047 |
| HTR_05 | Psal-DTO334-B6 | 0.930 | 0.173 |
| HTR_05 | Psal-LCP06521 | 0.985 | 0.894 |
| HTR_05 | Psal-LCP06522 | 0.975 | 0.910 |
| HTR_05 | Psal-LCP06523 | 0.986 | 0.941 |
| HTR_06 | Pnal-CBS112438 | 0.956 | 0.557 |
| HTR_06 | Pnal-CBS297-97 | 0.906 | 0.308 |
| HTR_06 | Pnal-CBS318-92 | 0.944 | 0.594 |
| HTR_06 | Pnal-DTO053-A9 | 0.948 | 0.629 |
| HTR_06 | Pnal-DTO204-F1 | 0.942 | 0.326 |
| HTR_06 | Pnal-DTO259-D5 | 0.957 | 0.623 |
| HTR_06 | Pnal-DTO334-B7 | 0.948 | 0.620 |
| HTR_06 | Pnal-DTO334-C1 | 0.958 | 0.638 |
| HTR_06 | Pnal-DTO335-D4 | 0.938 | 0.632 |
| HTR_06 | Pnal-ESE00253 | 0.949 | 0.599 |
| HTR_06 | Pnal-ESE00254 | 0.958 | 0.627 |
| HTR_06 | Pnal-ESE00255 | 0.948 | 0.645 |
| HTR_06 | Pnal-ESE00256 | 0.958 | 0.610 |
| HTR_06 | Pnal-ESE00257 | 0.933 | 0.626 |
| HTR_06 | Pnal-ESE00258 | 0.947 | 0.624 |
| HTR_06 | Pnal-ESE00259 | 0.960 | 0.624 |
| HTR_06 | Pnal-ESE00260 | 0.947 | 0.627 |
| HTR_06 | Pnal-ESE00261 | 0.949 | 0.584 |
| HTR_06 | Pnal-ESE00262 | 0.968 | 0.572 |
| HTR_06 | Pnal-ESE00263 | 0.958 | 0.605 |
| HTR_06 | Pnal-ESE00264 | 0.948 | 0.627 |
| HTR_06 | Pnal-ESE00265 | 0.946 | 0.655 |
| HTR_06 | Pnal-ESE00267 | 0.950 | 0.608 |
| HTR_06 | Pnal-ESE00268 | 0.957 | 0.617 |
| HTR_06 | Pnal-LCP03435 | 0.936 | 0.634 |
| HTR_06 | Pnal-LCP03915 | 0.934 | 0.659 |
| HTR_06 | Pnal-LCP03991 | 0.948 | 0.641 |
| HTR_06 | Pnal-LCP05232 | 0.938 | 0.615 |
| HTR_06 | Psal-CBS135391 | 0.926 | 0.083 |
| HTR_06 | Psal-CBS135392 | 0.970 | 0.774 |
| HTR_06 | Psal-CBS135393 | 0.927 | 0.078 |
| HTR_06 | Psal-CBS135394 | 0.926 | 0.084 |
| HTR_06 | Psal-CBS135395 | 0.934 | 0.015 |
| HTR_06 | Psal-CBS135396 | 0.914 | 0.265 |
| HTR_06 | Psal-CBS135397 | 0.916 | 0.261 |
| HTR_06 | Psal-CBS135399 | 0.976 | 0.795 |
| HTR_06 | Psal-CBS135400 | 0.976 | 0.767 |
| HTR_06 | Psal-CBS135401 | 0.927 | 0.005 |
| HTR_06 | Psal-CBS135403 | 0.935 | 0.015 |
| HTR_06 | Psal-CBS135407 | 0.928 | 0.087 |
| HTR_06 | Psal-CCFL11ab2-1 | 0.889 | 0.186 |
| HTR_06 | Psal-CCFL5 | 0.939 | 0.221 |
| HTR_06 | Psal-DTO032-F3 | 0.926 | 0.021 |
| HTR_06 | Psal-DTO198-E3 | 0.958 | 0.190 |
| HTR_06 | Psal-DTO309-F9 | 0.928 | 0.140 |
| HTR_06 | Psal-DTO334-B6 | 0.953 | 0.109 |
| HTR_06 | Psal-LCP06521 | 0.971 | 0.776 |
| HTR_06 | Psal-LCP06522 | 0.972 | 0.723 |
| HTR_06 | Psal-LCP06523 | 0.969 | 0.757 |
| HTR_07 | Pnal-CBS112438 | 0.992 | 0.932 |
| HTR_07 | Pnal-CBS297-97 | 0.930 | 0.037 |
| HTR_07 | Pnal-CBS318-92 | 0.992 | 0.889 |
| HTR_07 | Pnal-DTO053-A9 | 0.990 | 0.896 |
| HTR_07 | Pnal-DTO204-F1 | 0.923 | 0.039 |
| HTR_07 | Pnal-DTO259-D5 | 0.992 | 0.909 |
| HTR_07 | Pnal-DTO334-B7 | 0.991 | 0.896 |
| HTR_07 | Pnal-DTO334-C1 | 0.989 | 0.896 |
| HTR_07 | Pnal-DTO335-D4 | 0.993 | 0.937 |
| HTR_07 | Pnal-ESE00253 | 0.993 | 0.962 |
| HTR_07 | Pnal-ESE00254 | 0.990 | 0.906 |
| HTR_07 | Pnal-ESE00255 | 0.992 | 0.933 |
| HTR_07 | Pnal-ESE00256 | 0.993 | 0.925 |
| HTR_07 | Pnal-ESE00257 | 0.989 | 0.863 |
| HTR_07 | Pnal-ESE00258 | 0.990 | 0.895 |
| HTR_07 | Pnal-ESE00259 | 0.990 | 0.896 |
| HTR_07 | Pnal-ESE00260 | 0.990 | 0.955 |
| HTR_07 | Pnal-ESE00261 | 0.989 | 0.864 |
| HTR_07 | Pnal-ESE00262 | 0.991 | 0.779 |
| HTR_07 | Pnal-ESE00263 | 0.990 | 0.942 |
| HTR_07 | Pnal-ESE00264 | 0.991 | 0.863 |
| HTR_07 | Pnal-ESE00265 | 0.989 | 0.864 |
| HTR_07 | Pnal-ESE00267 | 0.993 | 0.957 |
| HTR_07 | Pnal-ESE00268 | 0.991 | 0.939 |
| HTR_07 | Pnal-LCP03435 | 0.990 | 0.864 |
| HTR_07 | Pnal-LCP03915 | 0.992 | 0.907 |
| HTR_07 | Pnal-LCP03991 | 0.991 | 0.896 |
| HTR_07 | Pnal-LCP05232 | 0.989 | 0.891 |
| HTR_07 | Psal-CBS135391 | 0.991 | 0.947 |
| HTR_07 | Psal-CBS135392 | 0.985 | 0.947 |
| HTR_07 | Psal-CBS135393 | 0.988 | 0.919 |
| HTR_07 | Psal-CBS135394 | 0.988 | 0.947 |
| HTR_07 | Psal-CBS135395 | 0.849 | 0.762 |
| HTR_07 | Psal-CBS135396 | 0.975 | 0.962 |
| HTR_07 | Psal-CBS135397 | 0.974 | 0.962 |
| HTR_07 | Psal-CBS135399 | 0.933 | 0.135 |
| HTR_07 | Psal-CBS135400 | 0.933 | 0.135 |
| HTR_07 | Psal-CBS135401 | 0.895 | 0.144 |
| HTR_07 | Psal-CBS135403 | 0.849 | 0.758 |
| HTR_07 | Psal-CBS135407 | 0.989 | 0.938 |
| HTR_07 | Psal-CCFL11ab2-1 | 0.917 | 0.154 |
| HTR_07 | Psal-CCFL5 | 0.977 | 0.965 |
| HTR_07 | Psal-DTO198-E3 | 0.923 | 0.197 |
| HTR_07 | Psal-DTO309-F9 | 0.917 | 0.066 |
| HTR_07 | Psal-DTO334-B6 | 0.922 | 0.141 |
| HTR_07 | Psal-LCP06521 | 0.985 | 0.940 |
| HTR_07 | Psal-LCP06522 | 0.976 | 0.935 |
| HTR_07 | Psal-LCP06523 | 0.984 | 0.921 |
| HTR_08 | Pnal-CBS112438 | 0.960 | 0.503 |
| HTR_08 | Pnal-CBS297-97 | 0.940 | 0.424 |
| HTR_08 | Pnal-CBS318-92 | 0.952 | 0.496 |
| HTR_08 | Pnal-DTO053-A9 | 0.953 | 0.569 |
| HTR_08 | Pnal-DTO204-F1 | 0.943 | 0.259 |
| HTR_08 | Pnal-DTO259-D5 | 0.945 | 0.566 |
| HTR_08 | Pnal-DTO334-B7 | 0.951 | 0.597 |
| HTR_08 | Pnal-DTO334-C1 | 0.952 | 0.628 |
| HTR_08 | Pnal-DTO335-D4 | 0.950 | 0.589 |
| HTR_08 | Pnal-ESE00253 | 0.953 | 0.632 |
| HTR_08 | Pnal-ESE00254 | 0.947 | 0.636 |
| HTR_08 | Pnal-ESE00255 | 0.953 | 0.561 |
| HTR_08 | Pnal-ESE00256 | 0.946 | 0.653 |
| HTR_08 | Pnal-ESE00257 | 0.955 | 0.592 |
| HTR_08 | Pnal-ESE00258 | 0.947 | 0.600 |
| HTR_08 | Pnal-ESE00259 | 0.954 | 0.594 |
| HTR_08 | Pnal-ESE00260 | 0.955 | 0.634 |
| HTR_08 | Pnal-ESE00261 | 0.956 | 0.572 |
| HTR_08 | Pnal-ESE00262 | 0.956 | 0.530 |
| HTR_08 | Pnal-ESE00263 | 0.913 | 0.518 |
| HTR_08 | Pnal-ESE00264 | 0.950 | 0.603 |
| HTR_08 | Pnal-ESE00265 | 0.956 | 0.577 |
| HTR_08 | Pnal-ESE00267 | 0.948 | 0.525 |
| HTR_08 | Pnal-ESE00268 | 0.951 | 0.554 |
| HTR_08 | Pnal-LCP03435 | 0.952 | 0.618 |
| HTR_08 | Pnal-LCP03915 | 0.953 | 0.571 |
| HTR_08 | Pnal-LCP03991 | 0.955 | 0.581 |
| HTR_08 | Pnal-LCP05232 | 0.957 | 0.612 |
| HTR_08 | Psal-CBS135391 | 0.924 | 0.087 |
| HTR_08 | Psal-CBS135392 | 0.985 | 0.792 |
| HTR_08 | Psal-CBS135393 | 0.926 | 0.087 |
| HTR_08 | Psal-CBS135394 | 0.925 | 0.087 |
| HTR_08 | Psal-CBS135395 | 0.944 | 0.009 |
| HTR_08 | Psal-CBS135396 | 0.991 | 0.901 |
| HTR_08 | Psal-CBS135397 | 0.988 | 0.882 |
| HTR_08 | Psal-CBS135399 | 0.958 | 0.034 |
| HTR_08 | Psal-CBS135400 | 0.867 | 0.032 |
| HTR_08 | Psal-CBS135401 | 0.872 | 0.008 |
| HTR_08 | Psal-CBS135403 | 0.945 | 0.009 |
| HTR_08 | Psal-CBS135407 | 0.916 | 0.096 |
| HTR_08 | Psal-CCFL11ab2-1 | 0.941 | 0.260 |
| HTR_08 | Psal-CCFL5 | 0.989 | 0.862 |
| HTR_08 | Psal-DTO032-F3 | 0.939 | 0.021 |
| HTR_08 | Psal-DTO198-E3 | 0.941 | 0.248 |
| HTR_08 | Psal-DTO309-F9 | 0.952 | 0.199 |
| HTR_08 | Psal-DTO334-B6 | 0.940 | 0.221 |
| HTR_08 | Psal-LCP06521 | 0.985 | 0.819 |
| HTR_08 | Psal-LCP06522 | 0.985 | 0.729 |
| HTR_08 | Psal-LCP06523 | 0.984 | 0.737 |
| HTR_10 | Pnal-CBS112438 | 0.945 | 0.654 |
| HTR_10 | Pnal-CBS297-97 | 0.932 | 0.229 |
| HTR_10 | Pnal-CBS318-92 | 0.947 | 0.661 |
| HTR_10 | Pnal-DTO053-A9 | 0.936 | 0.613 |
| HTR_10 | Pnal-DTO204-F1 | 0.966 | 0.583 |
| HTR_10 | Pnal-DTO259-D5 | 0.956 | 0.597 |
| HTR_10 | Pnal-DTO334-B7 | 0.948 | 0.584 |
| HTR_10 | Pnal-DTO334-C1 | 0.947 | 0.613 |
| HTR_10 | Pnal-DTO335-D4 | 0.941 | 0.704 |
| HTR_10 | Pnal-ESE00253 | 0.938 | 0.641 |
| HTR_10 | Pnal-ESE00254 | 0.945 | 0.671 |
| HTR_10 | Pnal-ESE00255 | 0.941 | 0.648 |
| HTR_10 | Pnal-ESE00256 | 0.949 | 0.543 |
| HTR_10 | Pnal-ESE00257 | 0.942 | 0.730 |
| HTR_10 | Pnal-ESE00258 | 0.949 | 0.678 |
| HTR_10 | Pnal-ESE00259 | 0.943 | 0.670 |
| HTR_10 | Pnal-ESE00260 | 0.948 | 0.587 |
| HTR_10 | Pnal-ESE00261 | 0.945 | 0.584 |
| HTR_10 | Pnal-ESE00262 | 0.957 | 0.684 |
| HTR_10 | Pnal-ESE00263 | 0.960 | 0.687 |
| HTR_10 | Pnal-ESE00264 | 0.942 | 0.736 |
| HTR_10 | Pnal-ESE00265 | 0.942 | 0.639 |
| HTR_10 | Pnal-ESE00267 | 0.950 | 0.549 |
| HTR_10 | Pnal-ESE00268 | 0.961 | 0.677 |
| HTR_10 | Pnal-LCP03435 | 0.942 | 0.663 |
| HTR_10 | Pnal-LCP03915 | 0.938 | 0.666 |
| HTR_10 | Pnal-LCP03991 | 0.941 | 0.740 |
| HTR_10 | Pnal-LCP05232 | 0.946 | 0.654 |
| HTR_10 | Psal-CBS135391 | 0.913 | 0.019 |
| HTR_10 | Psal-CBS135392 | 0.986 | 0.766 |
| HTR_10 | Psal-CBS135393 | 0.913 | 0.019 |
| HTR_10 | Psal-CBS135396 | 0.935 | 0.402 |
| HTR_10 | Psal-CBS135397 | 0.932 | 0.382 |
| HTR_10 | Psal-CBS135399 | 0.999 | 0.912 |
| HTR_10 | Psal-CBS135400 | 0.996 | 0.898 |
| HTR_10 | Psal-CBS135407 | 0.913 | 0.019 |
| HTR_10 | Psal-CCFL11ab2-1 | 0.939 | 0.141 |
| HTR_10 | Psal-CCFL5 | 0.932 | 0.270 |
| HTR_10 | Psal-DTO198-E3 | 0.946 | 0.155 |
| HTR_10 | Psal-DTO309-F9 | 0.942 | 0.171 |
| HTR_10 | Psal-DTO334-B6 | 0.950 | 0.136 |
| HTR_10 | Psal-LCP06521 | 0.990 | 0.863 |
| HTR_10 | Psal-LCP06522 | 0.988 | 0.778 |
| HTR_10 | Psal-LCP06523 | 0.984 | 0.738 |
| HTR_12 | Pnal-CBS112438 | 0.968 | 0.903 |
| HTR_12 | Pnal-CBS297-97 | 0.891 | 0.137 |
| HTR_12 | Pnal-CBS318-92 | 0.969 | 0.919 |
| HTR_12 | Pnal-DTO053-A9 | 0.973 | 0.880 |
| HTR_12 | Pnal-DTO204-F1 | 0.916 | 0.067 |
| HTR_12 | Pnal-DTO259-D5 | 0.974 | 0.733 |
| HTR_12 | Pnal-DTO334-B7 | 0.970 | 0.835 |
| HTR_12 | Pnal-DTO334-C1 | 0.971 | 0.894 |
| HTR_12 | Pnal-DTO335-D4 | 0.974 | 0.831 |
| HTR_12 | Pnal-ESE00253 | 0.971 | 0.925 |
| HTR_12 | Pnal-ESE00254 | 0.966 | 0.870 |
| HTR_12 | Pnal-ESE00255 | 0.972 | 0.829 |
| HTR_12 | Pnal-ESE00256 | 0.972 | 0.819 |
| HTR_12 | Pnal-ESE00257 | 0.965 | 0.934 |
| HTR_12 | Pnal-ESE00258 | 0.969 | 0.823 |
| HTR_12 | Pnal-ESE00259 | 0.964 | 0.844 |
| HTR_12 | Pnal-ESE00260 | 0.969 | 0.925 |
| HTR_12 | Pnal-ESE00261 | 0.971 | 0.806 |
| HTR_12 | Pnal-ESE00262 | 0.976 | 0.642 |
| HTR_12 | Pnal-ESE00263 | 0.962 | 0.674 |
| HTR_12 | Pnal-ESE00264 | 0.975 | 0.863 |
| HTR_12 | Pnal-ESE00265 | 0.970 | 0.865 |
| HTR_12 | Pnal-ESE00267 | 0.974 | 0.811 |
| HTR_12 | Pnal-ESE00268 | 0.973 | 0.757 |
| HTR_12 | Pnal-LCP03435 | 0.969 | 0.872 |
| HTR_12 | Pnal-LCP03915 | 0.975 | 0.897 |
| HTR_12 | Pnal-LCP03991 | 0.974 | 0.909 |
| HTR_12 | Pnal-LCP05232 | 0.972 | 0.900 |
| HTR_12 | Psal-CBS135391 | 0.940 | 0.087 |
| HTR_12 | Psal-CBS135392 | 0.993 | 0.939 |
| HTR_12 | Psal-CBS135393 | 0.940 | 0.087 |
| HTR_12 | Psal-CBS135394 | 0.940 | 0.087 |
| HTR_12 | Psal-CBS135395 | 0.846 | 0.046 |
| HTR_12 | Psal-CBS135396 | 0.994 | 0.926 |
| HTR_12 | Psal-CBS135397 | 0.996 | 0.889 |
| HTR_12 | Psal-CBS135399 | 0.941 | 0.057 |
| HTR_12 | Psal-CBS135400 | 0.941 | 0.057 |
| HTR_12 | Psal-CBS135401 | 0.918 | 0.019 |
| HTR_12 | Psal-CBS135403 | 0.846 | 0.046 |
| HTR_12 | Psal-CBS135407 | 0.940 | 0.087 |
| HTR_12 | Psal-CCFL11ab2-1 | 0.949 | 0.146 |
| HTR_12 | Psal-CCFL5 | 0.998 | 0.960 |
| HTR_12 | Psal-DTO198-E3 | 0.944 | 0.091 |
| HTR_12 | Psal-DTO309-F9 | 0.940 | 0.084 |
| HTR_12 | Psal-DTO334-B6 | 0.909 | 0.170 |
| HTR_12 | Psal-LCP06521 | 0.993 | 0.948 |
| HTR_12 | Psal-LCP06522 | 0.995 | 0.862 |
| HTR_12 | Psal-LCP06523 | 0.993 | 0.909 |

**Table S5. Number of strains of *Penicillium nalgiovense* and *P. salamii* harboring each cluster of orthologous groups (COG) functions for the genes present in HTRs, and distribution of the COG functions of the genes present on the HTR 2, 4, 5, 6, 7, 15 and 25 in *P. biforme*, *P. camemberti* and *P. caseifulvum*.** The highlighted functions (in bold) are the most prevalent ones in *P. nalgiovense* and *P. salamii* strains. Note that many genes have an unknown function.

| COG functional category | Functional description | All strains  *P. nalgiovense* | All strains *P. salamii* | *P. biforme* FM169 | *P. biforme* ESE00211 | *P. camemberti* FM013 | *P. caseifulvum* LCP05630 |
| --- | --- | --- | --- | --- | --- | --- | --- |
| A | **RNA processing and modification** | **28** | **8** |  |  |  |  |
| G | **Carbohydrate transport and metabolism** | **27** | **19** | 1 | 5 | 1 | 1 |
| H | Coenzyme transport and metabolism | 0 | 1 |  |  |  |  |
| I | **Lipid transport and metabolism** | **26** | **0** |  | 1 | 1 | 1 |
| J | Translation, ribosomal structure and biogenesis | 2 | 0 |  |  |  |  |
| K | **Transcription** | **14** | **9** |  |  |  |  |
| M | **Cell wall/membrane/envelope biogenesis** | **21** | **1** |  |  |  |  |
| O | **Post-translational modification, protein turnover, and chaperones** | **28** | **14** | 2 | 2 | 2 |  |
| P | Inorganic ion transport and metabolism | 1 | 7 |  | 1 |  | 1 |
| Q | **Secondary metabolites biosynthesis, transport, and catabolism** | **25** | **1** |  | 1 | 1 | 1 |
| S | Function unknown | 28 | 20 | 3 | 9 | 3 | 2 |
| U | Intracellular trafficking, secretion, and vesicular transport | 0 | 2 |  |  | 1 | 1 |
| Z | **Cytoskeleton** | **26** | **7** | 6 | 2 |  |  |
|  | Total number of genes in HTR |  |  | 50 | 57 | 42 | 46 |

**Table S6. Detection of selection among 7,155 genes in *Penicillium nalgiovense* and 7,010 genes in *P. salamii*, associated with their cluster of orthologous group (COG) functional categories**. As each gene could have been assigned to more than one category or to none, we consider here the number of occurrences of each category (the sum can therefore be different from the number of genes). An excess of representation of negative or positive selection between one category and the rest of the genes is identified by a * when p<0.05, ** when p<0.01, *** when p<0.001 (Fisher’s exact test).

| COG functional category | Functional description | Number of functions displayed by the genes under negative selection | | | Number of functions displayed by the genes under positive selection | | |
| --- | --- | --- | --- | --- | --- | --- | --- |
|  |  | Both species | *P. nalgiovense* | *P. salamii* | Both species | *P. nalgiovense* | *P. salamii* |
| A | RNA processing and modification | 4 | 42 | 18 |  | 1 | 19 |
| B | Chromatin structure and dynamics |  | 17 | 3 |  |  | 13 |
| C | Energy production and conversion | 5 | 57 | 26* |  | 1 | 19 |
| D | Cell cycle control, cell division, chromosome partitioning |  | 29 | 5 | 1 |  | 9 |
| E | Amino acid transport and metabolism | 3 | 83* | 14 |  |  | 26 |
| F | Nucleotide transport and metabolism | 3 | 8 | 4 |  |  | 3 |
| G | Carbohydrate transport and metabolism | 3 | 120* | 35 |  | 1 | 54 |
| H | Coenzyme transport and metabolism |  | 29 | 5 |  |  | 5 |
| I | Lipid transport and metabolism | 3 | 54 | 16 |  |  | 22 |
| J | Translation, ribosomal structure and biogenesis | 5 | 70 | 16 |  | 1 | 25 |
| K | Transcription | 5 | 74 | 25 |  | 1 | 69*** |
| L | Replication, recombination and repair | 2 | 47 | 8 |  |  | 22 |
| M | Cell wall/membrane/envelope biogenesis | 1 | 13 | 6 |  |  | 11 |
| N | Cell motility |  | 1 |  |  |  | 1 |
| O | Post-translational modification, protein turnover, and chaperones | 3 | 70 | 23 |  | 1 | 58* |
| P | Inorganic ion transport and metabolism |  | 35 | 14 |  | 2 | 20 |
| Q | Secondary metabolites biosynthesis, transport, and catabolism | 5 | 60 | 24* |  | 4* | 23 |
| S | Function unknown | 18 | 393 | 76 | 1 | 5 | 216 |
| T | Signal transduction mechanisms | 5 | 71 | 18 | 1 | 1 | 42* |
| U | Intracellular trafficking, secretion, and vesicular transport | 2 | 72 | 24 |  |  | 31 |
| V | Defense mechanisms |  | 10 | 2 |  |  | 8* |
| Y | Nuclear structure |  | 6 | 1 |  |  | 3 |
| Z | Cytoskeleton | 3 | 19 | 6 |  | 1 | 5 |

**Table S7. Detection of twelve mycotoxins and extrolites potentially produced by *Penicillium* species among dry-cured meat *P. nalgiovense* and *P. salamii* and their respective sister species *P. chrysogenum* and *P. olsonii*.** All quantifications for ermefortins A & B, (iso)-fumigaclavin A, mycophenolic acid, ochratoxin A, penicillic acid, penitrem A, sterigmatocystin were below detection limit (<DL; not shown). All results are expressed in ng of mycotoxin per g of fungal dry weight.

|  | andrastin A | | penicillin G | | meleagrin | | roquefortin C | |
| --- | --- | --- | --- | --- | --- | --- | --- | --- |
| Species and strain | Mean | Standard deviation | Mean | Standard deviation | Mean | Standard deviation | Mean | Standard deviation |
| *P. chrysogenum* ESE00097 | <DL |  | <DL |  | <DL |  | 790.94 | 163.14 |
| *P. chrysogenum* ESE00106 | <DL |  | <DL |  | <DL |  | 286.04 | 28.13 |
| *P. chrysogenum* ESE00128 | <DL |  | <DL |  | <DL |  | 282.49 | 81.81 |
| *P. chrysogenum* ESE00131 | <DL |  | <DL |  | <DL |  | 121.54 | 46.26 |
| *P. chrysogenum* ESE00135 | <DL |  | <DL |  | <DL |  | 178.80 | 4.63 |
| *P. chrysogenum* ESE00313 | 639943.99 | 38225.74 | 41753.42 | 2432.92 | 996491.90 | 210929.16 | 48400.27 | 13779.99 |
| *P. chrysogenum* ESE00352 | 496241.47 | 115257.94 | 59229.37 | 14047.66 | 2102233.91 | 247062.20 | 61443.01 | 31547.42 |
| *P. chrysogenum* ESE00361 | 123207.95 | 158358.90 | 42299.58 | 10316.62 | 1024342.88 | 1083701.67 | 33346.53 | 37567.59 |
| *P. chrysogenum* ESE00434 | 603935.02 | 183051.35 | 59229.37 | 14047.66 | 2934637.82 | 1894519.24 | 75865.96 | 59018.28 |
| *P. chrysogenum* LCP00456 | 61532.92 | 13010.60 | <DL |  | <DL |  | 7636.07 | 857.08 |
| *P. chrysogenum* LCP00487 | <DL |  | <DL |  | <DL |  | 16474.74 | 25932.83 |
| *P. chrysogenum* LCP02616 | 317515.78 | 123252.84 | 44587.43 | 4904.40 | 885505.65 | 471765.36 | 33696.99 | 19466.95 |
| *P. chrysogenum* LCP04047 | 520129.29 | 69586.07 | 41272.57 | 2870.38 | 2893607.18 | 276002.22 | 27027.34 | 12477.69 |
| *P. chrysogenum* LCP04082 | 450254.73 | 153940.47 | 41590.81 | 6257.85 | 959675.69 | 661501.77 | 55783.00 | 40370.30 |
| *P. nalgiovense* CBS297.97 | <DL |  | <DL |  | <DL |  | <DL |  |
| *P. nalgiovense* CBS318.92 | <DL |  | <DL |  | <DL |  | <DL |  |
| *P. nalgiovense* DTO053-A9 | <DL |  | <DL |  | <DL |  | <DL |  |
| *P. nalgiovense* DTO204-F1 | <DL |  | <DL |  | <DL |  | <DL |  |
| *P. nalgiovense* DTO259-D5 | <DL |  | <DL |  | <DL |  | <DL |  |
| *P. nalgiovense* DTO334-B7 | <DL |  | <DL |  | <DL |  | <DL |  |
| *P. nalgiovense* DTO334-B8 | <DL |  | <DL |  | <DL |  | <DL |  |
| *P. nalgiovense* DTO335-D4 | <DL |  | <DL |  | <DL |  | <DL |  |
| *P. nalgiovense* LCP05232 | <DL |  | <DL |  | <DL |  | <DL |  |
| *P. olsonii* CBS193.88 | <DL |  | <DL |  | <DL |  | <DL |  |
| *P. olsonii* CBS626.72 | <DL |  | <DL |  | <DL |  | <DL |  |
| *P. olsonii* CBS833.88 | <DL |  | <DL |  | <DL |  | <DL |  |
| *P. olsonii* DTO056-H4 | <DL |  | <DL |  | <DL |  | <DL |  |
| *P. olsonii* DTO167-B8 | <DL |  | <DL |  | <DL |  | <DL |  |
| *P. olsonii* DTO273-A7 | <DL |  | <DL |  | <DL |  | <DL |  |
| *P. salamii* CBS135391 | <DL |  | <DL |  | <DL |  | <DL |  |
| *P. salamii* CBS135393 | <DL |  | <DL |  | <DL |  | <DL |  |
| *P. salamii* CBS135395 | <DL |  | <DL |  | <DL |  | <DL |  |
| *P. salamii* CBS135399 | <DL |  | <DL |  | <DL |  | <DL |  |
| *P. salamii* CBS135401 | <DL |  | <DL |  | <DL |  | <DL |  |
| *P. salamii* CBS135403 | <DL |  | <DL |  | <DL |  | <DL |  |
| *P. salamii* DTO032-F3 | <DL |  | <DL |  | <DL |  | <DL |  |
| *P. salamii* DTO309-F9 | <DL |  | <DL |  | <DL |  | <DL |  |
| *P. salamii* LCP06521 | <DL |  | <DL |  | <DL |  | <DL |  |
| *P. salamii* LCP06522 | <DL |  | <DL |  | <DL |  | <DL |  |
| *P. salamii* LCP06523 | <DL |  | <DL |  | <DL |  | <DL |  |

**Table S8. Method performance characteristics for metabolite quantification from fungal cultures grown on YES medium.** RT: Retention Time; R^2^: correlation coefficient; ESI: Electrospray Ionization; N/A: not applicable.

| Compound | Formula | RT (min) | Quantifier Ion (Q1) (m/z) | Qualifier Ion (Q2) (m/z) | Linear range (ng.µl^-1^) | R^2^ | ESI |
| --- | --- | --- | --- | --- | --- | --- | --- |
| andrastin A | C_28_H_38_O_7_ | 15.8 | 485.2541 | N/A | 1-5000 | 0.960 | - |
| ermefortin A | C_17_H_22_O_5_ | 12.5 | 307.1561 | 329.1358 | 1-5000 | 0.997 | + |
| ermefortin B | C_15_H_20_O_3_ | 10.2 | 249.1484 | 271.1252 | 10-5000 | 0.999 | + |
| (iso)-fumigaclavin A | C_18_H_22_N_2_O_2_ | 1.9 | 299.1751 | 322.1573 | 10-10000 | 0.994 | + |
| meleagrin | C_23_H_23_N_5_O_4_ | 10.4 | 434.1823 | 456.1642 | 1-10000 | 0.996 | + |
| mycophenolic acid | C_17_H_20_O_6_ | 13.3 | 321.1334 | 303.1231 | 1-5000 | 0.999 | + |
| ochratoxin A | C_20_H_18_ClNO_6_ | 14.7 | 404.0895 | 426.0715 | 50-5000 | 0.997 | + |
| penitrem A | C_37_H_44_ClNO_6_ | 18.5 | 632.2781 | N/A | 10-5000 | 0.999 | - |
| penicillin G | C_16_H_18_N_2_O_4_S | 9.6 | 335.1080 | 357.0882 | 1-10000 | 0.989 | + |
| penicillic acid | C_8_H_10_O_6_ | 1.7 | 171.0646 | 193.0466 | 1-10000 | 0.993 | + |
| roquefortin C | C_22_H_23_N_5_O_2_ | 11.4 | 390.1928 | N/A | 1-2500 | 0.971 | + |
| sterigmatocystin | C_18_H_12_O_6_ | 15.3 | 325.0707 | 347.0528 | 10-10000 | 0.999 | + |
